# Post-transcriptional regulation by copper with a new upstream Open Reading Frame

**DOI:** 10.1101/2022.03.18.484875

**Authors:** Gauthier Roy, Rudy Antoine, Annie Schwartz, Stéphanie Slupek, Alex Rivera-Millot, Marc Boudvillain, Françoise Jacob-Dubuisson

**Affiliations:** Univ. Lille, Inserm, CNRS, CHU Lille, Institut Pasteur de Lille, U1019-UMR9017-CIIL-Center for Infection and Immunity of Lille, F-59000, Lille, France; Centre de Biophysique moléculaire, CNRS UPR4301, F-45071 Orléans cedex 2, France; affiliated with Université d’Orléans, France

**Keywords:** copper homeostasis, post-transcriptional regulation, upstream ORF

## Abstract

Copper is essential to most living beings but also toxic. Bacteria have thus developed homeostatic mechanisms to tightly control its intracellular concentration. The 3-gene operon *bp2923-bfrG-bp2921* is down-regulated by copper and notably encodes a TonB-dependent transporter in *Bordetella pertussis*. We show that the protein encoded by *bp2923*, which is a member of the DUF2946 family, represents a new type of <u>u</u>pstream <u>O</u>pen <u>R</u>eading <u>F</u>rame (uORF) involved in post-transcriptional regulation of the downstream genes. In the absence of copper, the entire operon is transcribed and translated. Perception of copper by the nascent *bp2923*-coded protein via its conserved CXXC motif triggers Rho-dependent transcription termination between the first and second genes by relieving translation arrest on a conserved C-terminal RAPP motif. Homologues of *bp2923* are widespread in bacterial genomes, where they head operons predicted to participate in copper homeostasis. This work has unveiled an original mode of genetic regulation by a transition metal and identified a regulatory function for a member of an uncharacterized family of bacterial proteins that we have named CruR, for copper-<u>r</u>esponsive <u>u</u>pstream <u>r</u>egulator.

## Introduction

Bacteria have evolved complex mechanisms to respond to changes of their environment, and notably to strictly regulate the availability of necessary but harmful transition metals. Copper is such a metal both essential and harmful to living beings (Solioz, 2018). Its properties as a redox cycling metal have been put to use in electron transfer chains as a cofactor of heme-copper oxidases for aerobic respiration, photosynthesis and denitrification (Andrei et al., 2020). It is also involved in various hydrolytic and oxidoreduction reactions catalysed by metabolic enzymes, and in the protection against reactive oxygen species. Its high affinity for organic molecules makes copper very toxic, notably because it destroys iron-sulphur clusters during or after biogenesis and induces oxidative stress (Macomber and Imlay, 2009). For its capacity to kill microorganisms, copper is notably used in healthcare settings and agriculture and has become a common pollutant (Faundez et al., 2004, Lemire et al., 2013, Rehman et al., 2019, Rensing et al., 2018, Vincent et al., 2018). Eukaryotic phagocytes in natural milieus (e.g., amoeba) and at the host-pathogen interface notably employ copper to kill microorganisms (Hao et al., 2016, Sheldon and Skaar, 2019). Life with copper has therefore led bacteria to develop homeostatic mechanisms that strictly control its intracellular concentration (Arguello et al., 2013, Andrei et al., 2020, Chandrangsu et al., 2017). Defence systems against copper include export of Cu^1+^ from the cytoplasm or its passivation by sequestration, the detoxification of Cu^1+^ into Cu^2+^ in the periplasm and its extrusion to the extracellular medium (Solioz, 2018).

Bacteria also need to acquire copper from their environment (Stewart et al., 2019), and the few described copper uptake systems are dedicated to the assembly of specific cuproproteins (Ekici et al., 2014, Lee et al., 1989). The expression of homeostasis genes depends on the intracellular copper concentration. Copper controls homeostasis genes through transcriptional regulation with cytoplasmic regulators or two-component systems (Arguello et al., 2013, Changela et al., 2003, Ma et al., 2009). Although bacteria also make use of post-transcriptional regulatory mechanisms notably based on riboswitches and small RNAs to ensure homeostasis of other transition metals (Dambach et al., 2015, Dann et al., 2007, Furukawa et al., 2015, Oglesby-Sherrouse and Murphy, 2013), such post-transcriptional regulation mechanisms are yet to be identified for copper.

*Bordetella pertussis* is a strictly aerobic, Gram-negative bacterium responsible for whooping cough (Belcher et al., 2021). Compared with other β proteobacteria, *B. pertussis* has lost most copper resistance mechanisms (Antoine et al., 2019, Rivera-Millot et al., 2021), probably because its specialized lifestyle as a host-restricted pathogen reduces its exposure to copper except when it is phagocytosed, a fate that it strives to avoid (Ahmad et al., 2019, Belcher et al., 2021, Kamanova et al., 2008). Transcriptomic analyses have identified a 3-gene operon predicted to participate in copper import in *B. pertussis, bp2923-22-21*, indicating that the bacterium needs to acquire copper in specific circumstances (Rivera-Millot et al., 2021). This three-gene operon is down-regulated by excess copper in the medium (Rivera-Millot et al., 2021). In this study, we characterized its regulation, which is original for transition metals, and revealed a post-transcriptional mechanism involving an upstream ORF widespread among Proteobacteria.

## Results and discussion

### Characterization of a Cu-regulated operon harbouring a TonB-dependent transporter gene in *B. pertussis*

RNA-seq experiments have identified a three-gene locus, *bp2923-bp2922-bp2921*, of which the last two genes are strongly down-regulated by copper (Rivera-Millot et al., 2021). The first open reading frame (ORF) is separated from the following gene by a long intergenic region (IGR) of 162 bp. The average (G+C) content of this locus, 71.6%, is higher than that of the *B. pertussis* genome (Fig. 1a). The three genes form an operon, as shown by RT-PCR on the IGR (Supplementary Fig. S1).

**Figure 1.**
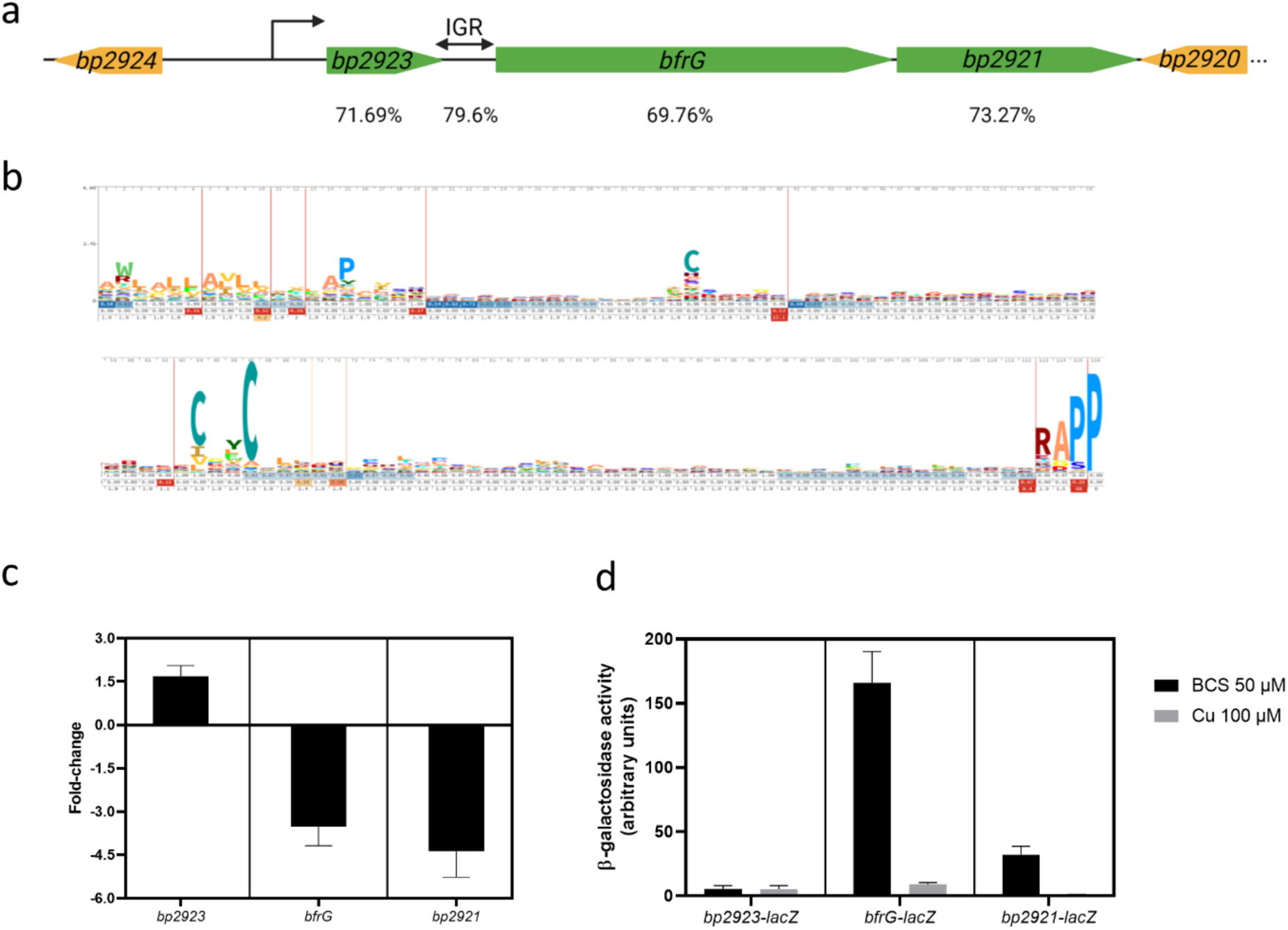
Role of *bp2923* in the regulation of *bfrG* and *bp2921* by copper. **a**, Schematic representation of the locus in *B. pertussis*. The transcription start site was identified by 5’RACE analyses. IGR represents the intergenic region between *bp2923* and *bfrG*, and the percentages given underneath the various regions of the operon indicate their (G+C) contents. **b**, HMM logo of the DUF2946 protein family. Two distinct motifs are well conserved, a putative copper-binding motif CxxC and a C-terminal RAPP motif. Another Cys residue is also semi-conserved. **c**, qRT-PCR analyses showing the ratios of transcription (in log_2_) of each gene in bacteria grown for 16 h in medium supplemented with 100 µM CuSO_4_ relative to bacteria grown in Cu-restricted medium (Cu chelator BCS added to 50 µM). The results were normalized against a housekeeping gene. **d**, translational *lacZ* fusions for the three genes. β-galactosidase activities were measured for bacteria grown in the same conditions as in (**c**). In panels **c** and **d**, the bars represent the means of three or four biological replicates, and the error bars show the standard deviations (SD).

*bp2923* encodes a putative 145-residue-long protein of unknown function of the DUF2946 Pfam family, predicted to be exported. Those proteins are characterized by two conserved sequence motifs, CXXC (where X represent non-conserved residues) and RAPP (Fig. 1b). *bp2922* is predicted to encode a TonB-dependent transporter (TBDT) previously named BfrG (Brickman et al., 2007). TBDTs form a large family of outer membrane transporters mediating import of various small molecules and notably iron, in the form of Fe-siderophore complexes or scavenged from host proteins (Noinaj et al., 2010). Finally, *bp2921* encodes a protein of the PepSY_TM family, a member of which has been described as a siderophore reductase (Josts et al., 2021). Proteomics analyses of *B. pertussis* extracts have identified peptides of the last two proteins but not of the *bp2923* gene product (Rivera-Millot et al., 2021).

To identify transcription start site(s) (TSS) in the *bp2923-2921* locus, we performed 5’ Rapid Amplification of cDNA Ends (5’ RACE) experiments. As the high (G+C) content of *bp2923* made it intractable for this technique, we introduced silent mutations to match its codon usage with that of *B. pertussis* and inserted the modified gene in the chromosome by allelic exchange, yielding the recombinant strain *BP2923*-OCU (<u>O</u>ptimized <u>C</u>odon <u>U</u>sage). 5’RACE analyses of the locus conducted in *BP2923*-OCU identified a TSS 45 bp before the potential initiation codon of *bp2923* (Fig. 1a; Supplementary Fig. S2). No additional TSS was identified between *bp2923* and *bfrG*, consistent with RT-PCR results showing transcripts that straddle the *bp2923-bfrG* intergenic region (Supplementary Fig. S1).

BfrG was detected by immunoblot analyses of cellular extracts of *B. pertussis* grown in the absence but not in the presence of copper, whereas Fe and Zn had no effect on its expression, indicating that the regulation of the operon is copper-specific (Expanded Fig. 1). Those results were confirmed by qRT-PCR experiments showing a dramatic reduction of *bfrG* mRNA abundance in the presence of Cu (Expanded Fig. 1). In contrast, Fe had no effect and Zn moderately affected *bfrG* mRNA levels, suggesting limited cross-regulation.

We performed qRT-PCR experiments on each of the three genes of the operon and normalized the results against a housekeeping gene. We also generated chromosomal reporter fusions by inserting *lacZ* in frame with the first codons of each gene to assess the effect of copper on their expression. *bfrG* and *bp2921* were expressed at moderate levels in copper-restricted medium (Fig. 1c,d). Addition of copper to the medium abolished their transcription and translation, consistent with RNAseq data (Rivera-Millot et al., 2021). Interestingly, *bp2923* was hardly translated despite being the first gene of the operon, and it did not appear to be regulated by copper (Fig. 1c,d). qRT-PCR analyses on *bfrG* at various times after Cu addition showed a fast decrease in transcript abundance (Supplementary Fig. S3).

### Importance of *bp2923* for post-transcriptional regulation of *bfrG* and *bp2921*

We investigated a potential regulatory role of the 5’ region of the operon by introducing a large, in-frame deletion of *bp2923* to avoid polar effects, yielding the recombinant BPΔ*2923* strain (Fig. 2a). We then tested the effect of copper on the expression of *bfrG* using both the *bfrG-lacZ* fusion and qRT-PCR. Copper regulation was abolished in BPΔ*2923*, and *bfrG* was expressed at lower levels than in the parental strain, BPSM (Fig. 2b). The introduction of *bp2923* at another chromosomal locus in BPΔ*2923* did not complement the deletion (Fig. 2b). Thus, its first position in the operon is required to control both the expression levels and the regulation by copper of the downstream gene. To determine if the Bp2923 protein or the *bp2923* mRNA was involved in this regulation, we tested the *bfrG-lacZ* fusion in BP*2923*-OCU, in which the protein sequence is intact but the mRNA sequence is modified (Fig. 2a). *bfrG* was expressed, although at a slightly lower level than in BPSM, and its expression responded poorly to Cu, suggesting a role of the mRNA sequence in regulation (Fig. 2b).

**Figure 2.**
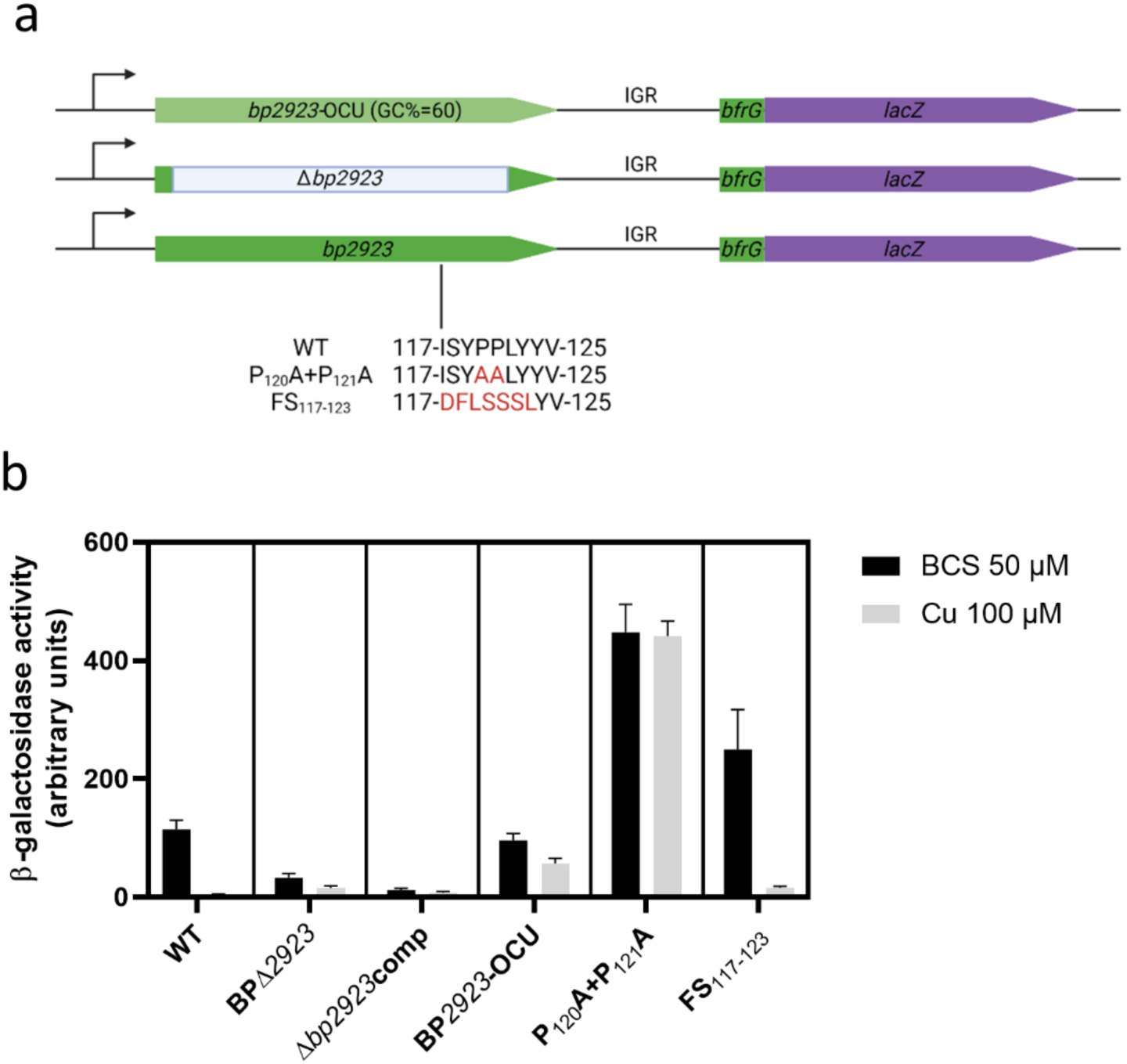
Effects of modifications of *bp2923* on *bfrG* expression. **a**, Mutations introduced in *bp2923. bp2923*-OCU represents the *bp2923* variant whose codon usage was optimized for *B. pertussis*. The deletion of *bp2923* (in grey) consists in an in-frame fusion between the first few and last few codons of the gene to avoid polar effects. P_120_A+P_121_A denotes the substitutions of two rare Pro codons by Ala codons, and FS_117-123_ indicates a frameshift obtained by introducing one nucleotide before codon 117 and removing one after codon 123. The resulting amino acid sequences are shown in red. The *bp2923* variants were introduced in the chromosome of *BPΔ2923* by allelic exchange. **b**, β-galactosidase activities of strains harbouring the chromosomal *bfrG-lacZ* fusion in various *bp2923* backgrounds. *Δbp2923*comp represents the complementation of *BPΔ2923* by *bp2923* under the control of its own promoter at a distinct chromosomal locus. The bars represent the means of four biological replicates, and the error bars show the standard deviations (SD).

Prediction of the mRNA structure of wt *bp2923* with MFold (http://www.unafold.org/mfold/applications/rna-folding-form-v2.php) indicated very stable potential stem-loop structures in the 5’ part of the operon, consistent with its high (G+C) content (Supplementary Fig. S4). Strikingly, an unstructured 29-bp sequence in the second moiety of *bp2923* has a skewed nucleotide content rich in C and T, resulting in several rare codons for *B. pertussis*. To alter the amino acid sequence of the unstructured mRNA region with minimal perturbation of the mRNA sequence and structure, we introduced reciprocal frameshift mutations (i.e., a frameshift mutation at the beginning of the target sequence followed by a frameshift mutation downstream of that sequence to restore the correct reading frame of the rest of the protein; mutant FS_117-123_; Fig. 2a, Supplementary Fig. S4). We also replaced two rare CCT (Pro) codons at positions 120 and 121 with frequent GCC (Ala) codons. The P_120_A+P_121_A mutations, which affect the mRNA structure in this region, caused overexpression of the *bfrG-lacZ* fusion and abolished its regulation by copper, unlike the FS_117-123_ mutations (Fig. 2a,b). Thus, the amino acid sequence encoded in this region appears to be unimportant, whereas mutations that generate secondary structures in the mRNA affect Cu regulation (Fig. 2b). The lack of structure in this mRNA stretch most likely contributes to the post-transcriptional regulation of the downstream genes.

### Rho-dependent transcription termination of the operon

Post-transcriptional regulation in bacteria may be mediated through transcription attenuation, which occurs by intrinsic or Rho-dependent mechanisms involving distinct mRNA signatures (Ray-Soni et al., 2016). The bacterial motor protein Rho is widely used to control expression of metabolic or stress response genes (Turnbough, 2019). Rho-utilization (rut) sites are unstructured mRNA regions to which Rho can bind, composed of repeated, C-rich patterns (Hao et al., 2021, Nadiras et al., 2018). Analyses of the nucleotide sequence of the locus revealed a so-called C>G bubble, *i*.*e*., a region where the percentage of C is higher than that of G on the coding strand, starting in the unstructured region of *bp2923* and ending in IGR (Fig. 3a). As C>G bubbles are indicative of Rho-dependent terminators we analysed the effect of copper on cultures treated with the Rho-specific inhibitor bicyclomycin using qRT-PCR. This treatment abolished the down-regulation of *bfrG* and *bp2921* by copper but had little effect on *bp2923* expression (Fig. 3b). This indicated that the presence of copper triggers Rho-dependent transcription termination within the *bp2923-bfrG-bp2921* operon between the first two genes.

**Figure 3.**
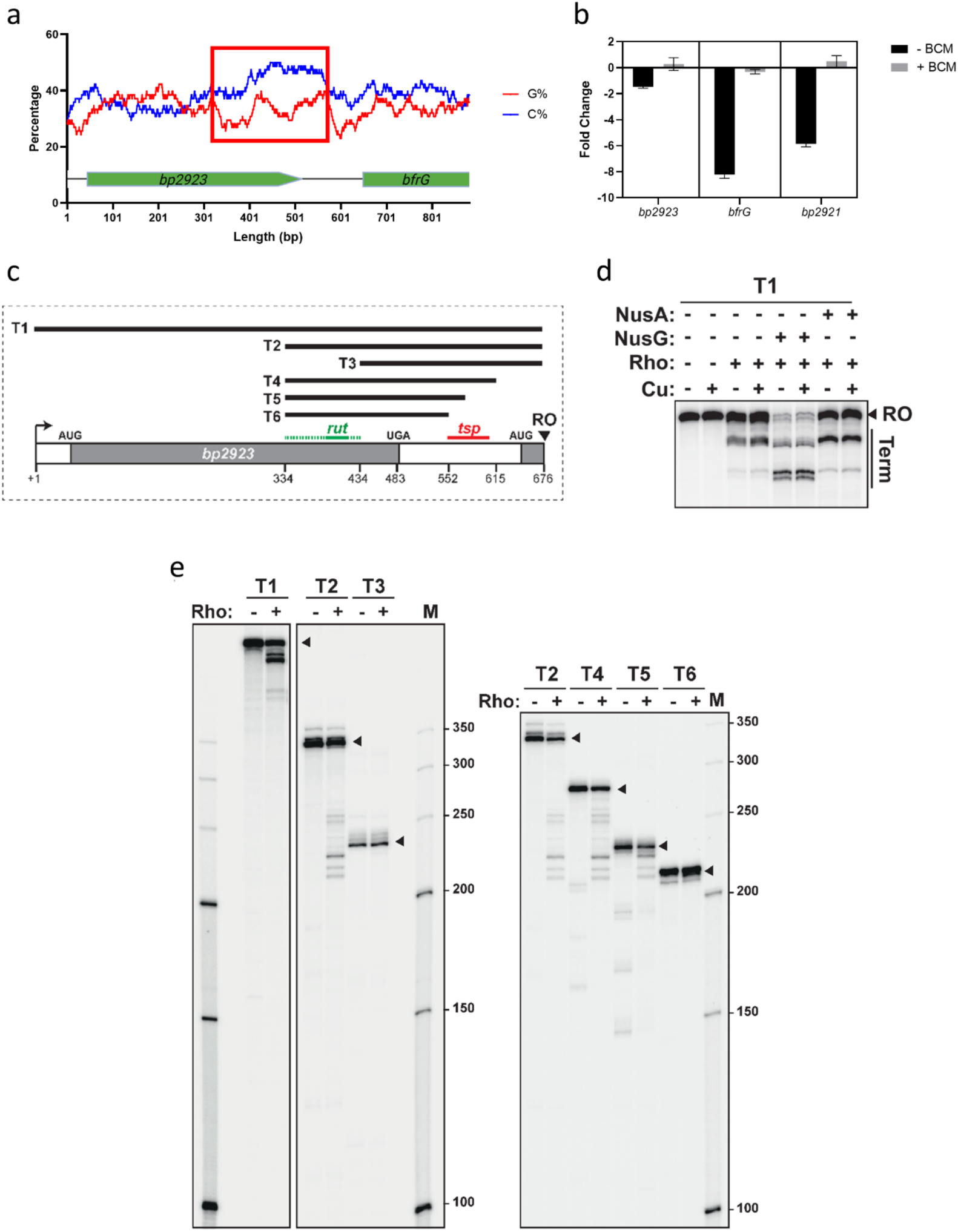
Rho-dependent termination between *bp2923* and *bfrG*. **a**, Percentages of C and G in the coding strand using a sliding window of 78 nucleotides from the TSS of *bp2923* to the beginning of *bfrG*. The C>G bubble is indicated with a red square. **b**, qRT-PCR analyses showing the ratios of transcription (in log_2_) of the 3 genes in bacteria grown in medium supplemented with 100 µM CuSO_4_ relative to bacteria grown in Cu-restricted medium (BCS added to 50 µM), with or without a 30-min treatment with bicyclomycin (BCM). The bars represent the means of three biological replicates and the error bars show the SD. **c**, Representation of the DNA templates used in the *in vitro* transcription experiments. The regions of the Rho-utilization site (rut) and of the transcription stop point (tsp) identified by the analyses shown in (**e)** are indicated. **d**, *In vitro* transcription experiments conducted on the T1 template show the presence of a Rho-dependent terminator in the *bp2923*-IGR region, the enhancement of transcription termination by NusG and NusA and the absence of effect of Cu on termination. RO denotes the run-off product. **e**, *In vitro* transcription experiments were conducted on all DNA templates to determine the rut and the tsp regions. Arrowheads indicate the RO products.

Rho-dependent transcription termination was confirmed by *in vitro* transcription experiments with a DNA template encompassing the sequence from the TSS upstream of *bp2923* to the first nucleotides of *bfrG*. Addition of Rho to the transcription reaction resulted in premature termination that was enhanced by the presence of factors known to facilitate Rho-dependent termination, NusA and NusG (Ray-Soni et al., 2016, Turnbough, 2019) (Fig. 3c,d). Copper did not affect transcription termination *in vitro* in those conditions, arguing that the mRNA does not sense copper by itself, as would be expected if it contained a riboswitch. By using DNA templates truncated from the 5’ or 3’ end, we mapped the transcription stop point (tsp) region in IGR and the putative rut site in *bp2923* in the C>G bubble region (Fig. 3c,e).

A Rho-dependent termination mechanism accounts for the effects of the mutations in the unstructured region (Fig. 2b). Thus, the frameshift did not affect regulation by copper because the rut site was preserved. In contrast, mutations that disrupt the rut site abolished regulation. The observation that they also increased the expression level of *bfrG* in the absence of copper suggests a background level of termination in the wt operon without added copper.

### Importance of Bp2923 protein for *bfrG* expression and regulation

We next investigated a potential role of the *bp2923*-encoded protein for regulation. We introduced nonsense codons at positions 50 or 133 to cause premature translation termination with minimal disruption of the mRNA sequence and structure (mutants Y_50_STOP and Y_133_STOP; Fig. 4a). Both mutations abrogated reporter activity of the *bfrG-lacZ* fusion (Fig. 4b), showing that premature release of the ribosome abolishes expression of the downstream gene even in the absence of Cu. The observation that expression and regulation of the downstream genes depends on *bp2923* translation indicates that this gene represents a new type of regulatory upstream ORF (uORF) (Dever et al., 2020).

**Figure 4.**
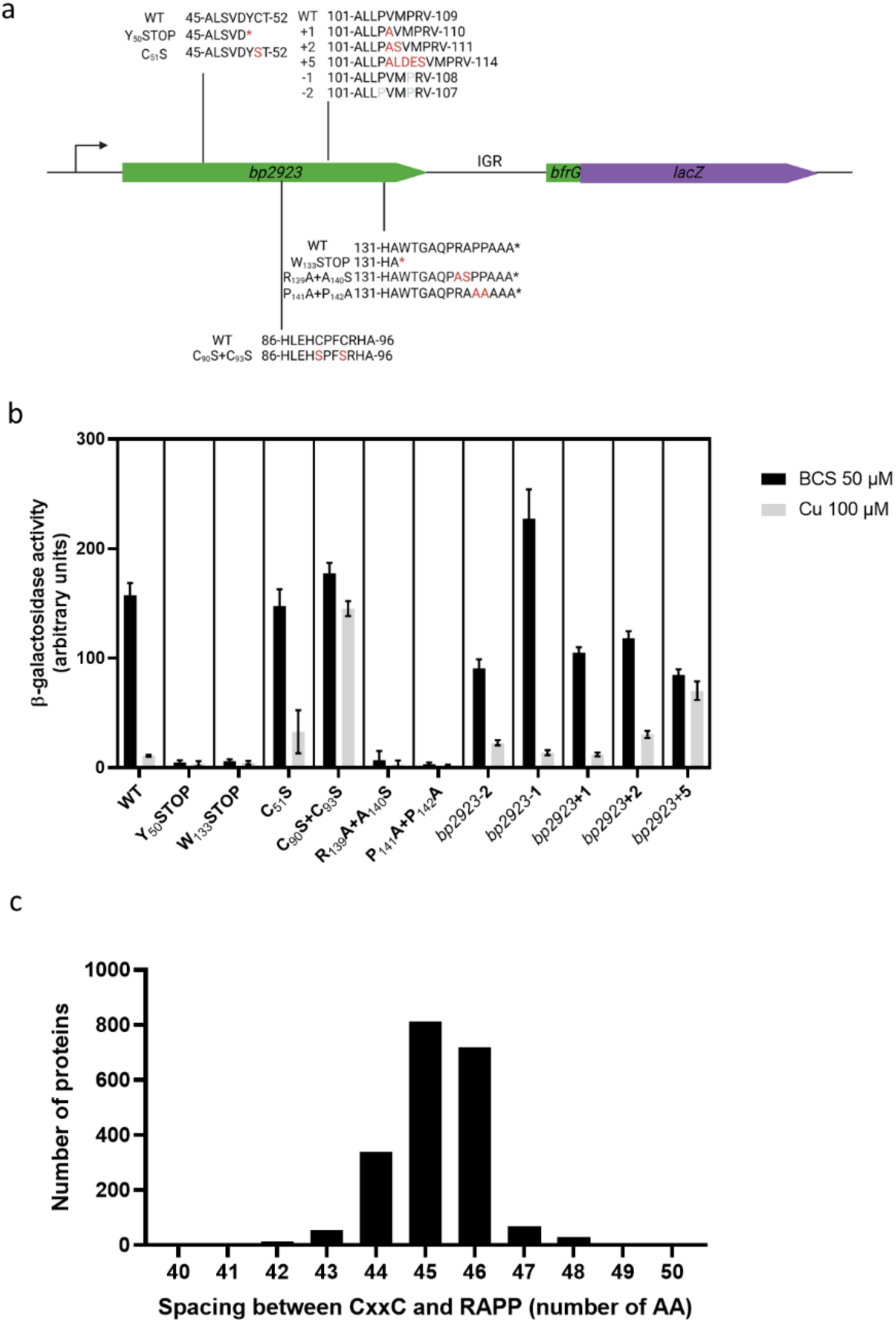
Role of conserved features of the Bp2923 protein for expression and Cu regulation of *bfrG*. **a**, Chromosomal mutations introduced in *bp2923*. The wt sequence in the region of interest is shown in first position in all cases. The point mutations, the nonsense mutation (denoted with *) or the insertions are indicated in red, and the deletions are in pale grey. We designed these insertions and deletions in such a way as to minimize their impact on the mRNA structure (Supplementary Fig. S4). **b**, β-galactosidase activities of strains harbouring a chromosomal *bfrG-lacZ* fusion in various *bp2923* backgrounds. The bars represent the means of four biological replicates, and error bars show the SD. **c**, Conservation of the spacing between the CxxC and the RAPP motifs in >2000 DUF2946-family proteins.

### Role of conserved features of the Bp2923 protein for regulation

The Bp2923 protein belongs to the DUF2946 family, which is characterized by two highly conserved sequence motifs. The RAPP motif, which is encoded by frequent *B. pertussis* codons, is located at positions 139-142 three residues before the C terminus (Fig. 1b). It is reminiscent of the C-terminal RAGP sequence of the so-called ‘arrest peptide’ of a well-known uORF that regulates *secA* expression in *Escherichia coli*, SecM (Ito and Chiba, 2013). Arrest peptides cause ribosome stalling, which affects the expression of downstream genes (Ito and Chiba, 2013). We thus tested the importance of the C-proximal RAPP motif by replacing Arg_139_ and Ala_140_ with Ala and Ser, or the two Pro with Ala residues (mutants R_139_A+A_140_S and P_141_A+P_142_A, respectively; Fig. 4a). Both sets of modifications abolished *bfrG* expression even in the absence of copper (Fig. 4b). This ‘constitutive’ termination of transcription shows that conversely, the RAPP sequence is required to promote transcription of the rest of the operon in copper-restricted conditions. Therefore, the conserved RAPP motif is most likely part of a ribosome arrest peptide involved in the regulation process. In enterobacteria, slow translation of consecutive Pro residues is alleviated by a specific elongation factor, EF-P (Doerfel et al., 2013). However, a knock-out mutation of this gene in *B. pertussis* had no effect on the expression of *bfrG* or its regulation by copper (Supplementary Fig. S5).

The other hallmark feature in the DUF2946 protein family, CXXC at positions 90-93, is a recognized Cu-binding motif. We replaced the two Cys residues with Ser and determined the effect of the SXXS sequence on regulation of *bfrG-lacZ* by copper (mutant C_90_S+C_93_S; Fig. 4a). These modifications did not affect the expression level of *bfrG* but abolished its control by copper, demonstrating the involvement of the CXXC motif in regulation (Fig. 4b). In contrast, replacement of a semi-conserved Cys in the DUF2946 family preserved *bfrG* regulation (mutant C_51_S; Fig. 4b).

The CXXC and the RAPP motifs are separated from each other by 45 intervening residues. Genome mining identified more than 2000 DUF2946 protein sequences in databases, mostly in Proteobacteria (Suppl. Table S1), and their analysis showed that the spacing between the two motifs is conserved to within one or two residues in the family (Fig. 4c). We therefore probed its importance for expression and regulation of *bfrG* by shortening or lengthening the spacing by one, two or five residues in the Bp2923 protein (mutants *bp2923-1, bp2923-2, bp2923+1, bp2923+2* and *bp2923+5*, respectively; Fig. 4a). We designed those mutations to minimize their effects on the mRNA structure (Supplementary Fig. S4). Deletion or addition of one or two residues moderately affected the *bfrG* expression levels, but not its regulation by copper. In contrast, addition of five residues abolished the effect of copper, showing that the two motifs must be adequately spaced for proper regulation, within a limited degree of variation (Fig. 4b).

Bp2923 is therefore a new regulatory protein that we propose to name CruR for <u>C</u>opper-<u>r</u>esponsive <u>u</u>pstream <u>R</u>egulator. The following model of a ligand-dependent relief of translation arrest is consistent with our data (Fig. 5). Following transcription of *bp2923*, the RNA polymerase most likely pauses in the intergenic region. Transcriptional pausing notably facilitates interactions with regulatory proteins, and termination stop points often coincide with pausing sites (Artsimovitch and Landick, 2000). In the absence of copper, stalling of the lead ribosome at the conserved RAPP motif of CruR prevents Rho from binding to the rut site and contacting the RNA polymerase, which enables the latter to resume transcription of the rest of the operon. The stalled ribosome is presumably rescued by a quality control mechanism (Keiler, 2008), as reported for SecM (Sunohara et al., 2004, Saito et al., 2022). In the presence of Cu in the cytoplasm, its perception by the invariant CXXC motif of nascent CruR relieves ribosome stalling by a mechanism that remains to be deciphered. The observation that the spacing between the two conserved motifs plays a critical role in regulation suggests that Cu binding to the nascent protein triggers some co-translational folding. Protein folding might exert a force that relieves stalling, as described in other systems (Kudva et al., 2018, Marino et al., 2016). Ribosome dissociation from the mRNA enables Rho to contact the RNA polymerase, causing transcription termination before *bfrG*. CruR is the first example of an uORF mediating post-transcriptional regulation in response to a transition metal.

**Figure 5.**
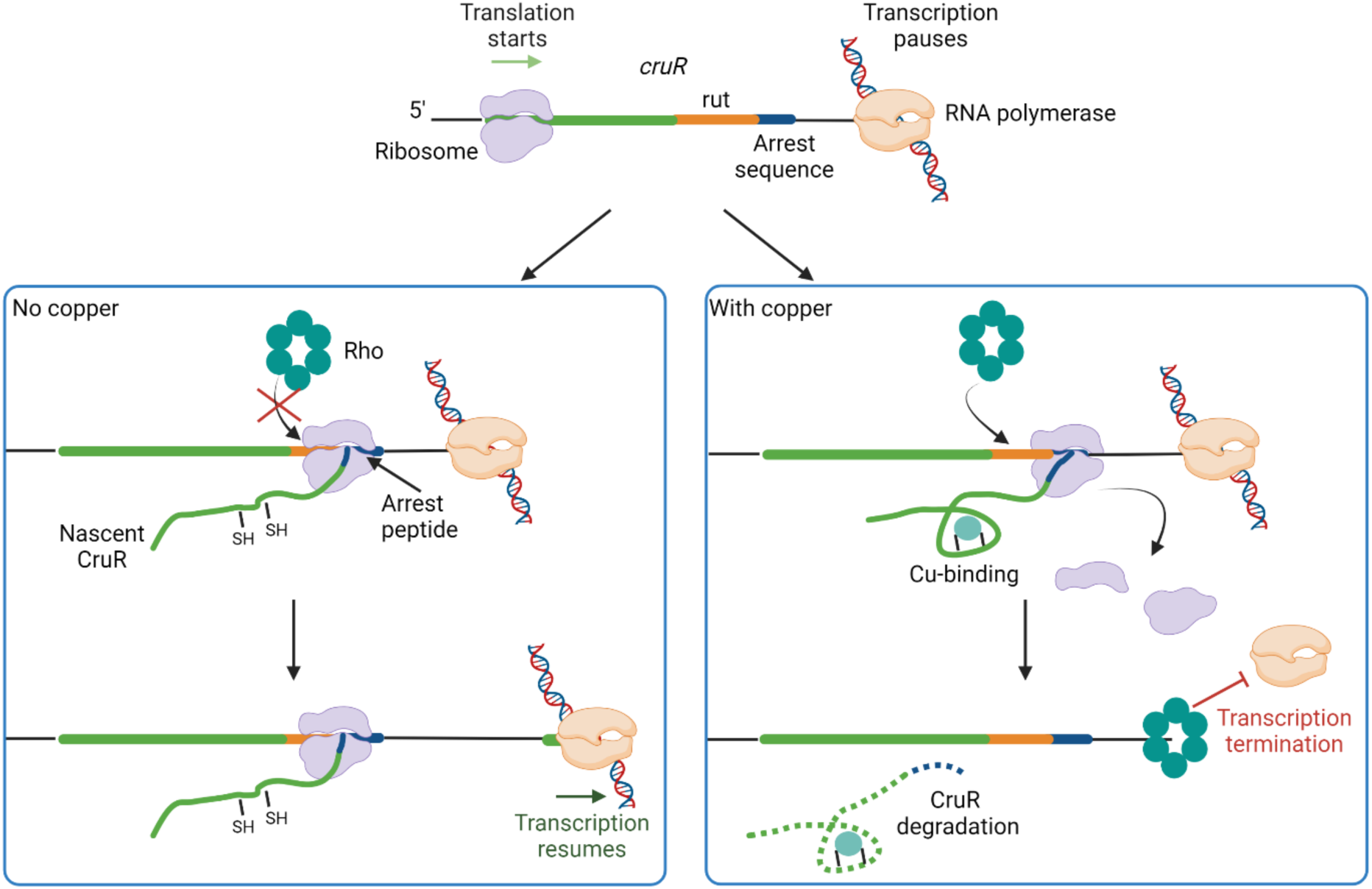
Model of *bfrG* regulation by the uORF CruR. The RNA polymerase pauses in the intergenic region between *cruR* and *bfrG*, and the lead ribosome translating CruR stalls at the RAPP motif. In the absence of copper (left panel), ribosome occupancy on this motif prevents Rho from binding to the rut site (in orange) on the mRNA or from accessing the RNAP polymerase, and transcription resumes. The perception of Cu (right panel) by nascent CruR through its CXXC motif relieves stalling of the ribosome. Rho can access the rut site and the RNA polymerase, leading to transcription termination. CruR is most likely degraded.

*In silico* analyses predicted the presence of an export signal, *i*.*e*., a signal-peptide or an N-terminal transmembrane segment with an N_in_-C_out_ orientation in more than 95% of all DUF2946 proteins. As the signal-peptide of SecM has been implicated in post-transcriptional regulation of the downstream *secA* gene (Ito and Chiba, 2013, Nakatogawa et al., 2004), we tested the possibility that the N-terminal region of Bp2923 similarly participates in copper regulation. We thus introduced reciprocal frameshift mutations to replace 40 residues encompassing the putative export signal by an out-of-frame sequence devoid of signal-peptide features while keeping the natural sequence of the rest of the protein (mutant FS_5-44_; Supplementary Fig. S6). This modification did not affect the expression or the regulation of *bfrG*.

CruR appears to be translated at very low levels (Fig. 1), and it was not detected in earlier proteomic analyses (Rivera-Millot et al., 2021). It is predicted to be poorly structured (Supp. Fig. S7), and our attempts to produce it as a recombinant protein were unsuccessful. All these observations indicate that mature CruR is probably short-lived and therefore unlikely to exert an additional function in *B. pertussis*. CruR is most likely rapidly degraded given that C-terminal non-polar residues constitute a signal for degradation by various proteases (Parsell et al., 1990, Keiler et al., 1995). Notably, metal-binding proteins and regulatory proteins are ranked as fast-degrading in *E. coli* (Nagar et al., 2021).

### Genes in synteny with *bp2923* homologues

*In silico* analyses revealed that CruR homologues are widespread among β, γ and α Proteobacteria (Suppplementary Table S1). A few were also found in Planctomycetes, Firmicutes and Deinococcus. In most cases, potential operonic structures were identified with these genes in first position (Expanded Fig. 2). *bp2923* homologues frequently precede TBDT- and/or PepSY_TM-coding genes, as in *B. pertussis*, indicating a widespread genetic organization. Among those TBDTs are OprC-type transporters (signature TIGR01778), one of which was recently shown to be specific for Cu (Bhamidimarri et al., 2021). Other genes frequently found in operons with *bp2923* homologues code for proteins involved in Cu transport, binding or homeostasis, including the chaperones ScoC and Pcu_A_C that participate in the assembly of heme-copper subunits of respiratory complexes (Serventi et al., 2012, Trasnea et al., 2016), and the Cu-binding proteins YcnI and CopC (Damle et al., 2021, Lawton et al., 2016). Some of these operons also comprise MbnPH-like genes notably found in biosynthetic operons of Cu-binding methanobactin-type molecules (Kenney and Rosenzweig, 2018, Manesis et al., 2021), and AhpC_TSA genes, whose products detoxify peroxides (Dubbs and Mongkolsuk, 2007). Long intergenic distances between *cruR* homologues and the following genes are generally observed. Notably, other types of uORFs are also followed by long intergenic regions (Choi et al., 2021), suggesting that this feature is necessary for regulation. Such distances are incompatible with translational coupling between *cruR* and the following gene, which implies *de novo* translation initiation of the next gene. In the *B. pertussis* case, this is consistent with the higher levels of activity of the BfrG translational fusion compared with the CruR fusion.

As *cruR* genes are overwhelmingly found in first position of putative operons linked to Cu homeostasis, and the two functional motifs of the proteins and their spacing are extremely well conserved, *cruR* is most likely the paradigm of a new family of uORF involved in post-transcriptional regulation in response to copper. There are indeed indications that our model of CruR being a copper-responsive uORF that regulates copper homeostasis genes may apply to other cases. The *oprC* gene in *Pseudomonas aeruginosa* is preceded by a *cruR* homologue down-regulated by Cu (Yoneyama and Nakae, 1996). In *Bradyrhizobium japonicum*, an operon for cytochrome oxidase biogenesis induced by copper starvation is also headed by a *cruR* homologue (Serventi et al., 2012).

Other biologically relevant transition metals can regulate gene expression through post-transcriptional mechanisms involving sRNAs or riboswitches (Bossi et al., 2020, Chareyre and Mandin, 2018, Dambach et al., 2015). It is thus interesting that no riboswitches have been described for transition metals at the top of the Irving-Williams series, Cu and Zn. Known metallo-riboswitches have micromolar affinities for their ligands (Furukawa et al., 2015, Price et al., 2015). The extremely low levels of free intracellular copper would prevent low-affinity riboswitches from outcompeting high-affinity copper-binding proteins (Changela et al., 2003), which may be the reason why protein-based post-transcriptional regulation mechanisms have evolved for copper. There are preliminary indications that other families of uORFs post-transcriptionally regulate genes coding for copper export or detoxification systems in response to this metal (de Freitas et al., 2019, Rademacher et al., 2012, Voloudakis et al., 2005). In *Rhodobacter capsulatus*, the activity of a multi-copper oxidase, CutO, was shown to depend on an upstream ORF with similarities to CruR, CutF, which the authors conjectured works as a Cu chaperone (Öztürk et al., 2021). It is tempting to speculate that CutF may also be an uORF that regulates the expression of downstream genes in response to copper. Many variations on the theme of Cu-responsive post-transcriptional regulation by uORFs most likely remain to be discovered.

## Materials and methods

### Bacterial strains and culture conditions

*B. pertussis* strains were grown on Bordet-Gengou (BG) agar supplemented with 10% sheep blood for 48 hours at 37°C, and then in modified Stainer-Scholte (SS) liquid medium at 37°C with agitation. SS medium was supplemented with 50 µM of the copper chelator bathocuproine disulfonate (BCS) to limit copper availability or with 100 µM CuSO_4_, FeSO_4_ or ZnSO_4_ as indicated. Antibiotics were added at final concentrations of 100 µg/mL streptomycin, 10 µg/mL gentamycin, 30 µg/mL nalidixic acid, 150 µg/mL ampicillin and 25 µg/mL kanamycin. Where indicated, cultures were treated with 20 µg/mL bicyclomycin (Santa Cruz Biotechnology) for 30 min at mid-log phase.

### Construction of mutant strains

Deletion mutants of *cruR* and of *bfrG* were constructed by amplifying their flanking regions as EcoRI-XbaI and XbaI-HindIII fragments and cloning the two amplicons in tandem in pSS1129 (Stibitz, 1994). The *bp2923*-OCU modified gene was purchased from GeneCust as a synthetic gene also containing the flanking regions of *cruR* (*bp2923)* and introduced in pSS1129. Recombinant pSS1129 plasmids were used to transform *E. coli* SM10 for conjugation with *B. pertussis* BPSM, in order to perform allelic exchange. Antibiotic selection was then conducted appropriately to select the recombinant strains. To construct the *cruR-lacZ* fusion, the sequence encompassing *bp2924*, the intergenic *bp2924-cruR* region, and the first 10 codons of *cruR* was amplified using oligonucleotides carrying EcoRI and XhoI sites, respectively, and the amplicon was cloned in pQC2123 (Chen et al., 2018) digested with EcoRI and SalI. For *bfrG-lacZ*, the amplicon included *bp2924, cruR*, IGR and the first 10 codons of *bfrG*. For *bp2921-lacZ*, the amplicon included 600 bp upstream of *bp2921* and its first 10 codons. Frameshift mutations, the C_51_S mutation and the mutations of the P_120_-P_121_ residues were introduced by site-directed mutagenesis of *cruR* on pUC57-*cruR* using the kit QuikChange II XL. This plasmid carries a synthetic 800-bp EcoRI-XhoI fragment starting in *bp2924* and ending after the first 10 codons of *bfrG*. The mutated fragments were excised with the same enzymes and introduced in pQC2123 as above. Synthetic gene fragments were ordered from GeneCust to introduce the Y_50_STOP, C_90_S+C_93_S, P_120_A+P_121_A, R_139_A+A_140_S and P_141_A+P_142_A mutations in *cruR*. Using the natural NcoI site in *cruR*, the EcoRI-NcoI or NcoI-XhoI fragments of pUC57-*cruR* were replaced with the relevant mutated fragments, and the complete EcoRI-XhoI fragment was ligated with pQC2123. All pQC2123 variants were introduced in BPΔ*2923* by conjugation and integrated in its chromosome by homologous recombination using the 600-bp sequence upstream of *cruR* as the recombination region. For complementation of *cruR* in trans in the BP*β2923* chromosome, the BamHI-XbaI fragment of pRM1 was replaced by an amplicon encompassing *cruR* and its promoter region. pRM1 derives from pXR1 (Kammoun et al., 2013), from which the HindIII-ApaLI fragment was replaced with a synthetic fragment containing a 666-bp HindIII-BamHI portion of *ureJ*, a central 3191-bp BamHI-XbaI portion of *fhaB* and a 1064-bp XbaI-ApaLI portion of *ureC. bp2923* was thus introduced at the inactive *ure* locus of *B. pertussis* by homologous recombination. A knock-out mutant of *efp* was constructed by interrupting the gene with a recombinant pFUS2 suicide plasmid carrying an internal sequence of *efp* using homologous recombination. The plasmids and oligonucleotides are described in Supplementary Tables S2 and S3.

### Immunoblot analyses

The bacterial pellets obtained from 10-mL *B. pertussis* cultures grown overnight to an OD_600_ of 1-1.5 in the indicated conditions were resuspended to an OD_600_ of 5 in 50 mM Tris-HCl (pH 8) and lysed using a Ribolyser at speed 6 for 50 s. Laemmli buffer was added to the clarified lysates, and samples were heated at 95°C for 10 min before SDS-PAGE. The proteins were transferred on a nitrocellulose membrane and BfrG was detected by immunoblotting using a polyclonal antibody produced in guinea pig (Eurogentec, Belgium) at a 1:2,500 dilution, followed with anti-guinea pig-HRP antibodies at a 1:5,000 dilution. Blots were revealed using the Amersham ECL Prime Western Blotting System with the Amersham Imager 600 (General Electrics). Loading controls were run on a separate gel stained with Coomassie Blue.

### RNA techniques

8 mL of liquid *B. pertussis* cultures grown in Cu-restricted or Cu-supplemented media to an OD_600_ of 1-1.5 were centrifuged at 4000 rpm at 4°C for 10 minutes after adding 2 mL of a 95/5 ethanol/phenol mix. Pellets were flash-frozen in liquid nitrogen and kept at -80°C. For qRT-PCR, RNA extraction was performed by using Tri-Reagent (InVitrogen), followed by a DNAse I treatment (Sigma Aldrich) to remove remaining genomic DNA. Retro-transcription was performed with the Verso cDNA synthesis kit (ThermoFisher). qRT-PCR was performed in a Roche LightCycler 480 Instrument II using the Takyon LowROX SYBR kit (Eurogentec). All qRT-PCR experiments were conducted with 3 biological replicates and 3 technical replicates, and data were normalized using the housekeeping gene *bp3416*. For the detection of transcripts in the IGR, a PCR was performed on 75 ng of total cDNA, and 75 ng of DNAse-treated RNA or purified genomic DNA as controls. 5’RACE experiments were conducted on total RNA extracted from cultures of *BP2923*-OCU supplemented with BCS, using the Generacer kit (Invitrogen) and specific RACE primers for *bfrG* and *cruR* according to the manufacturer’s instructions. For *cruR*, after PCR amplification of the cDNA with a first primer annealing within the gene, a nested PCR was performed using a second primer annealing immediately before the *cruR* start codon to enhance specificity. The cDNA isolated in the RACE experiment was used to build a library using the Illumina TruSeq Stranded RNA LT library preparation kit, followed by sequencing on an Illumina NextSeq 500 benchtop sequencer. The GeneRacer adapter sequence was removed from the reads using Cutadapt (https://github.com/marcelm/cutadapt) and the reads were mapped using the CLC Genomics software (Qiagen).

### β-galactosidase activity measurements

*B. pertussis* strains carrying chromosomal translational fusions with *lacZ* were cultured to an OD_600_ of 1.5-2 in the indicated conditions and harvested by centrifugation. Pellets were resuspended to an OD_600_ of 5 and lysed using a Ribolyser at speed 6 for 50 s. β-galactosidase activity was measured as previously described (Antoine et al., 2000). Experiments were conducted with 4 biological replicates and 3 technical replicates.

### Transcription termination experiments

DNA templates T1 to T6 which contain distinct parts of the 5’UTR-*cruR-*IGR region were prepared by standard PCR procedures (Simon et al., 2021). Briefly, recombinant pQC2123 with the wt locus sequence was amplified with pairs of forward and reverse primers. Forward primers allow introduction of the sequence of the T7A1 promoter upstream of the probed *B. pertussis* sequence. Purification of the Rho, NusA and NusG proteins from *E. coli* was described (Simon et al., 2021). These proteins were used as proxies for their *B. pertussis* counterparts to seek Rho-dependent termination sites within the 5’UTR-*bp2323*-IGR region. Standard transcription termination experiments were performed with minor modifications. Briefly, DNA template (0.1 pmol), *E. coli* RNA polymerase (0.3 pmol; New England Biolabs), Rho (0 or 1.4 pmol hexamers), NusA (0 or 2.8 pmol), NusG (0 or 2.8 pmol), Superase-In (0.5 U/µL; Ambion), and CuCl_2_ (0 or 10 µM, final concentration) were mixed in 18 µL of transcription buffer (40 mM Tris-HCl, pH 8.0, 5 mM MgCl_2_, 1.5 mM DTT, and 100 mM KCl). DTT in the transcription buffer reduces Cu^2+^ to Cu^1+^, the latter being the form found in the bacterial cytoplasm. Mixtures were incubated for 10 min at 37°C before addition of 2 µL of initiation solution (250 µg/mL rifampicin, 2 mM ATP, GTP, and CTP, 0.2 mM UTP, and 2.5 µCi/µL α[^32^P]UTP in transcription buffer). After 20 min of incubation at 37°C, transcription reactions were stopped by the addition of 4 µL of EDTA (0.5 M), 6 µL of tRNA (0.25 mg/mL), and 80 µL of sodium acetate (0.42 M), followed by ethanol precipitation. Reaction pellets were resuspended in loading buffer (95% formamide; 5 mM EDTA) and analysed by denaturing 7% polyacrylamide gel electrophoresis and by phosphorimaging with a Typhoon FLA-9500 instrument and ImageQuant TL software (GE Healthcare). Potential Cu^2+^ scavenging-by-buffer effects (Mash et al., 2003) were ruled out in control transcription termination experiments where Tris-HCl was replaced by MOPS, pH 7.9 (not shown).

### *In silico* analyses

The non-redundant NCBI database (release of December 2020) was searched for the occurrence of DUF2946 using its PFAM hmm profile (PF11162) and the hmmsearch program (http://hmmer.org/). The results were curated to retain proteins less than 80% identical in sequence using cd-hit (http://cd-hit.org). Using a locally built database containing all the bacterial GenBank files of NCBI, the 5 genes flanking the DUF2946 genes on either side were retrieved irrespective of their distance and translated, and the corresponding protein identification numbers were attached. The sense of transcription of the neighbouring genes and their intergenic distances were determined. Putative operons harbouring genes transcribed in the same direction as the DUF2946-coding gene were retained. The numbers of occurrences of given Pfam domain-coding genes at each position relative to the gene of interest, itself found at position ‘0’ of each locus, were then computed. The CLC software was used for sequence manipulation and analyses such as Pfam domain prediction.

## Acknowledgements

We thank Dr Qing Chen (FDA, USA) for the kind gift of the pQC2123 plasmid, and Dr Axel Innis (IECB Bordeaux) for discussions. This work was funded by the Institut National de la Santé et de la Recherche Médicale (INSERM) and the University of Lille. G. Roy acknowledges the support of a doctoral fellowship from the University of Lille.

## Author contributions

GR, RA, FJD conceived the study; GR, RA, AS, SS, ARM performed the experiments; GR, RA, MB, FJD analyzed the data; GR, MB and FJD wrote the paper; all authors reviewed the paper.

## Expanded figure legends

**Expanded Figure 1.**
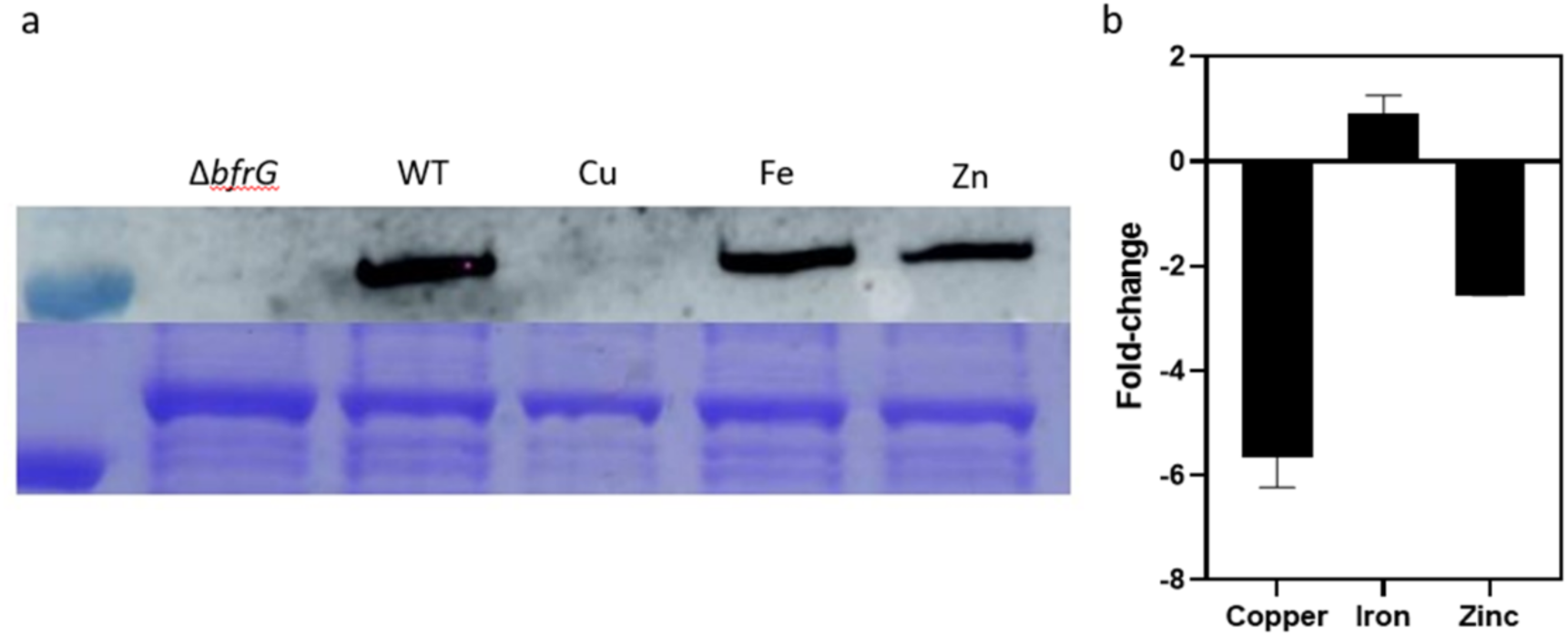
Specificity of the regulation of *bfrG* by metals. **a**, Analysis of *B. pertussis* extracts by immunoblotting using anti-BfrG antibodies. A Coomassie-stained gel underneath shows a loading control. **b**, Ratios of transcription (in log_2_) of the 3 genes in bacteria grown in medium supplemented with 100 µM CuSO_4_, FeSO_4_ or ZnSO_4_ relative to bacteria grown in standard medium.Data represent the means of three biological replicates, and the error bars show the SD.

**Expanded Figure 2.**
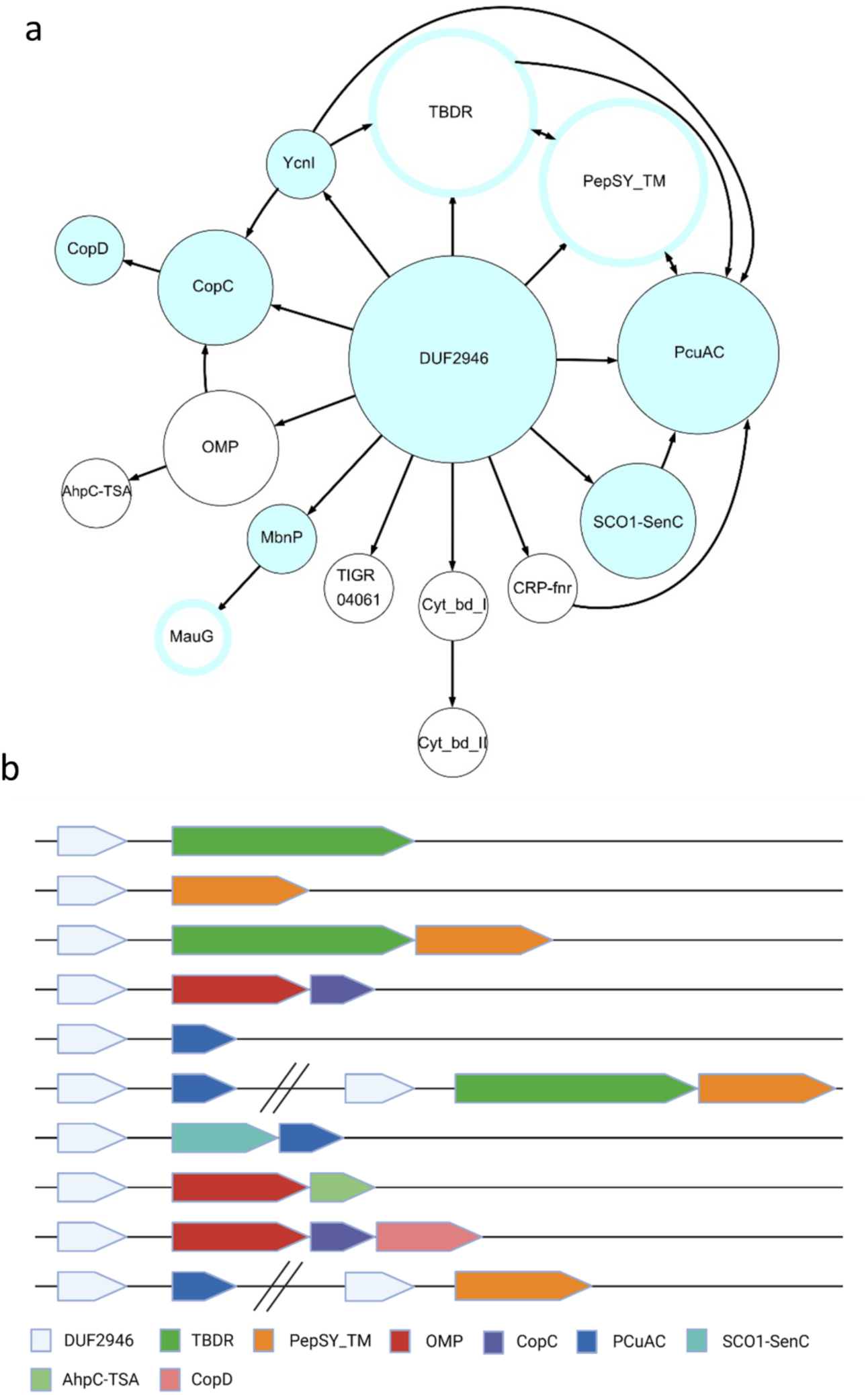
Genetic environments of *cruR* homologues. **a**, DUF2946-coding genes were systematically found in first positions of putative operons that comprise genes coding for proteins of the indicated families, represented by circles whose sizes correspond to the current numbers of occurrences of each protein type in those operons (large circle, 400-1000; medium-size, 50-399; small circles, 1-49). Blue-filled circles represent protein families involved in copper homeostasis or utilization, and those surrounded by blue lines represent families some members of which are found in copper-related operons, while others are involved in distinct processes. The arrows indicate the order of the genes in the putative operons. **b**, Representation of the most frequent genetic organizations. Parallel bars separate potentially distinct transcriptional units at the same locus.

