## Supplementary material for "Post-transcriptional regulation by copper with a new upstream Open Reading Frame": Suppl figures and tables

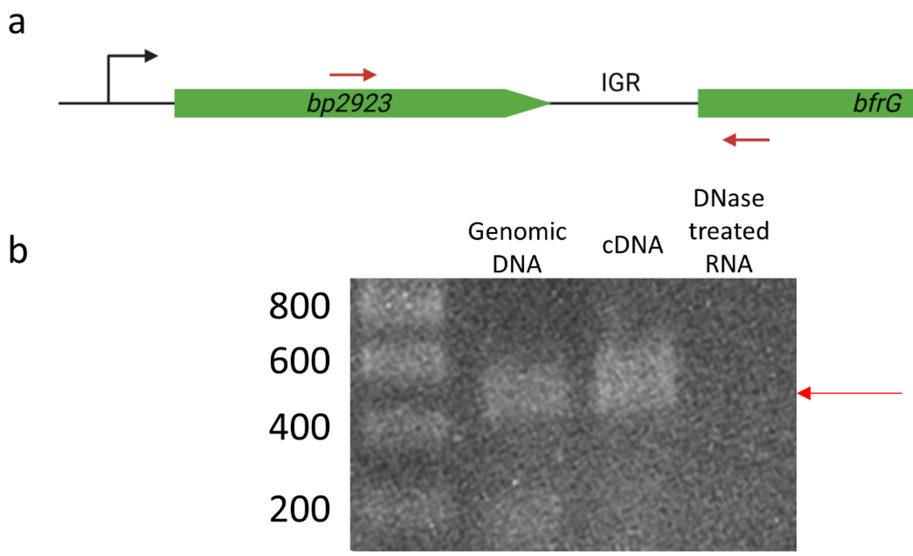


**Figure S1. Operonic structure of *bp2923* and *bfrG*. a,** Localization of the primers used for the PCR reactions (red arrows). **b,** The PCR were conducted with the primers shown above to amplify a 458-bp region straddling IGR. Reverse transcription followed with PCR was conducted on DNAse-treated RNA extracted from a mid-log phase culture of BPSM grown in copper-restriction conditions (i.e., addition of BCS to 50 μM) (lane denoted cDNA). PCR was also conducted on genomic DNA, and on DNAse-treated RNA without reverse transcription as positive and negative controls, respectively. The position of the amplicon of the expected size after RT-PCR ( red arrow) shows that the presence of transcripts in IGR, indicating that *bp2923* and *bfrG* are in operon.

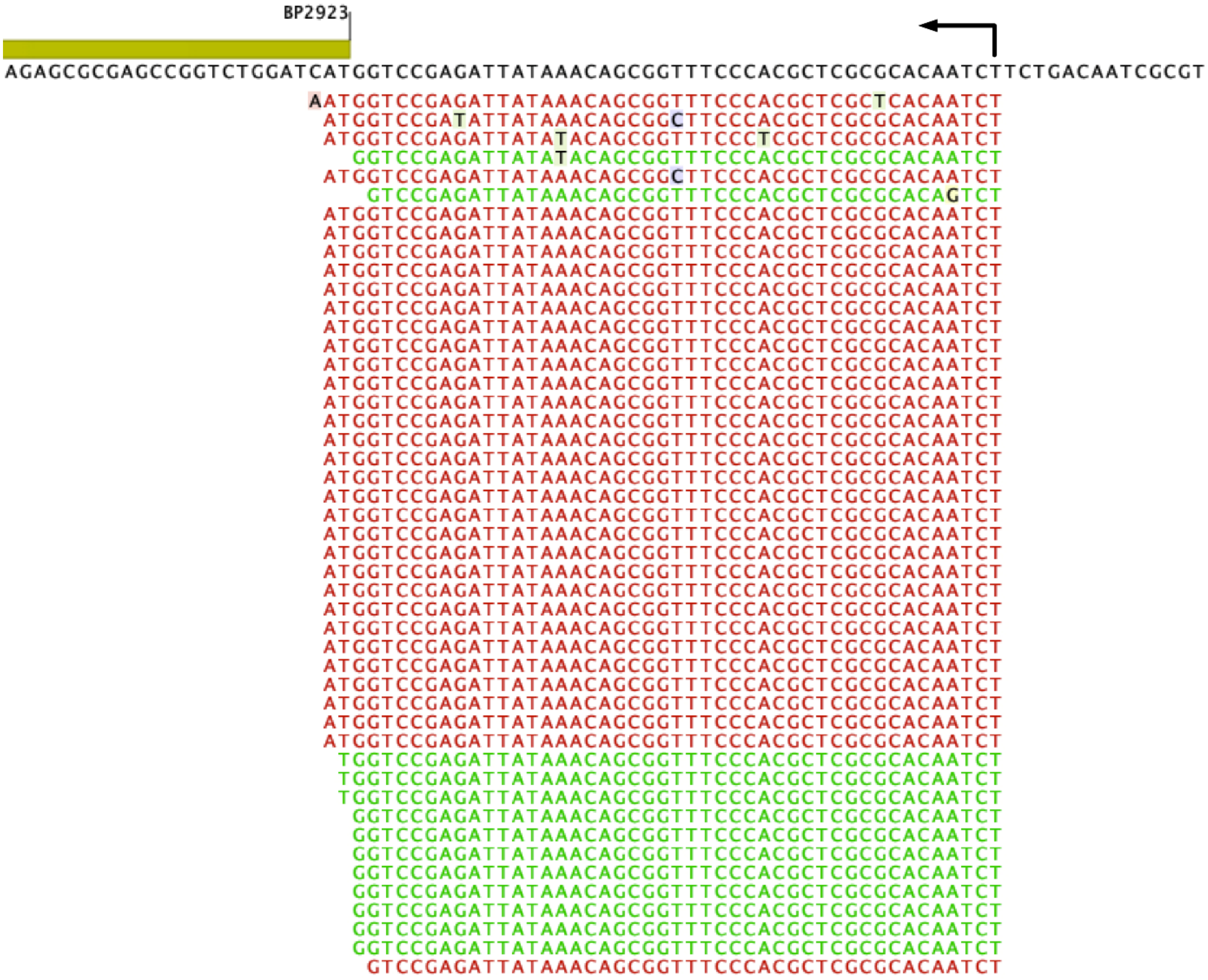

**Figure S2. Determination of the operon transcription start site (TSS)**. 5'RACE experiments were conducted using successively a primer located in *bp2923* and a nested primer starting directly upstream of the *bp2923* start codon. The screen shot shows the reads from the sequencing reaction. The arrow indicates the TSS. The sense of transcription is from right to left.

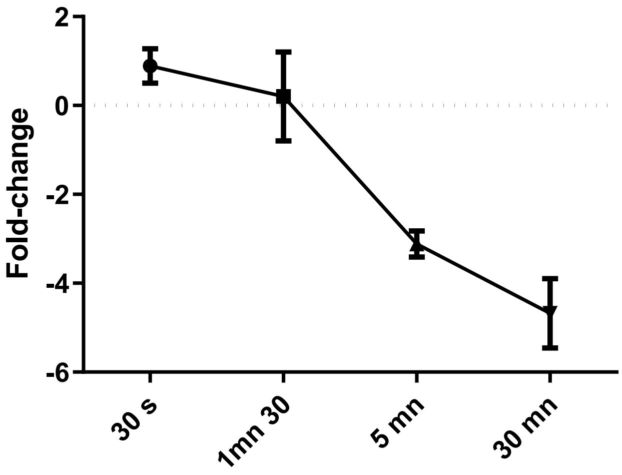


**Supplementary Figure S3**. **Kinetics of *bfrG* regulation by copper.** Aliquots were taken at the indicated time after addition of 100 µM $CuSO_4$ to liquid cultures of the wt strain BPSM for qRT-PCR analyses on *bfrG*. Data represent the means of three biological replicates, and the error bars show the SD.

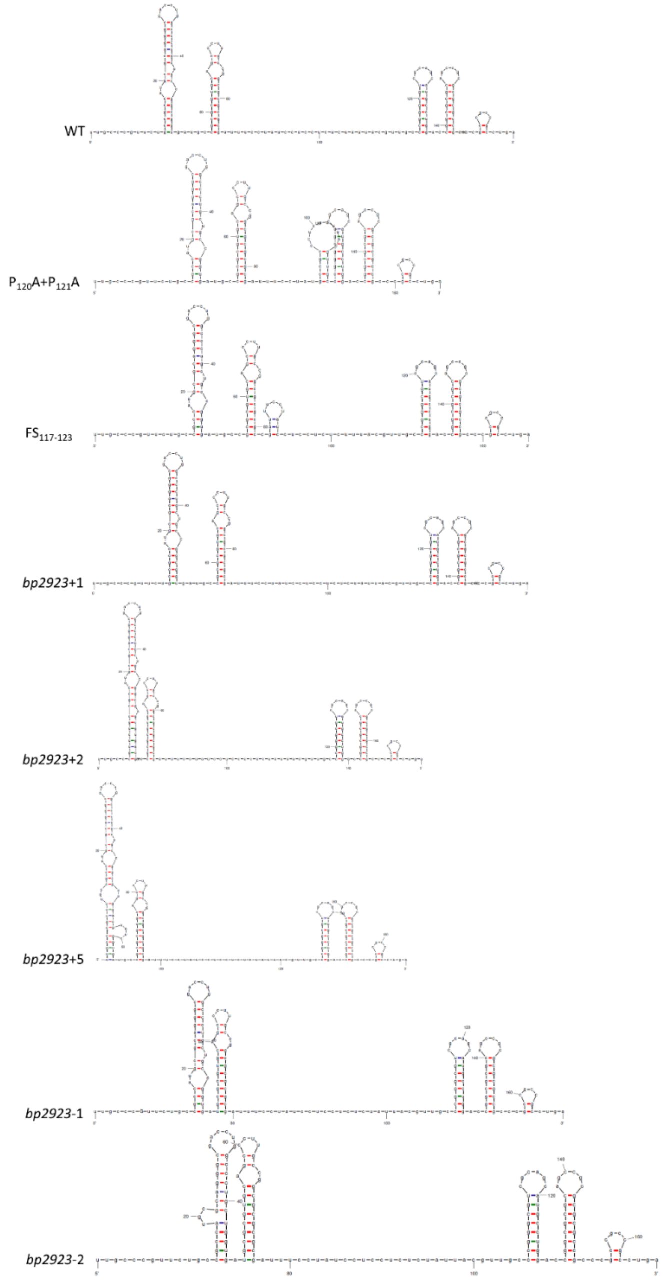


**Figure S4. Predictions of the mRNA structure of *bp2923* and variants generated in this work.** The wt sequence and the sequences of mutants were analysed between the region coding for the CXXC motif and the STOP codon of *bp2923* using the mfold server (http://www.unafold.org/mfold/applications/rna-folding-form-v2.php).

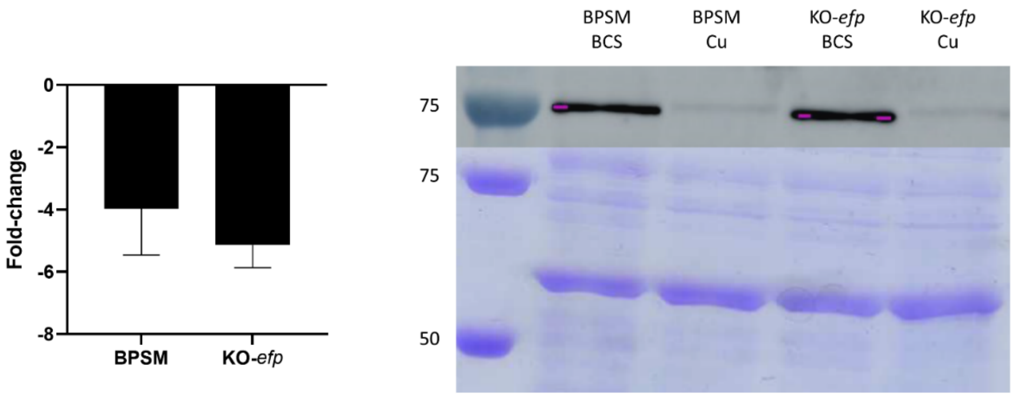


**Supplementary Fig. S5. Effect of an EF-P knock-out on *bfrG* regulation.** Left panel, qRT-PCR analyses showing the ratios of transcription of *bfrG* in bacteria grown for 16 h in medium supplemented with 100 µM $CuSO_4$ relative to bacteria grown in Cu-restricted medium (BCS added to 50 µM). The results were normalized against a housekeeping gene. Right panel, analysis of *B. pertussis* extracts by immunoblotting using anti-BfrG antibodies. The bacteria were grown for 16 h in SS medium supplemented with 100 µM of Cu or with 50 µM BCS. A Coomassie-stained gel underneath shows an unidentified protein band as a loading control.

a

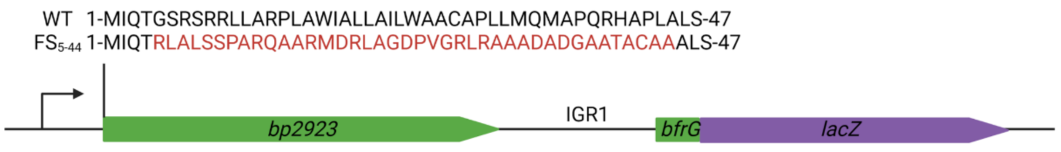


b

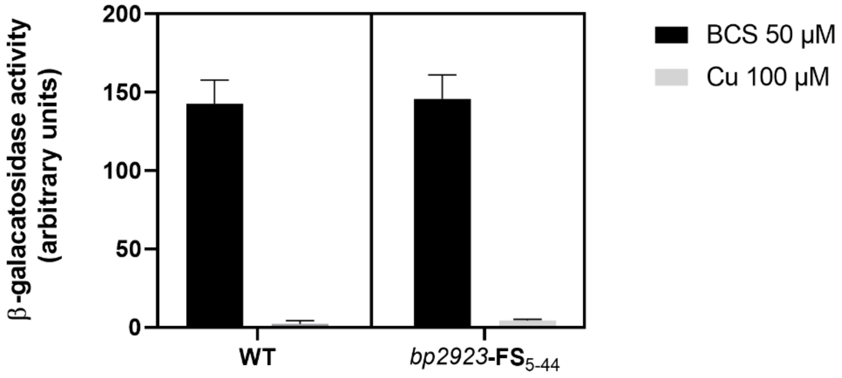


**Supplementary Figure S6. Replacement of the predicted signal-peptide of Bp2923 by reciprocal frameshift mutations. a,** Wt and mutant protein sequences in that region. **b,** β-galactosidase activities of strains harboring the chromosomal *bfrG-lacZ* fusion in the indicated *bp2923* backgrounds. The bars represent the means of four biological replicates, and the error bars show the SD.

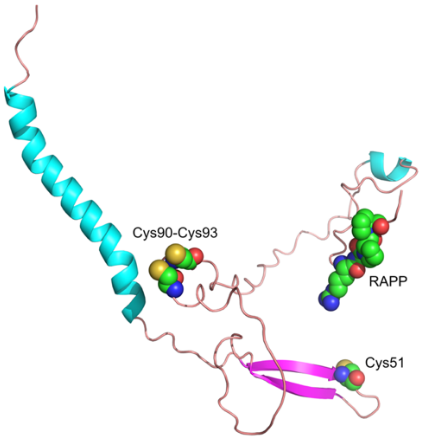


**Supplementary Figure S7. Predicted structure of the Bp2923 protein.** The signal-peptide and the residues of interest are in cyan and in space filling, respectively. The prediction was conducted with AlphaFold2.
(https://colab.research.google.com/github/sokrypton/ColabFold/blob/main/AlphaFold2.ipynb)

**Supplementary Table S1. List of CruR homologues and taxonomy of the host organisms**

| | | | | |
|---|---|---|---|---|
| gi\|WP_161882635.1 | Deinococcus alpinitundrae | Deinococci | Deinococcales | Deinococcaceae |
| gi\|CDQ34181.1 | Virgibacillus halodenitrificans | Bacilli | Bacillales | Bacillaceae |
| gi\|WP_074041421.1 | Kiritimatiella glycovorans | Kiritimatiellae | Kiritimatiellales | Kiritimatiellaceae |
| gi\|EKJ99229.1 | Rhodopirellula baltica SH28 | Planctomycetia | Planctomycetales | Planctomycetaceae |
| gi\|WP_157605409.1 | Schlesneria paludicola | Planctomycetia | Planctomycetales | Planctomycetaceae |
| gi\|WP_056053347.1 | unclassified Caulobacter | Alphaproteobacteria | Caulobacterales | Caulobacteraceae |
| gi\|OYW30099.1 | Caulobacter sp. 12-67-6 | Alphaproteobacteria | Caulobacterales | Caulobacteraceae |
| gi\|WP_010919643.1 | Caulobacter vibrioides | Alphaproteobacteria | Caulobacterales | Caulobacteraceae |
| gi\|WP_062142995.1 | Caulobacter henricii | Alphaproteobacteria | Caulobacterales | Caulobacteraceae |
| gi\|WP_035045921.1 | Caulobacter henricii | Alphaproteobacteria | Caulobacterales | Caulobacteraceae |
| gi\|WP_125157032.1 | Caulobacter sp. 602-1 | Alphaproteobacteria | Caulobacterales | Caulobacteraceae |
| gi\|ACK49443.1 | Methylocella silvestris BL2 | Alphaproteobacteria | Hyphomicrobiales | Beijerinckiaceae |
| gi\|WP_129396367.1 | Methylovirgula ligni | Alphaproteobacteria | Hyphomicrobiales | Beijerinckiaceae |
| gi\|WP_166952180.1 | Pseudochelatococcus lubricantis | Alphaproteobacteria | Hyphomicrobiales | Beijerinckiaceae |
| gi\|AMA58454.1 | Bradyrhizobium sp. CCGE-LA001 | Alphaproteobacteria | Hyphomicrobiales | Bradyrhizobiaceae |
| gi\|WP_024341457.1 | Bradyrhizobium | Alphaproteobacteria | Hyphomicrobiales | Bradyrhizobiaceae |
| gi\|WP_146985765.1 | Bradyrhizobium macuxiense | Alphaproteobacteria | Hyphomicrobiales | Bradyrhizobiaceae |
| gi\|WP_079446668.1 | Nitrobacter vulgaris | Alphaproteobacteria | Hyphomicrobiales | Bradyrhizobiaceae |
| gi\|WP_011510671.1 | Nitrobacter hamburgensis | Alphaproteobacteria | Hyphomicrobiales | Bradyrhizobiaceae |
| gi\|KJC59346.1 | Bradyrhizobium sp. LTSPM299 | Alphaproteobacteria | Hyphomicrobiales | Bradyrhizobiaceae |
| gi\|WP_041748274.1 | Bradyrhizobium cosmicum | Alphaproteobacteria | Hyphomicrobiales | Bradyrhizobiaceae |
| gi\|WP_011316081.1 | Nitrobacter winogradskyi | Alphaproteobacteria | Hyphomicrobiales | Bradyrhizobiaceae |
| gi\|WP_122403025.1 | Bradyrhizobium vignae | Alphaproteobacteria | Hyphomicrobiales | Bradyrhizobiaceae |
| gi\|WP_146686543.1 | Bradyrhizobium canariense | Alphaproteobacteria | Hyphomicrobiales | Bradyrhizobiaceae |
| gi\|WP_057849643.1 | Bradyrhizobium valentinum | Alphaproteobacteria | Hyphomicrobiales | Bradyrhizobiaceae |
| gi\|WP_008968438.1 | Bradyrhizobium sp. STM 3843 | Alphaproteobacteria | Hyphomicrobiales | Bradyrhizobiaceae |
| gi\|AWM05971.1 | Bradyrhizobium symbiodeficiens | Alphaproteobacteria | Hyphomicrobiales | Bradyrhizobiaceae |
| gi\|WP_027559459.1 | Bradyrhizobium | Alphaproteobacteria | Hyphomicrobiales | Bradyrhizobiaceae |
| gi\|WP_079574113.1 | Bradyrhizobium erythrophlei | Alphaproteobacteria | Hyphomicrobiales | Bradyrhizobiaceae |
| gi\|WP_079603425.1 | Bradyrhizobium erythrophlei | Alphaproteobacteria | Hyphomicrobiales | Bradyrhizobiaceae |
| gi\|EAQ36134.1 | Nitrobacter sp. Nb-311A | Alphaproteobacteria | Hyphomicrobiales | Bradyrhizobiaceae |
| gi\|WP_139483879.1 | Bradyrhizobium ivorense | Alphaproteobacteria | Hyphomicrobiales | Bradyrhizobiaceae |
| gi\|ANW01924.1 | Bradyrhizobium icense | Alphaproteobacteria | Hyphomicrobiales | Bradyrhizobiaceae |
| gi\|WP_029584485.1 | Bradyrhizobium sp. URHD0069 | Alphaproteobacteria | Hyphomicrobiales | Bradyrhizobiaceae |
| gi\|WP_011512080.1 | Nitrobacter hamburgensis | Alphaproteobacteria | Hyphomicrobiales | Bradyrhizobiaceae |
| gi\|ABA05571.1 | Nitrobacter winogradskyi Nb-255 | Alphaproteobacteria | Hyphomicrobiales | Bradyrhizobiaceae |
| gi\|WP_074276509.1 | Bradyrhizobium erythrophlei | Alphaproteobacteria | Hyphomicrobiales | Bradyrhizobiaceae |
| gi\|WP_161856703.1 | Bradyrhizobium sp. CCBAU 051011 | Alphaproteobacteria | Hyphomicrobiales | Bradyrhizobiaceae |
| gi\|WP_194454045.1 | Bradyrhizobium sp. CCBAU 53421 | Alphaproteobacteria | Hyphomicrobiales | Bradyrhizobiaceae |
| gi\|WP_128953352.1 | Bradyrhizobium guangzhouense | Alphaproteobacteria | Hyphomicrobiales | Bradyrhizobiaceae |
| gi\|WP_084293192.1 | Bradyrhizobium sp. WSM3983 | Alphaproteobacteria | Hyphomicrobiales | Bradyrhizobiaceae |
| gi\|WP_009340768.1 | Afipia sp. 1NLS2 | Alphaproteobacteria | Hyphomicrobiales | Bradyrhizobiaceae |

| | | | | |
|---|---|---|---|---|
| gi\|WP_195788374.1 | Bradyrhizobium genosp. L | Alphaproteobacteria | Hyphomicrobiales | Bradyrhizobiaceae |
| gi\|WP_122404083.1 | Bradyrhizobium vignae | Alphaproteobacteria | Hyphomicrobiales | Bradyrhizobiaceae |
| gi\|WP_140977060.1 | Bradyrhizobium guangdongense | Alphaproteobacteria | Hyphomicrobiales | Bradyrhizobiaceae |
| gi\|WP_184255499.1 | Rhodopseudomonas rhenobacensis | Alphaproteobacteria | Hyphomicrobiales | Bradyrhizobiaceae |
| gi\|WP_166817957.1 | Bradyrhizobium sp. 1(2017) | Alphaproteobacteria | Hyphomicrobiales | Bradyrhizobiaceae |
| gi\|WP_110784444.1 | Rhodopseudomonas palustris | Alphaproteobacteria | Hyphomicrobiales | Bradyrhizobiaceae |
| gi\|WP_008968433.1 | Bradyrhizobium sp. STM 3843 | Alphaproteobacteria | Hyphomicrobiales | Bradyrhizobiaceae |
| gi\|WP_048758142.1 | Afipia felis | Alphaproteobacteria | Hyphomicrobiales | Bradyrhizobiaceae |
| gi\|WP_079545913.1 | Bradyrhizobium lablabi | Alphaproteobacteria | Hyphomicrobiales | Bradyrhizobiaceae |
| gi\|WP_092229576.1 | Bradyrhizobium sp. Gha | Alphaproteobacteria | Hyphomicrobiales | Bradyrhizobiaceae |
| gi\|WP_027537266.1 | Bradyrhizobium sp. URHA0002 | Alphaproteobacteria | Hyphomicrobiales | Bradyrhizobiaceae |
| gi\|WP_035961274.1 | Bradyrhizobium sp. URHA0013 | Alphaproteobacteria | Hyphomicrobiales | Bradyrhizobiaceae |
| gi\|WP_027522816.1 | Bradyrhizobium sp. Ec3.3 | Alphaproteobacteria | Hyphomicrobiales | Bradyrhizobiaceae |
| gi\|WP_024506901.1 | Bradyrhizobium sp. ARR65 | Alphaproteobacteria | Hyphomicrobiales | Bradyrhizobiaceae |
| gi\|WP_024518720.1 | Bradyrhizobium sp. Tv2a-2 | Alphaproteobacteria | Hyphomicrobiales | Bradyrhizobiaceae |
| gi\|QOZ45217.1 | Bradyrhizobium sp. CCBAU 53340 | Alphaproteobacteria | Hyphomicrobiales | Bradyrhizobiaceae |
| gi\|TYO62278.1 | Bradyrhizobium hipponense | Alphaproteobacteria | Hyphomicrobiales | Bradyrhizobiaceae |
| gi\|WP_074825245.1 | Bradyrhizobium | Alphaproteobacteria | Hyphomicrobiales | Bradyrhizobiaceae |
| gi\|WP_084807138.1 | Bradyrhizobium sp. NAS80.1 | Alphaproteobacteria | Hyphomicrobiales | Bradyrhizobiaceae |
| gi\|WP_130579395.1 | Bradyrhizobium sp. Leo170 | Alphaproteobacteria | Hyphomicrobiales | Bradyrhizobiaceae |
| gi\|WP_024515210.1 | Bradyrhizobium sp. Tv2a-2 | Alphaproteobacteria | Hyphomicrobiales | Bradyrhizobiaceae |
| gi\|WP_054160643.1 | Rhodopseudomonas sp. AAP120 | Alphaproteobacteria | Hyphomicrobiales | Bradyrhizobiaceae |
| gi\|WP_027529225.1 | Bradyrhizobium sp. WSM3983 | Alphaproteobacteria | Hyphomicrobiales | Bradyrhizobiaceae |
| gi\|AUC98923.1 | Bradyrhizobium sp. SK17 | Alphaproteobacteria | Hyphomicrobiales | Bradyrhizobiaceae |
| gi\|WP_184087442.1 | Afipia massiliensis | Alphaproteobacteria | Hyphomicrobiales | Bradyrhizobiaceae |
| gi\|WP_024506899.1 | Bradyrhizobium sp. ARR65 | Alphaproteobacteria | Hyphomicrobiales | Bradyrhizobiaceae |
| gi\|WP_022724158.1 | Rhodopseudomonas sp. B29 | Alphaproteobacteria | Hyphomicrobiales | Bradyrhizobiaceae |
| gi\|WP_130579972.1 | Bradyrhizobium sp. Leo170 | Alphaproteobacteria | Hyphomicrobiales | Bradyrhizobiaceae |
| gi\|WP_013913159.1 | Afipia carboxidovorans | Alphaproteobacteria | Hyphomicrobiales | Bradyrhizobiaceae |
| gi\|WP_177248173.1 | Bradyrhizobium sp. Ghvi | Alphaproteobacteria | Hyphomicrobiales | Bradyrhizobiaceae |
| gi\|WP_079447150.1 | Nitrobacter vulgaris | Alphaproteobacteria | Hyphomicrobiales | Bradyrhizobiaceae |
| gi\|WP_054359698.1 | Prosthecomicrobium hirschii | Alphaproteobacteria | Hyphomicrobiales | Hyphomicrobiaceae |
| gi\|WP_013214625.1 | Hyphomicrobium denitrificans | Alphaproteobacteria | Hyphomicrobiales | Hyphomicrobiaceae |
| gi\|WP_046477349.1 | Candidatus Filomicrobium marinum | Alphaproteobacteria | Hyphomicrobiales | Hyphomicrobiaceae |
| gi\|WP_170936960.1 | Rhodomicrobium | Alphaproteobacteria | Hyphomicrobiales | Hyphomicrobiaceae |
| gi\|WP_183851922.1 | Prosthecomicrobium pneumaticum | Alphaproteobacteria | Hyphomicrobiales | Hyphomicrobiaceae |
| gi\|WP_069096256.1 | Methyloligella halotolerans | Alphaproteobacteria | Hyphomicrobiales | Hyphomicrobiaceae |
| gi\|ACA18979.1 | Methylobacterium sp. 4-46 | Alphaproteobacteria | Hyphomicrobiales | Methylobacteriaceae |
| gi\|WP_165055868.1 | Methylocystis sp. MJC1 | Alphaproteobacteria | Hyphomicrobiales | Methylocystaceae |
| gi\|QCI66377.1 | Phreatobacter stygius | Alphaproteobacteria | Hyphomicrobiales | Phreatobacteraceae |
| gi\|WP_047506981.1 | Rhizobium sp. YR528 | Alphaproteobacteria | Hyphomicrobiales | Rhizobiaceae |
| gi\|WP_165933165.1 | Rhizobium sp. BK068 | Alphaproteobacteria | Hyphomicrobiales | Rhizobiaceae |
| gi\|PLX34769.1 | Hyphomicrobiales bacterium | Alphaproteobacteria | Hyphomicrobiales | |

| | | | | |
|---|---|---|---|---|
| gi\|PLX44011.1 | Hyphomicrobiales bacterium | Alphaproteobacteria | Hyphomicrobiales | |
| gi\|WP_013299735.1 | Parvularcula bermudensis | Alphaproteobacteria | Parvularculales | Parvularculaceae |
| gi\|PDT56766.1 | Bradyrhizobium diazoefficiens | Alphaproteobacteria | Rhizobiales | Bradyrhizobiaceae |
| gi\|RTM12335.1 | Bradyrhizobiaceae bacterium | Alphaproteobacteria | Rhizobiales | Bradyrhizobiaceae |
| gi\|WP_128956486.1 | Bradyrhizobium zhanjiangense | Alphaproteobacteria | Rhizobiales | Bradyrhizobiaceae |
| gi\|PIT04588.1 | Bradyrhizobium nitroreducens | Alphaproteobacteria | Rhizobiales | Bradyrhizobiaceae |
| gi\|RTL54519.1 | Bradyrhizobiaceae bacterium | Alphaproteobacteria | Rhizobiales | Bradyrhizobiaceae |
| gi\|OJV02895.1 | Nitrobacter sp. 62-23 | Alphaproteobacteria | Rhizobiales | Bradyrhizobiaceae |
| gi\|OJW62195.1 | Afipia sp. 64-13 | Alphaproteobacteria | Rhizobiales | Bradyrhizobiaceae |
| gi\|SIO52686.1 | Bradyrhizobium erythrophlei | Alphaproteobacteria | Rhizobiales | Bradyrhizobiaceae |
| gi\|PJG51612.1 | Bradyrhizobium forestalis | Alphaproteobacteria | Rhizobiales | Bradyrhizobiaceae |
| gi\|TWC05401.1 | Bradyrhizobium macuxiense | Alphaproteobacteria | Rhizobiales | Bradyrhizobiaceae |
| gi\|PJJ13682.1 | Bradyrhizobium lablabi | Alphaproteobacteria | Rhizobiales | Bradyrhizobiaceae |
| gi\|TXN21254.1 | Methylobacterium sp. WL9 | Alphaproteobacteria | Rhizobiales | Methylobacteriaceae |
| gi\|PPD41678.1 | Methylocystis sp. | Alphaproteobacteria | Rhizobiales | Methylocystaceae |
| gi\|TXI11742.1 | Rhizobium sp. | Alphaproteobacteria | Rhizobiales | Rhizobiaceae |
| gi\|OJY10277.1 | Rhizobiales bacterium 62-47 | Alphaproteobacteria | Rhizobiales | |
| gi\|WP_165978909.1 | Antarcticimicrobium luteum | Alphaproteobacteria | Rhodobacterales | Rhodobacteraceae |
| gi\|WP_167601659.1 | Celeribacter sp. HF31 | Alphaproteobacteria | Rhodobacterales | Roseobacteraceae |
| gi\|SFI48938.1 | Celeribacter neptunius | Alphaproteobacteria | Rhodobacterales | Roseobacteraceae |
| gi\|TYC49223.1 | Rhodobacterales bacterium | Alphaproteobacteria | Rhodobacterales | |
| gi\|GCD52509.1 | Acetobacter pasteurianus NBRC 3188 | Alphaproteobacteria | Rhodospirillales | Acetobacteraceae |
| gi\|WP_194255797.1 | Gluconobacter cerevisiae | Alphaproteobacteria | Rhodospirillales | Acetobacteraceae |
| gi\|GFE93475.1 | Acetobacter persici | Alphaproteobacteria | Rhodospirillales | Acetobacteraceae |
| gi\|WP_146795768.1 | Gluconobacter wancherniae | Alphaproteobacteria | Rhodospirillales | Acetobacteraceae |
| gi\|WP_146887991.1 | Acetobacter oeni | Alphaproteobacteria | Rhodospirillales | Acetobacteraceae |
| gi\|WP_186772809.1 | Siccirubricoccus deserti | Alphaproteobacteria | Rhodospirillales | Acetobacteraceae |
| gi\|WP_119781869.1 | Oleomonas sp. K1W22B-8 | Alphaproteobacteria | Rhodospirillales | Acetobacteraceae |
| gi\|WP_182979855.1 | Gluconacetobacter asukensis | Alphaproteobacteria | Rhodospirillales | Acetobacteraceae |
| gi\|PAK77424.1 | Acetobacter fabarum | Alphaproteobacteria | Rhodospirillales | Acetobacteraceae |
| gi\|WP_168046515.1 | Roseomonas frigidaquae | Alphaproteobacteria | Rhodospirillales | Acetobacteraceae |
| gi\|WP_095351963.1 | Acetobacter syzygii | Alphaproteobacteria | Rhodospirillales | Acetobacteraceae |
| gi\|KXV20849.1 | Gluconobacter japonicus | Alphaproteobacteria | Rhodospirillales | Acetobacteraceae |
| gi\|TCZ64361.1 | Paracraurococcus sp. NE82 | Alphaproteobacteria | Rhodospirillales | Acetobacteraceae |
| gi\|WP_119832995.1 | Azospirillum sp. K2W22B-5 | Alphaproteobacteria | Rhodospirillales | Azospirillaceae |
| gi\|ALG73010.1 | Azospirillum thiophilum | Alphaproteobacteria | Rhodospirillales | Azospirillaceae |
| gi\|ALG72630.1 | Azospirillum thiophilum | Alphaproteobacteria | Rhodospirillales | Azospirillaceae |
| gi\|WP_149224261.1 | Azospirillum sp. B21 | Alphaproteobacteria | Rhodospirillales | Azospirillaceae |
| gi\|WP_119833109.1 | Azospirillum sp. K2W22B-5 | Alphaproteobacteria | Rhodospirillales | Azospirillaceae |
| gi\|WP_109120916.1 | Azospirillum sp. TSO22-1 | Alphaproteobacteria | Rhodospirillales | Azospirillaceae |
| gi\|WP_085940773.1 | Azospirillum sp. B506 | Alphaproteobacteria | Rhodospirillales | Azospirillaceae |
| gi\|WP_109154981.1 | Azospirillum sp. TSO5 | Alphaproteobacteria | Rhodospirillales | Azospirillaceae |
| gi\|QCG98360.1 | Azospirillum sp. TSA2s | Alphaproteobacteria | Rhodospirillales | Azospirillaceae |

| | | | | |
|---|---|---|---|---|
| gi\|WP_108546737.1 | Azospirillum humicireducens | Alphaproteobacteria | Rhodospirillales | Azospirillaceae |
| gi\|WP_109105952.1 | Azospirillum sp. TSO35-2 | Alphaproteobacteria | Rhodospirillales | Azospirillaceae |
| gi\|WP_109118359.1 | Azospirillum sp. TSO22-1 | Alphaproteobacteria | Rhodospirillales | Azospirillaceae |
| gi\|WP_042704792.1 | Azospirillum sp. B506 | Alphaproteobacteria | Rhodospirillales | Azospirillaceae |
| gi\|WP_029008038.1 | Azospirillum halopraeferens | Alphaproteobacteria | Rhodospirillales | Azospirillaceae |
| gi\|WP_098737002.1 | Azospirillum palustre | Alphaproteobacteria | Rhodospirillales | Azospirillaceae |
| gi\|WP_126997618.1 | Azospirillum doebereinerae | Alphaproteobacteria | Rhodospirillales | Azospirillaceae |
| gi\|WP_126615361.1 | Azospirillum griseum | Alphaproteobacteria | Rhodospirillales | Azospirillaceae |
| gi\|WP_109118358.1 | Azospirillum sp. TSO22-1 | Alphaproteobacteria | Rhodospirillales | Azospirillaceae |
| gi\|WP_114861889.1 | Azospirillum brasilense | Alphaproteobacteria | Rhodospirillales | Azospirillaceae |
| gi\|WP_029012946.1 | Niveispirillum irakense | Alphaproteobacteria | Rhodospirillales | Azospirillaceae |
| gi\|WP_180281532.1 | Azospirillum oleiclasticum | Alphaproteobacteria | Rhodospirillales | Azospirillaceae |
| gi\|WP_098737664.1 | Azospirillum palustre | Alphaproteobacteria | Rhodospirillales | Azospirillaceae |
| gi\|WP_160106109.1 | unclassified Azospirillum | Alphaproteobacteria | Rhodospirillales | Azospirillaceae |
| gi\|WP_169789456.1 | Skermanella aerolata | Alphaproteobacteria | Rhodospirillales | Azospirillaceae |
| gi\|ALG73012.1 | Azospirillum thiophilum | Alphaproteobacteria | Rhodospirillales | Azospirillaceae |
| gi\|WP_045583148.1 | Azospirillum thiophilum | Alphaproteobacteria | Rhodospirillales | Azospirillaceae |
| gi\|ALJ38878.1 | Azospirillum brasilense | Alphaproteobacteria | Rhodospirillales | Azospirillaceae |
| gi\|WP_174473933.1 | Azospirillum melinis | Alphaproteobacteria | Rhodospirillales | Azospirillaceae |
| gi\|WP_108548598.1 | Azospirillum humicireducens | Alphaproteobacteria | Rhodospirillales | Azospirillaceae |
| gi\|BAI74109.1 | Azospirillum sp. B510 | Alphaproteobacteria | Rhodospirillales | Azospirillaceae |
| gi\|WP_052293713.1 | Azospirillum sp. B510 | Alphaproteobacteria | Rhodospirillales | Azospirillaceae |
| gi\|WP_173981495.1 | Magnetospirillum sp. SS-4 | Alphaproteobacteria | Rhodospirillales | Rhodospirillaceae |
| gi\|WP_176525116.1 | Caenispirillum bisanense | Alphaproteobacteria | Rhodospirillales | Rhodospirillaceae |
| gi\|WP_096703471.1 | Magnetospirillum sp. 15-1 | Alphaproteobacteria | Rhodospirillales | Rhodospirillaceae |
| gi\|PWC80091.1 | Azospirillum sp. TSH64 | Alphaproteobacteria | Rhodospirillales | Rhodospirillaceae |
| gi\|TWA82718.1 | Azospirillum brasilense | Alphaproteobacteria | Rhodospirillales | Rhodospirillaceae |
| gi\|WP_011383878.1 | Magnetospirillum magneticum | Alphaproteobacteria | Rhodospirillales | Rhodospirillaceae |
| gi\|WP_184264826.1 | Novispirillum itersonii | Alphaproteobacteria | Rhodospirillales | Rhodospirillaceae |
| gi\|WP_155976174.1 | Novispirillum itersonii | Alphaproteobacteria | Rhodospirillales | Rhodospirillaceae |
| gi\|WP_188579298.1 | Tistrella bauzanensis | Alphaproteobacteria | Rhodospirillales | Rhodospirillaceae |
| gi\|WP_024082103.1 | Magnetospirillum gryphiswaldense | Alphaproteobacteria | Rhodospirillales | Rhodospirillaceae |
| gi\|WP_085935705.1 | Enhydrobacter aerosaccus | Alphaproteobacteria | Rhodospirillales | Rhodospirillales incerta sedis |
| gi\|WP_114129871.1 | Thalassospira | Alphaproteobacteria | Rhodospirillales | Thalassospiraceae |
| gi\|WP_175581271.1 | Thalassospira sp. HF15 | Alphaproteobacteria | Rhodospirillales | Thalassospiraceae |
| gi\|TAN58312.1 | Rhodospirillales bacterium | Alphaproteobacteria | Rhodospirillales | |
| gi\|TAN56822.1 | Rhodospirillales bacterium | Alphaproteobacteria | Rhodospirillales | |
| gi\|WP_028641260.1 | Novosphingobium acidiphilum | Alphaproteobacteria | Sphingomonadales | Sphingomonadaceae |
| gi\|WP_072383391.1 | Novosphingobium sp. NDB2Meth1 | Alphaproteobacteria | Sphingomonadales | Sphingomonadaceae |
| gi\|WP_154651274.1 | Sphingomonas echinoides | Alphaproteobacteria | Sphingomonadales | Sphingomonadaceae |
| gi\|WP_159760003.1 | Sphingomonas sp. 8AM | Alphaproteobacteria | Sphingomonadales | Sphingomonadaceae |
| gi\|WP_008069953.1 | Novosphingobium nitrogenifigens | Alphaproteobacteria | Sphingomonadales | Sphingomonadaceae |
| gi\|WP_130154896.1 | Sphingomonas populi | Alphaproteobacteria | Sphingomonadales | Sphingomonadaceae |

| | | | | |
|---|---|---|---|---|
| gi\|OYX65415.1 | Sphingomonadales bacterium 32-64-17 | Alphaproteobacteria | Sphingomonadales | unclassified Sphingomo |
| gi\|TMK48487.1 | Alphaproteobacteria bacterium | Alphaproteobacteria | | |
| gi\|TMK11737.1 | Alphaproteobacteria bacterium | Alphaproteobacteria | | |
| gi\|TMK09383.1 | Alphaproteobacteria bacterium | Alphaproteobacteria | | |
| gi\|TMJ58051.1 | Alphaproteobacteria bacterium | Alphaproteobacteria | | |
| gi\|RMF37420.1 | Alphaproteobacteria bacterium | Alphaproteobacteria | | |
| gi\|PCH99619.1 | Alphaproteobacteria bacterium | Alphaproteobacteria | | |
| gi\|WP_033460812.1 | Bordetella | Betaproteobacteria | Burkholderiales | Alcaligenaceae |
| gi\|WP_085316148.1 | Derxia lacustris | Betaproteobacteria | Burkholderiales | Alcaligenaceae |
| gi\|WP_088602348.1 | Candidimonas nitroreducens | Betaproteobacteria | Burkholderiales | Alcaligenaceae |
| gi\|WP_124080538.1 | Pigmentiphaga humi | Betaproteobacteria | Burkholderiales | Alcaligenaceae |
| gi\|WP_057284188.1 | Achromobacter sp. Root83 | Betaproteobacteria | Burkholderiales | Alcaligenaceae |
| gi\|WP_175212489.1 | Achromobacter aegrifaciens | Betaproteobacteria | Burkholderiales | Alcaligenaceae |
| gi\|WP_084025253.1 | Bordetella flabilis | Betaproteobacteria | Burkholderiales | Alcaligenaceae |
| gi\|WP_086056058.1 | Bordetella genomosp. 9 | Betaproteobacteria | Burkholderiales | Alcaligenaceae |
| gi\|WP_094840978.1 | Bordetella genomosp. 11 | Betaproteobacteria | Burkholderiales | Alcaligenaceae |
| gi\|WP_183006757.1 | Achromobacter sp. UMC71 | Betaproteobacteria | Burkholderiales | Alcaligenaceae |
| gi\|WP_028312171.1 | Derxia gummosa | Betaproteobacteria | Burkholderiales | Alcaligenaceae |
| gi\|WP_132583185.1 | Paralcaligenes ureilyticus | Betaproteobacteria | Burkholderiales | Alcaligenaceae |
| gi\|WP_176256873.1 | Derxia lacustris | Betaproteobacteria | Burkholderiales | Alcaligenaceae |
| gi\|WP_043544448.1 | Achromobacter sp. RTa | Betaproteobacteria | Burkholderiales | Alcaligenaceae |
| gi\|WP_073101061.1 | Candidimonas bauzanensis | Betaproteobacteria | Burkholderiales | Alcaligenaceae |
| gi\|WP_028311324.1 | Derxia gummosa | Betaproteobacteria | Burkholderiales | Alcaligenaceae |
| gi\|WP_024004599.1 | Advenella kashmirensis | Betaproteobacteria | Burkholderiales | Alcaligenaceae |
| gi\|WP_087840052.1 | unclassified Pigmentiphaga | Betaproteobacteria | Burkholderiales | Alcaligenaceae |
| gi\|WP_155257660.1 | Achromobacter xylosoxidans | Betaproteobacteria | Burkholderiales | Alcaligenaceae |
| gi\|WP_175127481.1 | Achromobacter piechaudii | Betaproteobacteria | Burkholderiales | Alcaligenaceae |
| gi\|TFL13425.1 | Pusillimonas caeni | Betaproteobacteria | Burkholderiales | Alcaligenaceae |
| gi\|WP_086063965.1 | Bordetella genomosp. 8 | Betaproteobacteria | Burkholderiales | Alcaligenaceae |
| gi\|WP_103275940.1 | Achromobacter sp. AONIH1 | Betaproteobacteria | Burkholderiales | Alcaligenaceae |
| gi\|WP_175168876.1 | Achromobacter kerstersii | Betaproteobacteria | Burkholderiales | Alcaligenaceae |
| gi\|WP_054422616.1 | Achromobacter kerstersii | Betaproteobacteria | Burkholderiales | Alcaligenaceae |
| gi\|WP_073101629.1 | Candidimonas bauzanensis | Betaproteobacteria | Burkholderiales | Alcaligenaceae |
| gi\|WP_083812502.1 | Pusillimonas sp. T7-7 | Betaproteobacteria | Burkholderiales | Alcaligenaceae |
| gi\|WP_163652592.1 | Orrella sp. NBD-18 | Betaproteobacteria | Burkholderiales | Alcaligenaceae |
| gi\|WP_083228676.1 | Bordetella sp. H567 | Betaproteobacteria | Burkholderiales | Alcaligenaceae |
| gi\|WP_128353649.1 | Pusillimonas thiosulfatoxidans | Betaproteobacteria | Burkholderiales | Alcaligenaceae |
| gi\|RCS59432.1 | Parvibium lacunae | Betaproteobacteria | Burkholderiales | Alcaligenaceae |
| gi\|WP_100694585.1 | Advenella sp. S44 | Betaproteobacteria | Burkholderiales | Alcaligenaceae |
| gi\|PVY68095.1 | Pusillimonas noertemannii | Betaproteobacteria | Burkholderiales | Alcaligenaceae |
| gi\|KAB0616721.1 | Castellaniella defragrans | Betaproteobacteria | Burkholderiales | Alcaligenaceae |
| gi\|AWB32483.1 | Algicoccus marinus | Betaproteobacteria | Burkholderiales | Alcaligenaceae |
| gi\|WP_102075276.1 | Pusillimonas sp. JR1/69-3-13 | Betaproteobacteria | Burkholderiales | Alcaligenaceae |

| | | | | |
|---|---|---|---|---|
| gi\|WP_102070847.1 | Pusillimonas sp. JR1/69-2-13 | Betaproteobacteria | Burkholderiales | Alcaligenaceae |
| gi\|WP_126708694.1 | Candidimonas sp. SYP-B2681 | Betaproteobacteria | Burkholderiales | Alcaligenaceae |
| gi\|WP_054445704.1 | Achromobacter xylosoxidans | Betaproteobacteria | Burkholderiales | Alcaligenaceae |
| gi\|WP_148304924.1 | Castellaniella defragrans | Betaproteobacteria | Burkholderiales | Alcaligenaceae |
| gi\|WP_019937105.1 | Bordetella sp. FB-8 | Betaproteobacteria | Burkholderiales | Alcaligenaceae |
| gi\|WP_081247757.1 | Achromobacter xylosoxidans | Betaproteobacteria | Burkholderiales | Alcaligenaceae |
| gi\|WP_175191147.1 | Achromobacter deleyi | Betaproteobacteria | Burkholderiales | Alcaligenaceae |
| gi\|WP_175141704.1 | Achromobacter pulmonis | Betaproteobacteria | Burkholderiales | Alcaligenaceae |
| gi\|WP_160977083.1 | Pusillimonas sp. TS35 | Betaproteobacteria | Burkholderiales | Alcaligenaceae |
| gi\|WP_006391004.1 | Achromobacter insuavis | Betaproteobacteria | Burkholderiales | Alcaligenaceae |
| gi\|WP_057284210.1 | Achromobacter sp. Root83 | Betaproteobacteria | Burkholderiales | Alcaligenaceae |
| gi\|QDQ86281.1 | Alcaligenaceae bacterium SJ-26 | Betaproteobacteria | Burkholderiales | Alcaligenaceae |
| gi\|WP_176090837.1 | Achromobacter anxifer | Betaproteobacteria | Burkholderiales | Alcaligenaceae |
| gi\|WP_169272010.1 | Achromobacter sp. Bel | Betaproteobacteria | Burkholderiales | Alcaligenaceae |
| gi\|WP_066132827.1 | Bordetella ansorpii | Betaproteobacteria | Burkholderiales | Alcaligenaceae |
| gi\|WP_132477078.1 | Paracandidimonas soli | Betaproteobacteria | Burkholderiales | Alcaligenaceae |
| gi\|WP_008167326.1 | Achromobacter arsenitoxydans | Betaproteobacteria | Burkholderiales | Alcaligenaceae |
| gi\|WP_165026533.1 | Parapusillimonas sp. SGNA-6 | Betaproteobacteria | Burkholderiales | Alcaligenaceae |
| gi\|WP_100854733.1 | Achromobacter spanius | Betaproteobacteria | Burkholderiales | Alcaligenaceae |
| gi\|SPT42347.1 | Achromobacter denitrificans | Betaproteobacteria | Burkholderiales | Alcaligenaceae |
| gi\|WP_086066183.1 | Bordetella genomosp. 8 | Betaproteobacteria | Burkholderiales | Alcaligenaceae |
| gi\|WP_073108033.1 | Candidimonas bauzanensis | Betaproteobacteria | Burkholderiales | Alcaligenaceae |
| gi\|WP_006221317.1 | Achromobacter piechaudii | Betaproteobacteria | Burkholderiales | Alcaligenaceae |
| gi\|WP_175200246.1 | Achromobacter insolitus | Betaproteobacteria | Burkholderiales | Alcaligenaceae |
| gi\|WP_175168878.1 | Achromobacter kerstersii | Betaproteobacteria | Burkholderiales | Alcaligenaceae |
| gi\|OXR48108.1 | Pusillimonas sp. T2 | Betaproteobacteria | Burkholderiales | Alcaligenaceae |
| gi\|WP_025138684.1 | Achromobacter sp. DH1f | Betaproteobacteria | Burkholderiales | Alcaligenaceae |
| gi\|WP_074046797.1 | Orrella dioscoreae | Betaproteobacteria | Burkholderiales | Alcaligenaceae |
| gi\|WP_046805166.1 | Achromobacter sp. LC458 | Betaproteobacteria | Burkholderiales | Alcaligenaceae |
| gi\|WP_088155741.1 | Achromobacter xylosoxidans | Betaproteobacteria | Burkholderiales | Alcaligenaceae |
| gi\|ARP95987.1 | Bordetella genomosp. 13 | Betaproteobacteria | Burkholderiales | Alcaligenaceae |
| gi\|WP_166409787.1 | Paenalcaligenes suwonensis | Betaproteobacteria | Burkholderiales | Alcaligenaceae |
| gi\|WP_102773768.1 | Achromobacter | Betaproteobacteria | Burkholderiales | Alcaligenaceae |
| gi\|RZS81195.1 | Pigmentiphaga kullae | Betaproteobacteria | Burkholderiales | Alcaligenaceae |
| gi\|WP_012250095.1 | Bordetella petrii | Betaproteobacteria | Burkholderiales | Alcaligenaceae |
| gi\|WP_175173024.1 | Achromobacter pestifer | Betaproteobacteria | Burkholderiales | Alcaligenaceae |
| gi\|WP_175168866.1 | Achromobacter kerstersii | Betaproteobacteria | Burkholderiales | Alcaligenaceae |
| gi\|WP_054439856.1 | Achromobacter xylosoxidans | Betaproteobacteria | Burkholderiales | Alcaligenaceae |
| gi\|WP_129245627.1 | Achromobacter veterisilvae | Betaproteobacteria | Burkholderiales | Alcaligenaceae |
| gi\|WP_133606869.1 | Aquabacterium commune | Betaproteobacteria | Burkholderiales | Aquabacterium |
| gi\|WP_058087620.1 | Aquabacterium parvum | Betaproteobacteria | Burkholderiales | Aquabacterium |
| gi\|WP_173122712.1 | Aquabacterium terrae | Betaproteobacteria | Burkholderiales | Aquabacterium |
| gi\|WP_166834419.1 | Aquabacterium sp. A08 | Betaproteobacteria | Burkholderiales | Aquabacterium |

| | | | | |
|---|---|---|---|---|
| gi\|WP_035038245.1 | Aquabacterium sp. NJ1 | Betaproteobacteria | Burkholderiales | Aquabacterium |
| gi\|TAK86256.1 | Aquabacterium sp. | Betaproteobacteria | Burkholderiales | Aquabacterium |
| gi\|WP_109035138.1 | Aquabacterium olei | Betaproteobacteria | Burkholderiales | Aquabacterium |
| gi\|WP_166832290.1 | Aquabacterium sp. A08 | Betaproteobacteria | Burkholderiales | Aquabacterium |
| gi\|TXJ02928.1 | Aquabacterium sp. | Betaproteobacteria | Burkholderiales | Aquabacterium |
| gi\|WP_161648904.1 | Aquabacterium fontiphilum | Betaproteobacteria | Burkholderiales | Aquabacterium |
| gi\|TBO31172.1 | Aquabacterium lacunae | Betaproteobacteria | Burkholderiales | Aquabacterium |
| gi\|TAK95144.1 | Aquabacterium sp. | Betaproteobacteria | Burkholderiales | Aquabacterium |
| gi\|WP_052736015.1 | Aquincola tertiaricarbonis | Betaproteobacteria | Burkholderiales | Aquincola |
| gi\|WP_128226675.1 | Aquincola rivuli | Betaproteobacteria | Burkholderiales | Aquincola |
| gi\|WP_149670301.1 | Paraburkholderia panacisoli | Betaproteobacteria | Burkholderiales | Burkholderiaceae |
| gi\|WP_133664538.1 | Paraburkholderia sp. BL10I2N1 | Betaproteobacteria | Burkholderiales | Burkholderiaceae |
| gi\|TAL96366.1 | Paraburkholderia sp. | Betaproteobacteria | Burkholderiales | Burkholderiaceae |
| gi\|WP_115779937.1 | Paraburkholderia caffeinilytica | Betaproteobacteria | Burkholderiales | Burkholderiaceae |
| gi\|TCK95019.1 | Paraburkholderia sp. BL9I2N2 | Betaproteobacteria | Burkholderiales | Burkholderiaceae |
| gi\|WP_063963604.1 | Caballeronia hypogeia | Betaproteobacteria | Burkholderiales | Burkholderiaceae |
| gi\|WP_064267253.1 | Paraburkholderia ginsengiterrae | Betaproteobacteria | Burkholderiales | Burkholderiaceae |
| gi\|WP_167282776.1 | Paraburkholderia sp. Cy-641 | Betaproteobacteria | Burkholderiales | Burkholderiaceae |
| gi\|WP_074764086.1 | Paraburkholderia fungorum | Betaproteobacteria | Burkholderiales | Burkholderiaceae |
| gi\|WP_082855023.1 | Paraburkholderia phytofirmans | Betaproteobacteria | Burkholderiales | Burkholderiaceae |
| gi\|WP_087646703.1 | Caballeronia choica | Betaproteobacteria | Burkholderiales | Burkholderiaceae |
| gi\|WP_087740094.1 | Paraburkholderia piptadeniae | Betaproteobacteria | Burkholderiales | Burkholderiaceae |
| gi\|WP_087043320.1 | Caballeronia ptereochthonis | Betaproteobacteria | Burkholderiales | Burkholderiaceae |
| gi\|WP_031364116.1 | Caballeronia sordidicola | Betaproteobacteria | Burkholderiales | Burkholderiaceae |
| gi\|WP_054043008.1 | Paraburkholderia | Betaproteobacteria | Burkholderiales | Burkholderiaceae |
| gi\|WP_094779537.1 | Paraburkholderia ribeironis | Betaproteobacteria | Burkholderiales | Burkholderiaceae |
| gi\|WP_038715581.1 | Burkholderia sp. lig30 | Betaproteobacteria | Burkholderiales | Burkholderiaceae |
| gi\|WP_066484541.1 | Burkholderia sp. BDU8 | Betaproteobacteria | Burkholderiales | Burkholderiaceae |
| gi\|WP_109481296.1 | Paraburkholderia sp. C35 | Betaproteobacteria | Burkholderiales | Burkholderiaceae |
| gi\|WP_165089356.1 | Caballeronia sp. SBC1 | Betaproteobacteria | Burkholderiales | Burkholderiaceae |
| gi\|TXH50240.1 | Burkholderiaceae bacterium | Betaproteobacteria | Burkholderiales | Burkholderiaceae |
| gi\|WP_061149684.1 | Caballeronia arvi | Betaproteobacteria | Burkholderiales | Burkholderiaceae |
| gi\|WP_175843841.1 | Burkholderia arboris | Betaproteobacteria | Burkholderiales | Burkholderiaceae |
| gi\|WP_153141262.1 | Paraburkholderia agricolaris | Betaproteobacteria | Burkholderiales | Burkholderiaceae |
| gi\|WP_175108525.1 | Pararobbsia alpina | Betaproteobacteria | Burkholderiales | Burkholderiaceae |
| gi\|WP_063493256.1 | Caballeronia sordidicola | Betaproteobacteria | Burkholderiales | Burkholderiaceae |
| gi\|WP_175107896.1 | Pararobbsia alpina | Betaproteobacteria | Burkholderiales | Burkholderiaceae |
| gi\|WP_152765416.1 | Paraburkholderia franconis | Betaproteobacteria | Burkholderiales | Burkholderiaceae |
| gi\|WP_097219475.1 | Burkholderia sp. YR290 | Betaproteobacteria | Burkholderiales | Burkholderiaceae |
| gi\|WP_042323428.1 | Paraburkholderia ginsengisoli | Betaproteobacteria | Burkholderiales | Burkholderiaceae |
| gi\|WP_121277867.1 | Trinickia fusca | Betaproteobacteria | Burkholderiales | Burkholderiaceae |
| gi\|WP_062091398.1 | Caballeronia udeis | Betaproteobacteria | Burkholderiales | Burkholderiaceae |
| gi\|WP_111928837.1 | Paraburkholderia bryophila | Betaproteobacteria | Burkholderiales | Burkholderiaceae |

| | | | | |
|---|---|---|---|---|
| gi\|WP_129563908.1 | Paraburkholderia dokdonella | Betaproteobacteria | Burkholderiales | Burkholderiaceae |
| gi\|WP_121323327.1 | Paraburkholderia sp. RAU2J | Betaproteobacteria | Burkholderiales | Burkholderiaceae |
| gi\|WP_029308973.1 | Cupriavidus metallidurans | Betaproteobacteria | Burkholderiales | Burkholderiaceae |
| gi\|WP_038751126.1 | Burkholderia | Betaproteobacteria | Burkholderiales | Burkholderiaceae |
| gi\|WP_073430397.1 | Paraburkholderia terricola | Betaproteobacteria | Burkholderiales | Burkholderiaceae |
| gi\|WP_112003695.1 | Burkholderia sp. yr520 | Betaproteobacteria | Burkholderiales | Burkholderiaceae |
| gi\|PRX36805.1 | Paraburkholderia sp. BL18I3N2 | Betaproteobacteria | Burkholderiales | Burkholderiaceae |
| gi\|WP_122171810.1 | Caballeronia sordidicola | Betaproteobacteria | Burkholderiales | Burkholderiaceae |
| gi\|WP_137957152.1 | Burkholderia sp. 4M9327F10 | Betaproteobacteria | Burkholderiales | Burkholderiaceae |
| gi\|WP_175108439.1 | Pararobbsia alpina | Betaproteobacteria | Burkholderiales | Burkholderiaceae |
| gi\|WP_132259281.1 | Paucimonas lemoignei | Betaproteobacteria | Burkholderiales | Burkholderiaceae |
| gi\|WP_062085155.1 | Caballeronia udeis | Betaproteobacteria | Burkholderiales | Burkholderiaceae |
| gi\|WP_158935356.1 | Burkholderia sp. S171 | Betaproteobacteria | Burkholderiales | Burkholderiaceae |
| gi\|WP_091993030.1 | Paraburkholderia lycopersici | Betaproteobacteria | Burkholderiales | Burkholderiaceae |
| gi\|AFQ49881.1 | Burkholderia cepacia GG4 | Betaproteobacteria | Burkholderiales | Burkholderiaceae |
| gi\|WP_017235416.1 | Pandoraea sp. B-6 | Betaproteobacteria | Burkholderiales | Burkholderiaceae |
| gi\|RSL25588.1 | Caballeronia sordidicola | Betaproteobacteria | Burkholderiales | Burkholderiaceae |
| gi\|WP_110328650.1 | Paraburkholderia tropica | Betaproteobacteria | Burkholderiales | Burkholderiaceae |
| gi\|TAM04724.1 | Paraburkholderia sp. | Betaproteobacteria | Burkholderiales | Burkholderiaceae |
| gi\|WP_186067797.1 | Burkholderia gladioli | Betaproteobacteria | Burkholderiales | Burkholderiaceae |
| gi\|WP_169498939.1 | Paraburkholderia sp. G-4-1-8 | Betaproteobacteria | Burkholderiales | Burkholderiaceae |
| gi\|WP_152764217.1 | Paraburkholderia franconis | Betaproteobacteria | Burkholderiales | Burkholderiaceae |
| gi\|WP_150598652.1 | Pandoraea fibrosis | Betaproteobacteria | Burkholderiales | Burkholderiaceae |
| gi\|WP_175228245.1 | Paraburkholderia humisilvae | Betaproteobacteria | Burkholderiales | Burkholderiaceae |
| gi\|WP_027820611.1 | Paraburkholderia bannensis | Betaproteobacteria | Burkholderiales | Burkholderiaceae |
| gi\|WP_120343044.1 | Paraburkholderia fungorum | Betaproteobacteria | Burkholderiales | Burkholderiaceae |
| gi\|WP_158952235.1 | Paraburkholderia acidisoli | Betaproteobacteria | Burkholderiales | Burkholderiaceae |
| gi\|WP_121278480.1 | Trinickia fusca | Betaproteobacteria | Burkholderiales | Burkholderiaceae |
| gi\|WP_006758221.1 | Burkholderia ambifaria | Betaproteobacteria | Burkholderiales | Burkholderiaceae |
| gi\|RKT22386.1 | Paraburkholderia sp. RAU2J | Betaproteobacteria | Burkholderiales | Burkholderiaceae |
| gi\|PLZ02209.1 | Burkholderia sp. WAC0059 | Betaproteobacteria | Burkholderiales | Burkholderiaceae |
| gi\|WP_087633833.1 | Caballeronia telluris | Betaproteobacteria | Burkholderiales | Burkholderiaceae |
| gi\|WP_144138188.1 | Paraburkholderia sp. BCC1884 | Betaproteobacteria | Burkholderiales | Burkholderiaceae |
| gi\|WP_144152001.1 | Paraburkholderia sp. BCC1885 | Betaproteobacteria | Burkholderiales | Burkholderiaceae |
| gi\|WP_121086208.1 | Pararobbsia silviterrae | Betaproteobacteria | Burkholderiales | Burkholderiaceae |
| gi\|WP_027797095.1 | Paraburkholderia acidipaludis | Betaproteobacteria | Burkholderiales | Burkholderiaceae |
| gi\|WP_069343806.1 | Pandoraea sp. ISTKB | Betaproteobacteria | Burkholderiales | Burkholderiaceae |
| gi\|WP_105510390.1 | Paraburkholderia sp. BL21I4N1 | Betaproteobacteria | Burkholderiales | Burkholderiaceae |
| gi\|WP_176120241.1 | Paraburkholderia youngii | Betaproteobacteria | Burkholderiales | Burkholderiaceae |
| gi\|WP_075297669.1 | Burkholderia sp. SRS-W-2-2016 | Betaproteobacteria | Burkholderiales | Burkholderiaceae |
| gi\|WP_074296964.1 | Paraburkholderia phenazinium | Betaproteobacteria | Burkholderiales | Burkholderiaceae |
| gi\|WP_013342783.1 | Burkholderia sp. CCGE1003 | Betaproteobacteria | Burkholderiales | Burkholderiaceae |
| gi\|WP_153099517.1 | Paraburkholderia hayleyella | Betaproteobacteria | Burkholderiales | Burkholderiaceae |

| | | | | |
|---|---|---|---|---|
| gi\|WP_167061470.1 | Burkholderia sp. Ax-1719 | Betaproteobacteria | Burkholderiales | Burkholderiaceae |
| gi\|WP_175240498.1 | Burkholderia cepacia complex | Betaproteobacteria | Burkholderiales | Burkholderiaceae |
| gi\|AMV44945.1 | Paraburkholderia caribensis | Betaproteobacteria | Burkholderiales | Burkholderiaceae |
| gi\|WP_153076893.1 | Paraburkholderia bonniea | Betaproteobacteria | Burkholderiales | Burkholderiaceae |
| gi\|RQS09125.1 | Burkholderia sp. Bp8998 | Betaproteobacteria | Burkholderiales | Burkholderiaceae |
| gi\|WP_084166997.1 | Paraburkholderia caledonica | Betaproteobacteria | Burkholderiales | Burkholderiaceae |
| gi\|WP_124151235.1 | Paraburkholderia dinghuensis | Betaproteobacteria | Burkholderiales | Burkholderiaceae |
| gi\|WP_087725952.1 | Pandoraea sp. PE-S2T-3 | Betaproteobacteria | Burkholderiales | Burkholderiaceae |
| gi\|WP_091011772.1 | Paraburkholderia megapolitana | Betaproteobacteria | Burkholderiales | Burkholderiaceae |
| gi\|WP_038714572.1 | Burkholderia sp. lig30 | Betaproteobacteria | Burkholderiales | Burkholderiaceae |
| gi\|RAS45593.1 | Burkholderia sp. yr520 | Betaproteobacteria | Burkholderiales | Burkholderiaceae |
| gi\|WP_118182675.1 | Paraburkholderia phosphatilytica | Betaproteobacteria | Burkholderiales | Burkholderiaceae |
| gi\|WP_157636172.1 | Burkholderia ubonensis | Betaproteobacteria | Burkholderiales | Burkholderiaceae |
| gi\|TGN96311.1 | Burkholderia sp. USMB20 | Betaproteobacteria | Burkholderiales | Burkholderiaceae |
| gi\|WP_059576339.1 | pseudomallei group | Betaproteobacteria | Burkholderiales | Burkholderiaceae |
| gi\|TAL79672.1 | Burkholderiaceae bacterium | Betaproteobacteria | Burkholderiales | Burkholderiaceae |
| gi\|WP_010807902.1 | Pandoraea | Betaproteobacteria | Burkholderiales | Burkholderiaceae |
| gi\|EDZ97299.1 | Burkholderia sp. H160 | Betaproteobacteria | Burkholderiales | Burkholderiaceae |
| gi\|TXH57438.1 | Burkholderiaceae bacterium | Betaproteobacteria | Burkholderiales | Burkholderiaceae |
| gi\|AFQ49837.1 | Burkholderia cepacia GG4 | Betaproteobacteria | Burkholderiales | Burkholderiaceae |
| gi\|WP_173260582.1 | Paraburkholderia sp. NMBU_R16 | Betaproteobacteria | Burkholderiales | Burkholderiaceae |
| gi\|WP_074985371.1 | Paraburkholderia tropica | Betaproteobacteria | Burkholderiales | Burkholderiaceae |
| gi\|WP_046569423.1 | Paraburkholderia fungorum | Betaproteobacteria | Burkholderiales | Burkholderiaceae |
| gi\|WP_013700192.1 | Burkholderia gladioli | Betaproteobacteria | Burkholderiales | Burkholderiaceae |
| gi\|OUL97655.1 | Paraburkholderia hospita | Betaproteobacteria | Burkholderiales | Burkholderiaceae |
| gi\|WP_081053151.1 | Burkholderia territorii | Betaproteobacteria | Burkholderiales | Burkholderiaceae |
| gi\|TDQ97254.1 | Caballeronia udeis | Betaproteobacteria | Burkholderiales | Burkholderiaceae |
| gi\|WP_081056211.1 | Burkholderia vietnamiensis | Betaproteobacteria | Burkholderiales | Burkholderiaceae |
| gi\|WP_085481221.1 | Paraburkholderia susongensis | Betaproteobacteria | Burkholderiales | Burkholderiaceae |
| gi\|WP_109481299.1 | Paraburkholderia sp. C35 | Betaproteobacteria | Burkholderiales | Burkholderiaceae |
| gi\|WP_081069325.1 | Burkholderia diffusa | Betaproteobacteria | Burkholderiales | Burkholderiaceae |
| gi\|WP_159834046.1 | Burkholderia sp. 8Y | Betaproteobacteria | Burkholderiales | Burkholderiaceae |
| gi\|WP_043365765.1 | Cupriavidus sp. WS | Betaproteobacteria | Burkholderiales | Burkholderiaceae |
| gi\|WP_087632743.1 | Caballeronia telluris | Betaproteobacteria | Burkholderiales | Burkholderiaceae |
| gi\|WP_061170309.1 | Caballeronia hypogeia | Betaproteobacteria | Burkholderiales | Burkholderiaceae |
| gi\|BAN23295.1 | Caballeronia insecticola | Betaproteobacteria | Burkholderiales | Burkholderiaceae |
| gi\|WP_136898790.1 | Trinickia sp. 7GSK02 | Betaproteobacteria | Burkholderiales | Burkholderiaceae |
| gi\|WP_028218675.1 | Paraburkholderia oxyphila | Betaproteobacteria | Burkholderiales | Burkholderiaceae |
| gi\|KXU87194.1 | Caballeronia megalochromosomata | Betaproteobacteria | Burkholderiales | Burkholderiaceae |
| gi\|WP_028225338.1 | Paraburkholderia ferrariae | Betaproteobacteria | Burkholderiales | Burkholderiaceae |
| gi\|WP_132019812.1 | Burkholderia sp. SRS-46 | Betaproteobacteria | Burkholderiales | Burkholderiaceae |
| gi\|WP_082932879.1 | Ralstonia | Betaproteobacteria | Burkholderiales | Burkholderiaceae |
| gi\|WP_061137486.1 | Caballeronia fortuita | Betaproteobacteria | Burkholderiales | Burkholderiaceae |

| | | | | |
|---|---|---|---|---|
| gi\|WP_137331366.1 | Burkholderia sp. DHOD12 | Betaproteobacteria | Burkholderiales | Burkholderiaceae |
| gi\|SAL25091.1 | Caballeronia turbans | Betaproteobacteria | Burkholderiales | Burkholderiaceae |
| gi\|WP_183707366.1 | Paraburkholderia tropica | Betaproteobacteria | Burkholderiales | Burkholderiaceae |
| gi\|SEA17514.1 | Paraburkholderia sartisoli | Betaproteobacteria | Burkholderiales | Burkholderiaceae |
| gi\|WP_040131171.1 | Burkholderia cepacia complex | Betaproteobacteria | Burkholderiales | Burkholderiaceae |
| gi\|WP_112169669.1 | Paraburkholderia unamae | Betaproteobacteria | Burkholderiales | Burkholderiaceae |
| gi\|WP_080413704.1 | Burkholderia ubonensis | Betaproteobacteria | Burkholderiales | Burkholderiaceae |
| gi\|GGC58311.1 | Paraburkholderia caffeinilytica | Betaproteobacteria | Burkholderiales | Burkholderiaceae |
| gi\|CAD6526668.1 | Paraburkholderia metrosideri | Betaproteobacteria | Burkholderiales | Burkholderiaceae |
| gi\|CAB3684654.1 | Paraburkholderia rhynchosiae | Betaproteobacteria | Burkholderiales | Burkholderiaceae |
| gi\|WP_133197370.1 | Candidatus Paraburkholderia sp. 4M-K11 | Betaproteobacteria | Burkholderiales | Burkholderiaceae |
| gi\|WP_075357783.1 | Caballeronia sordidicola | Betaproteobacteria | Burkholderiales | Burkholderiaceae |
| gi\|WP_150678268.1 | Pandoraea pneumonica | Betaproteobacteria | Burkholderiales | Burkholderiaceae |
| gi\|WP_115102707.1 | Paraburkholderia lacunae | Betaproteobacteria | Burkholderiales | Burkholderiaceae |
| gi\|WP_084515337.1 | Burkholderia sp. WSM2230 | Betaproteobacteria | Burkholderiales | Burkholderiaceae |
| gi\|AZQ53769.1 | Burkholderia cenocepacia | Betaproteobacteria | Burkholderiales | Burkholderiaceae |
| gi\|OXI24215.1 | Burkholderia sp. AU15512 | Betaproteobacteria | Burkholderiales | Burkholderiaceae |
| gi\|WP_105777623.1 | Burkholderia multivorans | Betaproteobacteria | Burkholderiales | Burkholderiaceae |
| gi\|WP_176026281.1 | Robbsia andropogonis | Betaproteobacteria | Burkholderiales | Burkholderiaceae |
| gi\|WP_091011642.1 | Paraburkholderia megapolitana | Betaproteobacteria | Burkholderiales | Burkholderiaceae |
| gi\|WP_052241106.1 | Pandoraea fibrosis | Betaproteobacteria | Burkholderiales | Burkholderiaceae |
| gi\|WP_157123166.1 | Pandoraea vervacti | Betaproteobacteria | Burkholderiales | Burkholderiaceae |
| gi\|WP_193100016.1 | Burkholderia sp. Z1 | Betaproteobacteria | Burkholderiales | Burkholderiaceae |
| gi\|WP_008349763.1 | Caballeronia zhejiangensis | Betaproteobacteria | Burkholderiales | Burkholderiaceae |
| gi\|WP_087135007.1 | Caballeronia arationis | Betaproteobacteria | Burkholderiales | Burkholderiaceae |
| gi\|WP_027794619.1 | Paraburkholderia acidipaludis | Betaproteobacteria | Burkholderiales | Burkholderiaceae |
| gi\|WP_087645763.1 | Caballeronia choica | Betaproteobacteria | Burkholderiales | Burkholderiaceae |
| gi\|WP_088812330.1 | Polynucleobacter victoriensis | Betaproteobacteria | Burkholderiales | Burkholderiaceae |
| gi\|WP_034185774.1 | Burkholderia seminalis | Betaproteobacteria | Burkholderiales | Burkholderiaceae |
| gi\|WP_158900344.1 | Burkholderia sp. L27(2015) | Betaproteobacteria | Burkholderiales | Burkholderiaceae |
| gi\|WP_061137550.1 | Caballeronia fortuita | Betaproteobacteria | Burkholderiales | Burkholderiaceae |
| gi\|WP_144159566.1 | Paraburkholderia sp. BCC1885 | Betaproteobacteria | Burkholderiales | Burkholderiaceae |
| gi\|WP_075643313.1 | Caballeronia sordidicola | Betaproteobacteria | Burkholderiales | Burkholderiaceae |
| gi\|CAB3796379.1 | Pararobbsia alpina | Betaproteobacteria | Burkholderiales | Burkholderiaceae |
| gi\|WP_074294579.1 | Paraburkholderia phenazinium | Betaproteobacteria | Burkholderiales | Burkholderiaceae |
| gi\|WP_136898007.1 | Trinickia sp. 7GSK02 | Betaproteobacteria | Burkholderiales | Burkholderiaceae |
| gi\|WP_169497434.1 | Paraburkholderia sp. G-4-1-8 | Betaproteobacteria | Burkholderiales | Burkholderiaceae |
| gi\|WP_105846929.1 | Burkholderia multivorans | Betaproteobacteria | Burkholderiales | Burkholderiaceae |
| gi\|WP_071753092.1 | Burkholderia ubonensis | Betaproteobacteria | Burkholderiales | Burkholderiaceae |
| gi\|WP_042299658.1 | Paraburkholderia kururiensis | Betaproteobacteria | Burkholderiales | Burkholderiaceae |
| gi\|OXL16367.1 | Polynucleobacter cosmopolitanus | Betaproteobacteria | Burkholderiales | Burkholderiaceae |
| gi\|RQZ62523.1 | Burkholderia cepacia | Betaproteobacteria | Burkholderiales | Burkholderiaceae |
| gi\|WP_175948207.1 | Burkholderia pyrrocinia | Betaproteobacteria | Burkholderiales | Burkholderiaceae |

| | | | | |
|---|---|---|---|---|
| gi\|OYY57966.1 | Polynucleobacter sp. 35-46-207 | Betaproteobacteria | Burkholderiales | Burkholderiaceae |
| gi\|WP_179405532.1 | Burkholderia guangdongensis | Betaproteobacteria | Burkholderiales | Burkholderiaceae |
| gi\|ABA52656.1 | Burkholderia pseudomallei 1710b | Betaproteobacteria | Burkholderiales | Burkholderiaceae |
| gi\|PMS15191.1 | Trinickia dabaoshanensis | Betaproteobacteria | Burkholderiales | Burkholderiaceae |
| gi\|WP_062169839.1 | Burkholderia sp. PAMC 26561 | Betaproteobacteria | Burkholderiales | Burkholderiaceae |
| gi\|WP_084162374.1 | Paraburkholderia bannensis | Betaproteobacteria | Burkholderiales | Burkholderiaceae |
| gi\|WP_045450475.1 | Burkholderia sp. RPE67 | Betaproteobacteria | Burkholderiales | Burkholderiaceae |
| gi\|RXV68922.1 | Burkholderia stabilis | Betaproteobacteria | Burkholderiales | Burkholderiaceae |
| gi\|WP_116135228.1 | Trinickia diaoshuihuensis | Betaproteobacteria | Burkholderiales | Burkholderiaceae |
| gi\|WP_150557668.1 | Pandoraea bronchicola | Betaproteobacteria | Burkholderiales | Burkholderiaceae |
| gi\|WP_175800753.1 | Burkholderia anthina | Betaproteobacteria | Burkholderiales | Burkholderiaceae |
| gi\|CAB3801714.1 | Paraburkholderia caffeinitolerans | Betaproteobacteria | Burkholderiales | Burkholderiaceae |
| gi\|WP_061161290.1 | Caballeronia temeraria | Betaproteobacteria | Burkholderiales | Burkholderiaceae |
| gi\|WP_061161229.1 | Caballeronia temeraria | Betaproteobacteria | Burkholderiales | Burkholderiaceae |
| gi\|WP_155627066.1 | Burkholderia diffusa | Betaproteobacteria | Burkholderiales | Burkholderiaceae |
| gi\|WP_115534954.1 | Trinickia dinghuensis | Betaproteobacteria | Burkholderiales | Burkholderiaceae |
| gi\|WP_090692929.1 | Paraburkholderia phenazinium | Betaproteobacteria | Burkholderiales | Burkholderiaceae |
| gi\|WP_028222506.1 | Paraburkholderia oxyphila | Betaproteobacteria | Burkholderiales | Burkholderiaceae |
| gi\|TXI14535.1 | Polynucleobacter sp. | Betaproteobacteria | Burkholderiales | Burkholderiaceae |
| gi\|WP_144110432.1 | Paraburkholderia sp. BCC1886 | Betaproteobacteria | Burkholderiales | Burkholderiaceae |
| gi\|WP_040048523.1 | Caballeronia concitans | Betaproteobacteria | Burkholderiales | Burkholderiaceae |
| gi\|WP_150697426.1 | Pandoraea terrae | Betaproteobacteria | Burkholderiales | Burkholderiaceae |
| gi\|TAM08479.1 | Paraburkholderia sp. | Betaproteobacteria | Burkholderiales | Burkholderiaceae |
| gi\|WP_137958770.1 | Burkholderia sp. 4M9327F10 | Betaproteobacteria | Burkholderiales | Burkholderiaceae |
| gi\|WP_082117889.1 | Pandoraea apista | Betaproteobacteria | Burkholderiales | Burkholderiaceae |
| gi\|WP_175940274.1 | Caballeronia sp. BCC1704 | Betaproteobacteria | Burkholderiales | Burkholderiaceae |
| gi\|WP_174987639.1 | Pandoraea pneumonica | Betaproteobacteria | Burkholderiales | Burkholderiaceae |
| gi\|EAY62524.1 | Burkholderia cenocepacia PC184 | Betaproteobacteria | Burkholderiales | Burkholderiaceae |
| gi\|WP_060601924.1 | Paraburkholderia caribensis | Betaproteobacteria | Burkholderiales | Burkholderiaceae |
| gi\|WP_048246784.1 | Burkholderia cepacia | Betaproteobacteria | Burkholderiales | Burkholderiaceae |
| gi\|WP_183969064.1 | Quisquiliibacterium transsilvanicum | Betaproteobacteria | Burkholderiales | Burkholderiaceae |
| gi\|WP_175113913.1 | Paraburkholderia solisilvae | Betaproteobacteria | Burkholderiales | Burkholderiaceae |
| gi\|WP_078223459.1 | Ralstonia | Betaproteobacteria | Burkholderiales | Burkholderiaceae |
| gi\|KMZ11866.1 | Candidatus Burkholderia humilis | Betaproteobacteria | Burkholderiales | Burkholderiaceae |
| gi\|WP_137334758.1 | Burkholderia sp. DHOD12 | Betaproteobacteria | Burkholderiales | Burkholderiaceae |
| gi\|WP_013434300.1 | Mycetohabitans rhizoxinica | Betaproteobacteria | Burkholderiales | Burkholderiaceae |
| gi\|WP_125095792.1 | Lautropia dentalis | Betaproteobacteria | Burkholderiales | Burkholderiaceae |
| gi\|WP_045594029.1 | Burkholderia multivorans | Betaproteobacteria | Burkholderiales | Burkholderiaceae |
| gi\|WP_059963759.1 | Burkholderia ubonensis | Betaproteobacteria | Burkholderiales | Burkholderiaceae |
| gi\|WP_175112689.1 | Paraburkholderia solisilvae | Betaproteobacteria | Burkholderiales | Burkholderiaceae |
| gi\|WP_134042809.1 | Paraburkholderia caballeronis | Betaproteobacteria | Burkholderiales | Burkholderiaceae |
| gi\|WP_137957350.1 | Burkholderia sp. 4M9327F10 | Betaproteobacteria | Burkholderiales | Burkholderiaceae |
| gi\|WP_089341799.1 | Burkholderia singularis | Betaproteobacteria | Burkholderiales | Burkholderiaceae |

| | | | | |
|---|---|---|---|---|
| gi\|APD12407.1 | Pandoraea sputorum | Betaproteobacteria | Burkholderiales | Burkholderiaceae |
| gi\|WP_128112458.1 | Polynucleobacter necessarius | Betaproteobacteria | Burkholderiales | Burkholderiaceae |
| gi\|WP_102646504.1 | Trinickia dabaoshanensis | Betaproteobacteria | Burkholderiales | Burkholderiaceae |
| gi\|WP_052760167.1 | Burkholderia | Betaproteobacteria | Burkholderiales | Burkholderiaceae |
| gi\|WP_150622392.1 | Pandoraea horticolens | Betaproteobacteria | Burkholderiales | Burkholderiaceae |
| gi\|WP_028227984.1 | Paraburkholderia ferrariae | Betaproteobacteria | Burkholderiales | Burkholderiaceae |
| gi\|WP_124510479.1 | Burkholderia sp. Bp9125 | Betaproteobacteria | Burkholderiales | Burkholderiaceae |
| gi\|WP_150808515.1 | Pandoraea sputorum | Betaproteobacteria | Burkholderiales | Burkholderiaceae |
| gi\|WP_025990335.1 | Burkholderia oklahomensis | Betaproteobacteria | Burkholderiales | Burkholderiaceae |
| gi\|WP_133645185.1 | Paraburkholderia sp. LD6 | Betaproteobacteria | Burkholderiales | Burkholderiaceae |
| gi\|WP_180726525.1 | Paraburkholderia sp. PGU16 | Betaproteobacteria | Burkholderiales | Burkholderiaceae |
| gi\|WP_091905239.1 | Burkholderia sp. JS23 | Betaproteobacteria | Burkholderiales | Burkholderiaceae |
| gi\|WP_035967416.1 | Caballeronia grimmiae | Betaproteobacteria | Burkholderiales | Burkholderiaceae |
| gi\|WP_090683232.1 | Paraburkholderia phenazinium | Betaproteobacteria | Burkholderiales | Burkholderiaceae |
| gi\|WP_028218685.1 | Paraburkholderia oxyphila | Betaproteobacteria | Burkholderiales | Burkholderiaceae |
| gi\|WP_052001354.1 | Burkholderia | Betaproteobacteria | Burkholderiales | Burkholderiaceae |
| gi\|WP_084908212.1 | Paraburkholderia acidophila | Betaproteobacteria | Burkholderiales | Burkholderiaceae |
| gi\|WP_080554728.1 | Burkholderia thailandensis | Betaproteobacteria | Burkholderiales | Burkholderiaceae |
| gi\|WP_027803788.1 | Paraburkholderia dilworthii | Betaproteobacteria | Burkholderiales | Burkholderiaceae |
| gi\|WP_017772942.1 | Paraburkholderia kururiensis | Betaproteobacteria | Burkholderiales | Burkholderiaceae |
| gi\|WP_181969689.1 | Paraburkholderia sp. DHOC27 | Betaproteobacteria | Burkholderiales | Burkholderiaceae |
| gi\|WP_121278021.1 | Trinickia fusca | Betaproteobacteria | Burkholderiales | Burkholderiaceae |
| gi\|WP_014899987.1 | Burkholderia cepacia | Betaproteobacteria | Burkholderiales | Burkholderiaceae |
| gi\|WP_132451256.1 | Paraburkholderia sp. BL8N3 | Betaproteobacteria | Burkholderiales | Burkholderiaceae |
| gi\|SAK77536.1 | Caballeronia hypogeia | Betaproteobacteria | Burkholderiales | Burkholderiaceae |
| gi\|WP_124603295.1 | Burkholderia sp. Bp8963 | Betaproteobacteria | Burkholderiales | Burkholderiaceae |
| gi\|WP_093633305.1 | Paraburkholderia aspalathi | Betaproteobacteria | Burkholderiales | Burkholderiaceae |
| gi\|KND61566.1 | Candidatus Burkholderia verschuerenii | Betaproteobacteria | Burkholderiales | Burkholderiaceae |
| gi\|ABB07714.1 | Burkholderia lata | Betaproteobacteria | Burkholderiales | Burkholderiaceae |
| gi\|WP_183727367.1 | Paraburkholderia | Betaproteobacteria | Burkholderiales | Burkholderiaceae |
| gi\|WP_153076393.1 | Paraburkholderia bonniea | Betaproteobacteria | Burkholderiales | Burkholderiaceae |
| gi\|WP_118182674.1 | Paraburkholderia phosphatilytica | Betaproteobacteria | Burkholderiales | Burkholderiaceae |
| gi\|WP_150583243.1 | Pandoraea communis | Betaproteobacteria | Burkholderiales | Burkholderiaceae |
| gi\|WP_124152555.1 | Paraburkholderia dinghuensis | Betaproteobacteria | Burkholderiales | Burkholderiaceae |
| gi\|WP_150986129.1 | Cupriavidus basilensis | Betaproteobacteria | Burkholderiales | Burkholderiaceae |
| gi\|TDV37340.1 | Paraburkholderia caballeronis | Betaproteobacteria | Burkholderiales | Burkholderiaceae |
| gi\|WP_150789421.1 | Pandoraea iniqua | Betaproteobacteria | Burkholderiales | Burkholderiaceae |
| gi\|WP_107151144.1 | Trinickia symbiotica | Betaproteobacteria | Burkholderiales | Burkholderiaceae |
| gi\|WP_062918288.1 | Paraburkholderia caribensis | Betaproteobacteria | Burkholderiales | Burkholderiaceae |
| gi\|RPA01407.1 | Burkholderia pseudomallei | Betaproteobacteria | Burkholderiales | Burkholderiaceae |
| gi\|WP_051319000.1 | Chitinimonas koreensis | Betaproteobacteria | Burkholderiales | Burkholderiaceae |
| gi\|WP_048812074.1 | Polynucleobacter asymbioticus | Betaproteobacteria | Burkholderiales | Burkholderiaceae |
| gi\|WP_104927585.1 | Pandoraea apista | Betaproteobacteria | Burkholderiales | Burkholderiaceae |

| | | | | |
|---|---|---|---|---|
| gi\|WP_082252657.1 | pseudomallei group | Betaproteobacteria | Burkholderiales | Burkholderiaceae |
| gi\|WP_096672309.1 | Polynucleobacter meluiroseus | Betaproteobacteria | Burkholderiales | Burkholderiaceae |
| gi\|WP_102129888.1 | Burkholderia sp. WAC0059 | Betaproteobacteria | Burkholderiales | Burkholderiaceae |
| gi\|WP_150624988.1 | Pandoraea captiosa | Betaproteobacteria | Burkholderiales | Burkholderiaceae |
| gi\|WP_105777541.1 | Burkholderia multivorans | Betaproteobacteria | Burkholderiales | Burkholderiaceae |
| gi\|WP_087647039.1 | Caballeronia choica | Betaproteobacteria | Burkholderiales | Burkholderiaceae |
| gi\|WP_175698199.1 | Burkholderia ambifaria | Betaproteobacteria | Burkholderiales | Burkholderiaceae |
| gi\|WP_027797436.1 | Paraburkholderia acidipaludis | Betaproteobacteria | Burkholderiales | Burkholderiaceae |
| gi\|WP_051381275.1 | Paraburkholderia mimosarum | Betaproteobacteria | Burkholderiales | Burkholderiaceae |
| gi\|WP_046424722.1 | Burkholderia vietnamiensis | Betaproteobacteria | Burkholderiales | Burkholderiaceae |
| gi\|WP_124599509.1 | Burkholderia sp. Bp8963 | Betaproteobacteria | Burkholderiales | Burkholderiaceae |
| gi\|WP_158952575.1 | Paraburkholderia acidisoli | Betaproteobacteria | Burkholderiales | Burkholderiaceae |
| gi\|WP_147298022.1 | Trinickia dinghuensis | Betaproteobacteria | Burkholderiales | Burkholderiaceae |
| gi\|WP_183708418.1 | Paraburkholderia tropica | Betaproteobacteria | Burkholderiales | Burkholderiaceae |
| gi\|WP_086910126.1 | Paraburkholderia hospita | Betaproteobacteria | Burkholderiales | Burkholderiaceae |
| gi\|WP_124149820.1 | Paraburkholderia dinghuensis | Betaproteobacteria | Burkholderiales | Burkholderiaceae |
| gi\|WP_047214942.1 | Pandoraea thiooxydans | Betaproteobacteria | Burkholderiales | Burkholderiaceae |
| gi\|WP_082088434.1 | Burkholderia sp. USMB20 | Betaproteobacteria | Burkholderiales | Burkholderiaceae |
| gi\|SAL45719.1 | Caballeronia terrestris | Betaproteobacteria | Burkholderiales | Burkholderiaceae |
| gi\|WP_114636624.1 | Polynucleobacter necessarius | Betaproteobacteria | Burkholderiales | Burkholderiaceae |
| gi\|WP_103704311.1 | Paraburkholderia eburnea | Betaproteobacteria | Burkholderiales | Burkholderiaceae |
| gi\|CUV31004.1 | Ralstonia solanacearum | Betaproteobacteria | Burkholderiales | Burkholderiaceae |
| gi\|WP_111488689.1 | Paraburkholderia sp. PDC91 | Betaproteobacteria | Burkholderiales | Burkholderiaceae |
| gi\|WP_107151470.1 | Trinickia symbiotica | Betaproteobacteria | Burkholderiales | Burkholderiaceae |
| gi\|WP_059516642.1 | Burkholderia pseudomultivorans | Betaproteobacteria | Burkholderiales | Burkholderiaceae |
| gi\|WP_028218692.1 | Paraburkholderia oxyphila | Betaproteobacteria | Burkholderiales | Burkholderiaceae |
| gi\|WP_108647762.1 | Polynucleobacter rarus | Betaproteobacteria | Burkholderiales | Burkholderiaceae |
| gi\|CAD18423.1 | Ralstonia solanacearum GMI1000 | Betaproteobacteria | Burkholderiales | Burkholderiaceae |
| gi\|WP_042115943.1 | Pandoraea apista | Betaproteobacteria | Burkholderiales | Burkholderiaceae |
| gi\|WP_006758218.1 | Burkholderia ambifaria | Betaproteobacteria | Burkholderiales | Burkholderiaceae |
| gi\|WP_150553739.1 | Pandoraea nosoerga | Betaproteobacteria | Burkholderiales | Burkholderiaceae |
| gi\|WP_059544769.1 | Burkholderia latens | Betaproteobacteria | Burkholderiales | Burkholderiaceae |
| gi\|WP_115100932.1 | Paraburkholderia lacunae | Betaproteobacteria | Burkholderiales | Burkholderiaceae |
| gi\|WP_124380659.1 | Ralstonia sp. SET104 | Betaproteobacteria | Burkholderiales | Burkholderiaceae |
| gi\|WP_152765406.1 | Paraburkholderia franconis | Betaproteobacteria | Burkholderiales | Burkholderiaceae |
| gi\|WP_088526804.1 | Polynucleobacter aenigmaticus | Betaproteobacteria | Burkholderiales | Burkholderiaceae |
| gi\|KND56431.1 | Candidatus Paraburkholderia kirkii | Betaproteobacteria | Burkholderiales | Burkholderiaceae |
| gi\|WP_133660560.1 | Paraburkholderia sp. BL10I2N1 | Betaproteobacteria | Burkholderiales | Burkholderiaceae |
| gi\|WP_027820607.1 | Paraburkholderia bannensis | Betaproteobacteria | Burkholderiales | Burkholderiaceae |
| gi\|WP_116565114.1 | Paraburkholderia sp. OV555 | Betaproteobacteria | Burkholderiales | Burkholderiaceae |
| gi\|WP_134190258.1 | Paraburkholderia rhizosphaerae | Betaproteobacteria | Burkholderiales | Burkholderiaceae |
| gi\|WP_042337411.1 | Paraburkholderia ferrariae | Betaproteobacteria | Burkholderiales | Burkholderiaceae |
| gi\|WP_105777595.1 | Burkholderia multivorans | Betaproteobacteria | Burkholderiales | Burkholderiaceae |

| | | | | |
|---|---|---|---|---|
| gi\|WP_063750814.1 | Paraburkholderia nodosa | Betaproteobacteria | Burkholderiales | Burkholderiaceae |
| gi\|WP_133665933.1 | Paraburkholderia sp. BL10I2N1 | Betaproteobacteria | Burkholderiales | Burkholderiaceae |
| gi\|WP_137333434.1 | Burkholderia sp. DHOD12 | Betaproteobacteria | Burkholderiales | Burkholderiaceae |
| gi\|WP_111483229.1 | Paraburkholderia sp. PDC91 | Betaproteobacteria | Burkholderiales | Burkholderiaceae |
| gi\|WP_092003017.1 | Paraburkholderia lycopersici | Betaproteobacteria | Burkholderiales | Burkholderiaceae |
| gi\|WP_071021039.1 | Cupriavidus | Betaproteobacteria | Burkholderiales | Burkholderiaceae |
| gi\|WP_159836118.1 | Burkholderia sp. 8Y | Betaproteobacteria | Burkholderiales | Burkholderiaceae |
| gi\|WP_085226387.1 | Trinickia caryophylli | Betaproteobacteria | Burkholderiales | Burkholderiaceae |
| gi\|PRE45526.1 | Burkholderia multivorans | Betaproteobacteria | Burkholderiales | Burkholderiaceae |
| gi\|WP_187292919.1 | Ralstonia solanacearum | Betaproteobacteria | Burkholderiales | Burkholderiaceae |
| gi\|WP_035492196.1 | Paraburkholderia atlantica | Betaproteobacteria | Burkholderiales | Burkholderiaceae |
| gi\|WP_082729664.1 | Burkholderia sp. FL-7-2-10-S1-D7 | Betaproteobacteria | Burkholderiales | Burkholderiaceae |
| gi\|WP_173961232.1 | Polynucleobacter asymbioticus | Betaproteobacteria | Burkholderiales | Burkholderiaceae |
| gi\|WP_100380043.1 | Polynucleobacter sp. UB-Domo-W1 | Betaproteobacteria | Burkholderiales | Burkholderiaceae |
| gi\|WP_175108508.1 | Pararobbsia alpina | Betaproteobacteria | Burkholderiales | Burkholderiaceae |
| gi\|KND58159.1 | Candidatus Paraburkholderia schumannianae | Betaproteobacteria | Burkholderiales | Burkholderiaceae |
| gi\|WP_116610141.1 | Paraburkholderia unamae | Betaproteobacteria | Burkholderiales | Burkholderiaceae |
| gi\|WP_150669354.1 | Pandoraea anhela | Betaproteobacteria | Burkholderiales | Burkholderiaceae |
| gi\|TCT32253.1 | Burkholderia vietnamiensis | Betaproteobacteria | Burkholderiales | Burkholderiaceae |
| gi\|WP_175225184.1 | Paraburkholderia humisilvae | Betaproteobacteria | Burkholderiales | Burkholderiaceae |
| gi\|WP_114809493.1 | Paraburkholderia kururiensis | Betaproteobacteria | Burkholderiales | Burkholderiaceae |
| gi\|WP_059714386.1 | Burkholderia ubonensis | Betaproteobacteria | Burkholderiales | Burkholderiaceae |
| gi\|WP_047909463.1 | Pandoraea faecigallinarum | Betaproteobacteria | Burkholderiales | Burkholderiaceae |
| gi\|WP_182594990.1 | Ralstonia pickettii | Betaproteobacteria | Burkholderiales | Burkholderiaceae |
| gi\|WP_174425429.1 | Cupriavidus basilensis | Betaproteobacteria | Burkholderiales | Burkholderiaceae |
| gi\|KQR87038.1 | Burkholderia sp. Leaf177 | Betaproteobacteria | Burkholderiales | Burkholderiaceae |
| gi\|OZB48178.1 | Polynucleobacter sp. 39-45-136 | Betaproteobacteria | Burkholderiales | Burkholderiaceae |
| gi\|AEG71877.1 | Ralstonia solanacearum Po82 | Betaproteobacteria | Burkholderiales | Burkholderiaceae |
| gi\|WP_084285901.1 | Polynucleobacter sp. VK13 | Betaproteobacteria | Burkholderiales | Burkholderiaceae |
| gi\|VXB92781.1 | Burkholderia sp. 8Y | Betaproteobacteria | Burkholderiales | Burkholderiaceae |
| gi\|APD13497.1 | Pandoraea pulmonicola | Betaproteobacteria | Burkholderiales | Burkholderiaceae |
| gi\|WP_076023594.1 | Polynucleobacter sphagniphilus | Betaproteobacteria | Burkholderiales | Burkholderiaceae |
| gi\|WP_183708412.1 | Paraburkholderia tropica | Betaproteobacteria | Burkholderiales | Burkholderiaceae |
| gi\|WP_102612338.1 | Trinickia soli | Betaproteobacteria | Burkholderiales | Burkholderiaceae |
| gi\|RSD12152.1 | Pandoraea apista | Betaproteobacteria | Burkholderiales | Burkholderiaceae |
| gi\|WP_080562479.1 | Burkholderia gladioli | Betaproteobacteria | Burkholderiales | Burkholderiaceae |
| gi\|WP_070106024.1 | Burkholderia plantarii | Betaproteobacteria | Burkholderiales | Burkholderiaceae |
| gi\|SAL62820.1 | Caballeronia peredens | Betaproteobacteria | Burkholderiales | Burkholderiaceae |
| gi\|WP_176956912.1 | Paraburkholderia caribensis | Betaproteobacteria | Burkholderiales | Burkholderiaceae |
| gi\|WP_072583813.1 | unclassified Polynucleobacter | Betaproteobacteria | Burkholderiales | Burkholderiaceae |
| gi\|WP_089475364.1 | Burkholderia sp. AU6039 | Betaproteobacteria | Burkholderiales | Burkholderiaceae |
| gi\|CCA86638.1 | Ralstonia syzygii R24 | Betaproteobacteria | Burkholderiales | Burkholderiaceae |
| gi\|ODS95011.1 | Lautropia sp. SCN 69-89 | Betaproteobacteria | Burkholderiales | Burkholderiaceae |

| | | | | |
|---|---|---|---|---|
| gi\|WP_084068779.1 | Paraburkholderia heleia | Betaproteobacteria | Burkholderiales | Burkholderiaceae |
| gi\|WP_012358093.1 | Polynucleobacter necessarius | Betaproteobacteria | Burkholderiales | Burkholderiaceae |
| gi\|WP_047839771.1 | Burkholderia gladioli | Betaproteobacteria | Burkholderiales | Burkholderiaceae |
| gi\|KJK24753.1 | Burkholderiaceae bacterium 16 | Betaproteobacteria | Burkholderiales | Burkholderiaceae |
| gi\|WP_085229887.1 | Trinickia caryophylli | Betaproteobacteria | Burkholderiales | Burkholderiaceae |
| gi\|WP_044041301.1 | Caballeronia insecticola | Betaproteobacteria | Burkholderiales | Burkholderiaceae |
| gi\|WP_191629081.1 | Pandoraea terrae | Betaproteobacteria | Burkholderiales | Burkholderiaceae |
| gi\|KXU88878.1 | Caballeronia megalochromosomata | Betaproteobacteria | Burkholderiales | Burkholderiaceae |
| gi\|WP_143107338.1 | Burkholderia pseudomallei | Betaproteobacteria | Burkholderiales | Burkholderiaceae |
| gi\|OYZ36565.1 | Polynucleobacter sp. 16-46-70 | Betaproteobacteria | Burkholderiales | Burkholderiaceae |
| gi\|WP_061124678.1 | Caballeronia catudaia | Betaproteobacteria | Burkholderiales | Burkholderiaceae |
| gi\|WP_038710762.1 | Burkholderia sp. lig30 | Betaproteobacteria | Burkholderiales | Burkholderiaceae |
| gi\|EON14617.1 | Pandoraea sp. SD6-2 | Betaproteobacteria | Burkholderiales | Burkholderiaceae |
| gi\|WP_112313792.1 | Polynucleobacter paneuropaeus | Betaproteobacteria | Burkholderiales | Burkholderiaceae |
| gi\|WP_039598211.1 | Ralstonia sp. A12 | Betaproteobacteria | Burkholderiales | Burkholderiaceae |
| gi\|WP_121278482.1 | Trinickia fusca | Betaproteobacteria | Burkholderiales | Burkholderiaceae |
| gi\|ACC69870.1 | Paraburkholderia phymatum STM815 | Betaproteobacteria | Burkholderiales | Burkholderiaceae |
| gi\|WP_114638870.1 | Polynucleobacter necessarius | Betaproteobacteria | Burkholderiales | Burkholderiaceae |
| gi\|SAK56199.1 | Caballeronia ptereochthonis | Betaproteobacteria | Burkholderiales | Burkholderiaceae |
| gi\|WP_175233010.1 | Paraburkholderia humisilvae | Betaproteobacteria | Burkholderiales | Burkholderiaceae |
| gi\|WP_089340329.1 | Burkholderia singularis | Betaproteobacteria | Burkholderiales | Burkholderiaceae |
| gi\|WP_085230303.1 | Trinickia caryophylli | Betaproteobacteria | Burkholderiales | Burkholderiaceae |
| gi\|WP_173942207.1 | Polynucleobacter sp. LimPoW16 | Betaproteobacteria | Burkholderiales | Burkholderiaceae |
| gi\|WP_124843144.1 | Burkholderia cepacia | Betaproteobacteria | Burkholderiales | Burkholderiaceae |
| gi\|WP_150670722.1 | Pandoraea anhela | Betaproteobacteria | Burkholderiales | Burkholderiaceae |
| gi\|WP_028218695.1 | Paraburkholderia oxyphila | Betaproteobacteria | Burkholderiales | Burkholderiaceae |
| gi\|WP_047906934.1 | Pandoraea faecigallinarum | Betaproteobacteria | Burkholderiales | Burkholderiaceae |
| gi\|TXH64264.1 | Burkholderiaceae bacterium | Betaproteobacteria | Burkholderiales | Burkholderiaceae |
| gi\|WP_116136662.1 | Trinickia diaoshuihuensis | Betaproteobacteria | Burkholderiales | Burkholderiaceae |
| gi\|WP_011493227.1 | Paraburkholderia xenovorans | Betaproteobacteria | Burkholderiales | Burkholderiaceae |
| gi\|WP_175197665.1 | Paraburkholderia caffeinitolerans | Betaproteobacteria | Burkholderiales | Burkholderiaceae |
| gi\|WP_034473600.1 | Caballeronia zhejiangensis | Betaproteobacteria | Burkholderiales | Burkholderiaceae |
| gi\|WP_167336081.1 | Paraburkholderia bannensis | Betaproteobacteria | Burkholderiales | Burkholderiaceae |
| gi\|WP_124383137.1 | Ralstonia sp. SET104 | Betaproteobacteria | Burkholderiales | Burkholderiaceae |
| gi\|WP_125347395.1 | Pandoraea apista | Betaproteobacteria | Burkholderiales | Burkholderiaceae |
| gi\|WP_183727359.1 | Paraburkholderia | Betaproteobacteria | Burkholderiales | Burkholderiaceae |
| gi\|KMY85720.1 | Candidatus Paraburkholderia calva | Betaproteobacteria | Burkholderiales | Burkholderiaceae |
| gi\|WP_052420836.1 | Paraburkholderia ferrariae | Betaproteobacteria | Burkholderiales | Burkholderiaceae |
| gi\|WP_162065514.1 | Burkholderia sp. THE68 | Betaproteobacteria | Burkholderiales | Burkholderiaceae |
| gi\|TAM51497.1 | Paraburkholderia sp. | Betaproteobacteria | Burkholderiales | Burkholderiaceae |
| gi\|ODS96386.1 | Lautropia sp. SCN 69-89 | Betaproteobacteria | Burkholderiales | Burkholderiaceae |
| gi\|WP_186023450.1 | Burkholderia gladioli | Betaproteobacteria | Burkholderiales | Burkholderiaceae |
| gi\|WP_084068777.1 | Paraburkholderia heleia | Betaproteobacteria | Burkholderiales | Burkholderiaceae |

| | | | | |
|---|---|---|---|---|
| gi\|WP_075583209.1 | Caballeronia grimmiae | Betaproteobacteria | Burkholderiales | Burkholderiaceae |
| gi\|WP_105777620.1 | Burkholderia multivorans | Betaproteobacteria | Burkholderiales | Burkholderiaceae |
| gi\|WP_061168608.1 | Caballeronia hypogeia | Betaproteobacteria | Burkholderiales | Burkholderiaceae |
| gi\|GAQ28711.1 | Ralstonia sp. NT80 | Betaproteobacteria | Burkholderiales | Burkholderiaceae |
| gi\|WP_047908000.1 | Pandoraea faecigallinarum | Betaproteobacteria | Burkholderiales | Burkholderiaceae |
| gi\|SAL26363.1 | Caballeronia turbans | Betaproteobacteria | Burkholderiales | Burkholderiaceae |
| gi\|WP_086971214.1 | Caballeronia glebae | Betaproteobacteria | Burkholderiales | Burkholderiaceae |
| gi\|WP_062604419.1 | Caballeronia calidae | Betaproteobacteria | Burkholderiales | Burkholderiaceae |
| gi\|WP_107154243.1 | Trinickia symbiotica | Betaproteobacteria | Burkholderiales | Burkholderiaceae |
| gi\|KIG10476.1 | Burkholderia sp. MR1 | Betaproteobacteria | Burkholderiales | Burkholderiaceae |
| gi\|WP_088524746.1 | Polynucleobacter campilacus | Betaproteobacteria | Burkholderiales | Burkholderiaceae |
| gi\|KAK49008.1 | Caballeronia jiangsuensis | Betaproteobacteria | Burkholderiales | Burkholderiaceae |
| gi\|WP_046329799.1 | Polynucleobacter duraquae | Betaproteobacteria | Burkholderiales | Burkholderiaceae |
| gi\|TAL64197.1 | Burkholderiaceae bacterium | Betaproteobacteria | Burkholderiales | Burkholderiaceae |
| gi\|ARM00259.1 | Burkholderia pseudomallei | Betaproteobacteria | Burkholderiales | Burkholderiaceae |
| gi\|KDR24799.1 | Caballeronia grimmiae | Betaproteobacteria | Burkholderiales | Burkholderiaceae |
| gi\|WP_116610138.1 | Paraburkholderia unamae | Betaproteobacteria | Burkholderiales | Burkholderiaceae |
| gi\|WP_074168874.1 | Caballeronia fortuita | Betaproteobacteria | Burkholderiales | Burkholderiaceae |
| gi\|WP_150777794.1 | Pandoraea sputorum | Betaproteobacteria | Burkholderiales | Burkholderiaceae |
| gi\|WP_010112857.1 | Burkholderia oklahomensis | Betaproteobacteria | Burkholderiales | Burkholderiaceae |
| gi\|WP_058371704.1 | Pandoraea pnomenusa | Betaproteobacteria | Burkholderiales | Burkholderiaceae |
| gi\|ODS99416.1 | Lautropia sp. SCN 69-89 | Betaproteobacteria | Burkholderiales | Burkholderiaceae |
| gi\|WP_086914794.1 | Paraburkholderia hospita | Betaproteobacteria | Burkholderiales | Burkholderiaceae |
| gi\|WP_175940193.1 | Caballeronia sp. M1242 | Betaproteobacteria | Burkholderiales | Burkholderiaceae |
| gi\|WP_176070248.1 | Schlegelella koreensis | Betaproteobacteria | Burkholderiales | Comamonadaceae |
| gi\|WP_066255852.1 | Hydrogenophaga flava | Betaproteobacteria | Burkholderiales | Comamonadaceae |
| gi\|WP_133603824.1 | Kinneretia asaccharophila | Betaproteobacteria | Burkholderiales | Comamonadaceae |
| gi\|ACS18703.1 | Variovorax paradoxus S110 | Betaproteobacteria | Burkholderiales | Comamonadaceae |
| gi\|OSZ73282.1 | Hydrogenophaga sp. IBVHS1 | Betaproteobacteria | Burkholderiales | Comamonadaceae |
| gi\|WP_105729888.1 | Malikia spinosa | Betaproteobacteria | Burkholderiales | Comamonadaceae |
| gi\|WP_077334200.1 | Hydrogenophaga sp. A37 | Betaproteobacteria | Burkholderiales | Comamonadaceae |
| gi\|WP_114483761.1 | Extensimonas vulgaris | Betaproteobacteria | Burkholderiales | Comamonadaceae |
| gi\|WP_051391950.1 | Rhodoferax saidenbachensis | Betaproteobacteria | Burkholderiales | Comamonadaceae |
| gi\|WP_137919170.1 | Hydrogenophaga sp. 2FB | Betaproteobacteria | Burkholderiales | Comamonadaceae |
| gi\|WP_157077060.1 | Curvibacter delicatus | Betaproteobacteria | Burkholderiales | Comamonadaceae |
| gi\|WP_183302386.1 | Comamonas terrigena | Betaproteobacteria | Burkholderiales | Comamonadaceae |
| gi\|WP_105747873.1 | Malikia granosa | Betaproteobacteria | Burkholderiales | Comamonadaceae |
| gi\|WP_180125146.1 | Rhodoferax sp. BLA1 | Betaproteobacteria | Burkholderiales | Comamonadaceae |
| gi\|WP_066273591.1 | Hydrogenophaga palleronii | Betaproteobacteria | Burkholderiales | Comamonadaceae |
| gi\|WP_105747877.1 | Malikia granosa | Betaproteobacteria | Burkholderiales | Comamonadaceae |
| gi\|WP_187306013.1 | Diaphorobacter polyhydroxybutyrativorans | Betaproteobacteria | Burkholderiales | Comamonadaceae |
| gi\|QHE75274.1 | Hydrogenophaga sp. PBL-H3 | Betaproteobacteria | Burkholderiales | Comamonadaceae |
| gi\|RJP69674.1 | Comamonadaceae bacterium | Betaproteobacteria | Burkholderiales | Comamonadaceae |

| | | | | |
|---|---|---|---|---|
| gi\|WP_177224993.1 | Variovorax sp. 770b2 | Betaproteobacteria | Burkholderiales | Comamonadaceae |
| gi\|PRD64688.1 | Malikia granosa | Betaproteobacteria | Burkholderiales | Comamonadaceae |
| gi\|RZL53925.1 | Variovorax sp. | Betaproteobacteria | Burkholderiales | Comamonadaceae |
| gi\|WP_042425391.1 | Comamonas granuli | Betaproteobacteria | Burkholderiales | Comamonadaceae |
| gi\|WP_114968125.1 | Rhodoferax sp. OTU1 | Betaproteobacteria | Burkholderiales | Comamonadaceae |
| gi\|WP_111881685.1 | Acidovorax anthurii | Betaproteobacteria | Burkholderiales | Comamonadaceae |
| gi\|WP_124222962.1 | Tibeticola sediminis | Betaproteobacteria | Burkholderiales | Comamonadaceae |
| gi\|WP_054257520.1 | Acidovorax caeni | Betaproteobacteria | Burkholderiales | Comamonadaceae |
| gi\|RYF36667.1 | Comamonadaceae bacterium | Betaproteobacteria | Burkholderiales | Comamonadaceae |
| gi\|WP_162017393.1 | Acidovorax sp. 210-6 | Betaproteobacteria | Burkholderiales | Comamonadaceae |
| gi\|WP_082876771.1 | Hydrogenophaga crassostreae | Betaproteobacteria | Burkholderiales | Comamonadaceae |
| gi\|WP_136180317.1 | Hydrogenophaga sp. PAMC20947 | Betaproteobacteria | Burkholderiales | Comamonadaceae |
| gi\|WP_176070253.1 | Schlegelella koreensis | Betaproteobacteria | Burkholderiales | Comamonadaceae |
| gi\|WP_158241610.1 | Ottowia sp. Marseille-P4747 | Betaproteobacteria | Burkholderiales | Comamonadaceae |
| gi\|WP_078364545.1 | Rhodoferax fermentans | Betaproteobacteria | Burkholderiales | Comamonadaceae |
| gi\|TXH91378.1 | Rhodoferax sp. | Betaproteobacteria | Burkholderiales | Comamonadaceae |
| gi\|WP_108312095.1 | Limnohabitans parvus | Betaproteobacteria | Burkholderiales | Comamonadaceae |
| gi\|WP_101047371.1 | Macromonas sp. BK-30 | Betaproteobacteria | Burkholderiales | Comamonadaceae |
| gi\|WP_066086811.1 | Hydrogenophaga crassostreae | Betaproteobacteria | Burkholderiales | Comamonadaceae |
| gi\|WP_108326872.1 | unclassified Limnohabitans | Betaproteobacteria | Burkholderiales | Comamonadaceae |
| gi\|ODU56869.1 | Comamonadaceae bacterium SCN 68-20 | Betaproteobacteria | Burkholderiales | Comamonadaceae |
| gi\|PUE43055.1 | Limnohabitans sp. Bal53 | Betaproteobacteria | Burkholderiales | Comamonadaceae |
| gi\|WP_105261610.1 | Rhodoferax sp. TS-BS-61-7 | Betaproteobacteria | Burkholderiales | Comamonadaceae |
| gi\|RYZ11898.1 | Comamonadaceae bacterium | Betaproteobacteria | Burkholderiales | Comamonadaceae |
| gi\|CBA32330.1 | Curvibacter putative symbiont of Hydra magnipapillata | Betaproteobacteria | Burkholderiales | Comamonadaceae |
| gi\|WP_119107948.1 | Simplicispira hankyongi | Betaproteobacteria | Burkholderiales | Comamonadaceae |
| gi\|WP_166227452.1 | Hydrogenophaga sp. BA0156 | Betaproteobacteria | Burkholderiales | Comamonadaceae |
| gi\|WP_119966938.1 | Simplicispira lacusdiani | Betaproteobacteria | Burkholderiales | Comamonadaceae |
| gi\|WP_077563634.1 | Polaromonas sp. C04 | Betaproteobacteria | Burkholderiales | Comamonadaceae |
| gi\|QCB48271.1 | Hydrogenophaga sp. PAMC20947 | Betaproteobacteria | Burkholderiales | Comamonadaceae |
| gi\|WP_158724846.1 | Xenophilus sp. L33 | Betaproteobacteria | Burkholderiales | Comamonadaceae |
| gi\|WP_019572434.1 | Curvibacter lanceolatus | Betaproteobacteria | Burkholderiales | Comamonadaceae |
| gi\|WP_127803494.1 | Hydrogenophaga sp. NH-16 | Betaproteobacteria | Burkholderiales | Comamonadaceae |
| gi\|RZL60800.1 | Variovorax sp. | Betaproteobacteria | Burkholderiales | Comamonadaceae |
| gi\|TXI59901.1 | Limnohabitans sp. | Betaproteobacteria | Burkholderiales | Comamonadaceae |
| gi\|OOG80171.1 | Hydrogenophaga sp. A37 | Betaproteobacteria | Burkholderiales | Comamonadaceae |
| gi\|WP_070397345.1 | unclassified Hydrogenophaga | Betaproteobacteria | Burkholderiales | Comamonadaceae |
| gi\|WP_101048871.1 | Macromonas sp. BK-30 | Betaproteobacteria | Burkholderiales | Comamonadaceae |
| gi\|WP_084382982.1 | Curvibacter delicatus | Betaproteobacteria | Burkholderiales | Comamonadaceae |
| gi\|WP_159595786.1 | Hydrogenophaga sp. BPS33 | Betaproteobacteria | Burkholderiales | Comamonadaceae |
| gi\|WP_177172937.1 | Giesbergeria anulus | Betaproteobacteria | Burkholderiales | Comamonadaceae |
| gi\|WP_094476415.1 | Rhodoferax sp. TH121 | Betaproteobacteria | Burkholderiales | Comamonadaceae |
| gi\|WP_142820722.1 | Rhodoferax sediminis | Betaproteobacteria | Burkholderiales | Comamonadaceae |

| | | | | |
|---|---|---|---|---|
| gi\|RYF83076.1 | Comamonadaceae bacterium | Betaproteobacteria | Burkholderiales | Comamonadaceae |
| gi\|TXT37785.1 | Comamonadaceae bacterium | Betaproteobacteria | Burkholderiales | Comamonadaceae |
| gi\|WP_068167253.1 | Hydrogenophaga taeniospiralis | Betaproteobacteria | Burkholderiales | Comamonadaceae |
| gi\|TAF80276.1 | Curvibacter sp. | Betaproteobacteria | Burkholderiales | Comamonadaceae |
| gi\|WP_005798477.1 | Acidovorax delafieldii | Betaproteobacteria | Burkholderiales | Comamonadaceae |
| gi\|TXH87375.1 | Rhodoferax sp. | Betaproteobacteria | Burkholderiales | Comamonadaceae |
| gi\|WP_195795865.1 | Kinneretia sp. DAIF2 | Betaproteobacteria | Burkholderiales | Comamonadaceae |
| gi\|WP_183302384.1 | Comamonas terrigena | Betaproteobacteria | Burkholderiales | Comamonadaceae |
| gi\|TXI23653.1 | Pelomonas sp. | Betaproteobacteria | Burkholderiales | Comamonadaceae |
| gi\|WP_086923622.1 | Variovorax sp. JS1663 | Betaproteobacteria | Burkholderiales | Comamonadaceae |
| gi\|OHC71850.1 | Rhodoferax sp. RIFCSPLOWO2_12_FULL_60_11 | Betaproteobacteria | Burkholderiales | Comamonadaceae |
| gi\|WP_077032847.1 | Pelomonas sp. KK5 | Betaproteobacteria | Burkholderiales | Comamonadaceae |
| gi\|WP_144729831.1 | Extensimonas perlucida | Betaproteobacteria | Burkholderiales | Comamonadaceae |
| gi\|TMU78024.1 | Hydrogenophaga intermedia | Betaproteobacteria | Burkholderiales | Comamonadaceae |
| gi\|WP_169928998.1 | Macromonas bipunctata | Betaproteobacteria | Burkholderiales | Comamonadaceae |
| gi\|WP_057297262.1 | Pelomonas sp. Root1217 | Betaproteobacteria | Burkholderiales | Comamonadaceae |
| gi\|PTT81865.1 | Pelomonas sp. HMWF004 | Betaproteobacteria | Burkholderiales | Comamonadaceae |
| gi\|WP_122226222.1 | Corticibacter populi | Betaproteobacteria | Burkholderiales | Comamonadaceae |
| gi\|WP_117179915.1 | Rhodoferax sp. IMCC26218 | Betaproteobacteria | Burkholderiales | Comamonadaceae |
| gi\|ODS71389.1 | Acidovorax sp. SCN 68-22 | Betaproteobacteria | Burkholderiales | Comamonadaceae |
| gi\|WP_108136162.1 | unclassified Variovorax | Betaproteobacteria | Burkholderiales | Comamonadaceae |
| gi\|AVO43275.1 | Simplicispira suum | Betaproteobacteria | Burkholderiales | Comamonadaceae |
| gi\|WP_140401010.1 | Comamonas testosteroni | Betaproteobacteria | Burkholderiales | Comamonadaceae |
| gi\|WP_056271985.1 | unclassified Pelomonas | Betaproteobacteria | Burkholderiales | Comamonadaceae |
| gi\|WP_162254490.1 | Pelomonas sp. Root1444 | Betaproteobacteria | Burkholderiales | Comamonadaceae |
| gi\|WP_057201769.1 | Acidovorax sp. Root217 | Betaproteobacteria | Burkholderiales | Comamonadaceae |
| gi\|QDL53006.1 | Rhodoferax sediminis | Betaproteobacteria | Burkholderiales | Comamonadaceae |
| gi\|OYY35267.1 | Polaromonas sp. 35-63-35 | Betaproteobacteria | Burkholderiales | Comamonadaceae |
| gi\|WP_056192754.1 | Pelomonas sp. Root1237 | Betaproteobacteria | Burkholderiales | Comamonadaceae |
| gi\|GGA87294.1 | Polaromonas eurypsychrophila | Betaproteobacteria | Burkholderiales | Comamonadaceae |
| gi\|WP_119047065.1 | Pelomonas sp. BT06 | Betaproteobacteria | Burkholderiales | Comamonadaceae |
| gi\|WP_101102170.1 | Macromonas bipunctata | Betaproteobacteria | Burkholderiales | Comamonadaceae |
| gi\|RZL89421.1 | Variovorax sp. | Betaproteobacteria | Burkholderiales | Comamonadaceae |
| gi\|WP_168173516.1 | Polaromonas sp. A23 | Betaproteobacteria | Burkholderiales | Comamonadaceae |
| gi\|WP_073356089.1 | Lampropedia hyalina | Betaproteobacteria | Burkholderiales | Comamonadaceae |
| gi\|WP_183023245.1 | Variovorax sp. UMC13 | Betaproteobacteria | Burkholderiales | Comamonadaceae |
| gi\|WP_126837546.1 | Variovorax sp. MHTC-1 | Betaproteobacteria | Burkholderiales | Comamonadaceae |
| gi\|WP_093130436.1 | Variovorax sp. OK605 | Betaproteobacteria | Burkholderiales | Comamonadaceae |
| gi\|WP_026433462.1 | Acidovorax oryzae | Betaproteobacteria | Burkholderiales | Comamonadaceae |
| gi\|WP_108285837.1 | Limnohabitans sp. T6-20 | Betaproteobacteria | Burkholderiales | Comamonadaceae |
| gi\|WP_169804384.1 | Polaromonas jejuensis | Betaproteobacteria | Burkholderiales | Comamonadaceae |
| gi\|RYY51961.1 | Comamonadaceae bacterium | Betaproteobacteria | Burkholderiales | Comamonadaceae |
| gi\|WP_092836562.1 | Acidovorax cattleyae | Betaproteobacteria | Burkholderiales | Comamonadaceae |

| | | | | |
|---|---|---|---|---|
| gi\|WP_073353348.1 | Lampropedia hyalina | Betaproteobacteria | Burkholderiales | Comamonadaceae |
| gi\|WP_157045761.1 | Polaromonas sp. JS666 | Betaproteobacteria | Burkholderiales | Comamonadaceae |
| gi\|WP_158219728.1 | Ideonella sp. A 288 | Betaproteobacteria | Burkholderiales | Ideonella |
| gi\|WP_040501240.1 | Ideonella sp. B508-1 | Betaproteobacteria | Burkholderiales | Ideonella |
| gi\|WP_054022556.1 | Ideonella sakaiensis | Betaproteobacteria | Burkholderiales | Ideonella |
| gi\|WP_151123765.1 | Ideonella dechloratans | Betaproteobacteria | Burkholderiales | Ideonella |
| gi\|WP_022980645.1 | Ideonella sp. B508-1 | Betaproteobacteria | Burkholderiales | Ideonella |
| gi\|WP_163457782.1 | Ideonella sp. TBM-1 | Betaproteobacteria | Burkholderiales | Ideonella |
| gi\|WP_088280507.1 | Ideonella sp. A 288 | Betaproteobacteria | Burkholderiales | Ideonella |
| gi\|WP_158219730.1 | Ideonella sp. A 288 | Betaproteobacteria | Burkholderiales | Ideonella |
| gi\|TDM09780.1 | Ideonella sp. MAG2 | Betaproteobacteria | Burkholderiales | Ideonella |
| gi\|WP_083876565.1 | Ideonella sp. B508-1 | Betaproteobacteria | Burkholderiales | Ideonella |
| gi\|WP_088278580.1 | Ideonella sp. A 288 | Betaproteobacteria | Burkholderiales | Ideonella |
| gi\|WP_022980703.1 | Ideonella sp. B508-1 | Betaproteobacteria | Burkholderiales | Ideonella |
| gi\|TDM08334.1 | Ideonella sp. MAG2 | Betaproteobacteria | Burkholderiales | Ideonella |
| gi\|WP_127683872.1 | Inhella crocodyli | Betaproteobacteria | Burkholderiales | Inhella |
| gi\|WP_012347025.1 | Leptothrix cholodnii | Betaproteobacteria | Burkholderiales | Leptothrix |
| gi\|WP_165396725.1 | Leptothrix mobilis | Betaproteobacteria | Burkholderiales | Leptothrix |
| gi\|WP_130481204.1 | Leptothrix mobilis | Betaproteobacteria | Burkholderiales | Leptothrix |
| gi\|WP_083772707.1 | Leptothrix cholodnii | Betaproteobacteria | Burkholderiales | Leptothrix |
| gi\|ABM93648.1 | Methylibium petroleiphilum PM1 | Betaproteobacteria | Burkholderiales | Methylibium |
| gi\|WP_082010865.1 | Methylibium sp. YR605 | Betaproteobacteria | Burkholderiales | Methylibium |
| gi\|WP_156155085.1 | Methylibium sp. CF468 | Betaproteobacteria | Burkholderiales | Methylibium |
| gi\|WP_082004412.1 | Methylibium sp. CF468 | Betaproteobacteria | Burkholderiales | Methylibium |
| gi\|WP_067272139.1 | Mitsuaria sp. 7 | Betaproteobacteria | Burkholderiales | Mitsuaria |
| gi\|WP_082938549.1 | Mitsuaria sp. 7 | Betaproteobacteria | Burkholderiales | Mitsuaria |
| gi\|WP_175538174.1 | Mitsuaria sp. PDC51 | Betaproteobacteria | Burkholderiales | Mitsuaria |
| gi\|WP_067276529.1 | Mitsuaria sp. 7 | Betaproteobacteria | Burkholderiales | Mitsuaria |
| gi\|WP_067061800.1 | Mitsuaria chitosanitabida | Betaproteobacteria | Burkholderiales | Mitsuaria |
| gi\|WP_094299085.1 | Noviherbaspirillum autotrophicum | Betaproteobacteria | Burkholderiales | Oxalobacteraceae |
| gi\|WP_119742249.1 | Herbaspirillum sp. K2R10-39 | Betaproteobacteria | Burkholderiales | Oxalobacteraceae |
| gi\|RBA24508.1 | Herminiimonas fonticola | Betaproteobacteria | Burkholderiales | Oxalobacteraceae |
| gi\|PIG27161.1 | Janthinobacterium sp. 35 | Betaproteobacteria | Burkholderiales | Oxalobacteraceae |
| gi\|CDG83147.1 | Janthinobacterium agaricidamnosum NBRC 102515 = DSM 9628 | Betaproteobacteria | Burkholderiales | Oxalobacteraceae |
| gi\|WP_094299050.1 | Noviherbaspirillum autotrophicum | Betaproteobacteria | Burkholderiales | Oxalobacteraceae |
| gi\|WP_094443920.1 | Janthinobacterium sp. PC23-8 | Betaproteobacteria | Burkholderiales | Oxalobacteraceae |
| gi\|RYE91279.1 | Oxalobacteraceae bacterium | Betaproteobacteria | Burkholderiales | Oxalobacteraceae |
| gi\|WP_152248211.1 | Janthinobacterium sp. FT58W | Betaproteobacteria | Burkholderiales | Oxalobacteraceae |
| gi\|GGB99365.1 | Oxalicibacterium flavum | Betaproteobacteria | Burkholderiales | Oxalobacteraceae |
| gi\|WP_176348640.1 | Massilia sp. BJB1822 | Betaproteobacteria | Burkholderiales | Oxalobacteraceae |
| gi\|WP_182158906.1 | Duganella sp. LX20W | Betaproteobacteria | Burkholderiales | Oxalobacteraceae |
| gi\|WP_136419132.1 | Herbaspirillum sp. ST 5-3 | Betaproteobacteria | Burkholderiales | Oxalobacteraceae |
| gi\|WP_182213588.1 | Duganella sp. FT3S | Betaproteobacteria | Burkholderiales | Oxalobacteraceae |

| | | | | |
|---|---|---|---|---|
| gi\|WP_052234009.1 | Massilia sp. WG5 | Betaproteobacteria | Burkholderiales | Oxalobacteraceae |
| gi\|WP_090437159.1 | Duganella sp. CF458 | Betaproteobacteria | Burkholderiales | Oxalobacteraceae |
| gi\|WP_188422427.1 | Oxalicibacterium solurbis | Betaproteobacteria | Burkholderiales | Oxalobacteraceae |
| gi\|WP_099789097.1 | Massilia eurypsychrophila | Betaproteobacteria | Burkholderiales | Oxalobacteraceae |
| gi\|WP_099792373.1 | Massilia eurypsychrophila | Betaproteobacteria | Burkholderiales | Oxalobacteraceae |
| gi\|WP_156116600.1 | Massilia sp. 9096 | Betaproteobacteria | Burkholderiales | Oxalobacteraceae |
| gi\|WP_161025138.1 | Massilia guangdongensis | Betaproteobacteria | Burkholderiales | Oxalobacteraceae |
| gi\|EGF32907.1 | Oxalobacteraceae bacterium IMCC9480 | Betaproteobacteria | Burkholderiales | Oxalobacteraceae |
| gi\|WP_076592697.1 | Herminiimonas arsenitoxidans | Betaproteobacteria | Burkholderiales | Oxalobacteraceae |
| gi\|WP_154383722.1 | Duganella sp. FT80W | Betaproteobacteria | Burkholderiales | Oxalobacteraceae |
| gi\|WP_170976919.1 | Massilia sp. HP4 | Betaproteobacteria | Burkholderiales | Oxalobacteraceae |
| gi\|WP_025916890.1 | Herminiimonas sp. CN | Betaproteobacteria | Burkholderiales | Oxalobacteraceae |
| gi\|WP_093386890.1 | Rugamonas rubra | Betaproteobacteria | Burkholderiales | Oxalobacteraceae |
| gi\|WP_106757806.1 | Massilia glaciei | Betaproteobacteria | Burkholderiales | Oxalobacteraceae |
| gi\|WP_040042882.1 | Noviherbaspirillum autotrophicum | Betaproteobacteria | Burkholderiales | Oxalobacteraceae |
| gi\|WP_152248247.1 | Janthinobacterium sp. FT58W | Betaproteobacteria | Burkholderiales | Oxalobacteraceae |
| gi\|WP_183681819.1 | unclassified Janthinobacterium | Betaproteobacteria | Burkholderiales | Oxalobacteraceae |
| gi\|WP_175344924.1 | Herbaspirillum sp. C9C3 | Betaproteobacteria | Burkholderiales | Oxalobacteraceae |
| gi\|TQK10285.1 | Herbaspirillum sp. SJZ107 | Betaproteobacteria | Burkholderiales | Oxalobacteraceae |
| gi\|WP_165930450.1 | unclassified Massilia | Betaproteobacteria | Burkholderiales | Oxalobacteraceae |
| gi\|WP_161021394.1 | Duganella sp. FT50W | Betaproteobacteria | Burkholderiales | Oxalobacteraceae |
| gi\|WP_099763228.1 | unclassified Janthinobacterium | Betaproteobacteria | Burkholderiales | Oxalobacteraceae |
| gi\|WP_188564707.1 | Undibacterium terreum | Betaproteobacteria | Burkholderiales | Oxalobacteraceae |
| gi\|WP_146333511.1 | Noviherbaspirillum sp. UKPF54 | Betaproteobacteria | Burkholderiales | Oxalobacteraceae |
| gi\|WP_093388588.1 | Rugamonas rubra | Betaproteobacteria | Burkholderiales | Oxalobacteraceae |
| gi\|WP_133324351.1 | Sapientia aquatica | Betaproteobacteria | Burkholderiales | Oxalobacteraceae |
| gi\|WP_119783777.1 | Noviherbaspirillum sp. K1S02-23 | Betaproteobacteria | Burkholderiales | Oxalobacteraceae |
| gi\|WP_126769918.1 | Undibacterium piscinae | Betaproteobacteria | Burkholderiales | Oxalobacteraceae |
| gi\|WP_056404995.1 | Massilia sp. Root418 | Betaproteobacteria | Burkholderiales | Oxalobacteraceae |
| gi\|WP_128902420.1 | Janthinobacterium sp. 17J80-10 | Betaproteobacteria | Burkholderiales | Oxalobacteraceae |
| gi\|RYE73676.1 | Oxalobacteraceae bacterium | Betaproteobacteria | Burkholderiales | Oxalobacteraceae |
| gi\|TFW10022.1 | Oxalobacteraceae bacterium OM1 | Betaproteobacteria | Burkholderiales | Oxalobacteraceae |
| gi\|PQO97319.1 | Massilia phosphatilytica | Betaproteobacteria | Burkholderiales | Oxalobacteraceae |
| gi\|WP_099759818.1 | unclassified Janthinobacterium | Betaproteobacteria | Burkholderiales | Oxalobacteraceae |
| gi\|WP_186913722.1 | Undibacterium jejuense | Betaproteobacteria | Burkholderiales | Oxalobacteraceae |
| gi\|WP_183681030.1 | unclassified Janthinobacterium | Betaproteobacteria | Burkholderiales | Oxalobacteraceae |
| gi\|WP_192805253.1 | Noviherbaspirillum | Betaproteobacteria | Burkholderiales | Oxalobacteraceae |
| gi\|WP_195876406.1 | Herminiimonas contaminans | Betaproteobacteria | Burkholderiales | Oxalobacteraceae |
| gi\|WP_081466572.1 | Collimonas fungivorans | Betaproteobacteria | Burkholderiales | Oxalobacteraceae |
| gi\|WP_119813095.1 | Massilia sp. K1S02-61 | Betaproteobacteria | Burkholderiales | Oxalobacteraceae |
| gi\|RJF91995.1 | Herbaspirillum sp. K1R23-30 | Betaproteobacteria | Burkholderiales | Oxalobacteraceae |
| gi\|WP_161088588.1 | Rugamonas sp. FT107W | Betaproteobacteria | Burkholderiales | Oxalobacteraceae |
| gi\|WP_152879086.1 | Duganella sp. FT27W | Betaproteobacteria | Burkholderiales | Oxalobacteraceae |

| | | | | |
|---|---|---|---|---|
| gi\|WP_070249633.1 | Duganella phyllosphaerae | Betaproteobacteria | Burkholderiales | Oxalobacteraceae |
| gi\|WP_188380807.1 | Oxalicibacterium faecigallinarum | Betaproteobacteria | Burkholderiales | Oxalobacteraceae |
| gi\|WP_186922539.1 | Undibacterium seohonense | Betaproteobacteria | Burkholderiales | Oxalobacteraceae |
| gi\|RZI42577.1 | Herbaspirillum sp. HC18 | Betaproteobacteria | Burkholderiales | Oxalobacteraceae |
| gi\|WP_161075125.1 | 'Massilia aquatica' Lu et al. 2020 | Betaproteobacteria | Burkholderiales | Oxalobacteraceae |
| gi\|SFU80941.1 | Massilia namucuonensis | Betaproteobacteria | Burkholderiales | Oxalobacteraceae |
| gi\|KAB8044350.1 | Janthinobacterium sp. FT58W | Betaproteobacteria | Burkholderiales | Oxalobacteraceae |
| gi\|RZU22588.1 | Duganella sp. BK054 | Betaproteobacteria | Burkholderiales | Oxalobacteraceae |
| gi\|WP_093386923.1 | Rugamonas rubra | Betaproteobacteria | Burkholderiales | Oxalobacteraceae |
| gi\|WP_134387719.1 | Massilia plicata | Betaproteobacteria | Burkholderiales | Oxalobacteraceae |
| gi\|WP_010398533.1 | Janthinobacterium lividum | Betaproteobacteria | Burkholderiales | Oxalobacteraceae |
| gi\|WP_038492362.1 | Janthinobacterium agaricidamnosum | Betaproteobacteria | Burkholderiales | Oxalobacteraceae |
| gi\|WP_182153161.1 | Duganella sp. LX47W | Betaproteobacteria | Burkholderiales | Oxalobacteraceae |
| gi\|WP_192049088.1 | unclassified Massilia | Betaproteobacteria | Burkholderiales | Oxalobacteraceae |
| gi\|WP_155472122.1 | Massilia buxea | Betaproteobacteria | Burkholderiales | Oxalobacteraceae |
| gi\|WP_169433872.1 | Duganella sp. GN2-R2 | Betaproteobacteria | Burkholderiales | Oxalobacteraceae |
| gi\|WP_051566847.1 | Herminiimonas sp. CN | Betaproteobacteria | Burkholderiales | Oxalobacteraceae |
| gi\|WP_155707829.1 | Massilia dura | Betaproteobacteria | Burkholderiales | Oxalobacteraceae |
| gi\|WP_186913797.1 | Undibacterium jejuense | Betaproteobacteria | Burkholderiales | Oxalobacteraceae |
| gi\|TFW13883.1 | Massilia arenosa | Betaproteobacteria | Burkholderiales | Oxalobacteraceae |
| gi\|WP_154373519.1 | Duganella sp. FT92W | Betaproteobacteria | Burkholderiales | Oxalobacteraceae |
| gi\|WP_194711049.1 | Noviherbaspirillum soli | Betaproteobacteria | Burkholderiales | Oxalobacteraceae |
| gi\|WP_192054283.1 | unclassified Massilia | Betaproteobacteria | Burkholderiales | Oxalobacteraceae |
| gi\|WP_179672390.1 | Duganella sp. 1224 | Betaproteobacteria | Burkholderiales | Oxalobacteraceae |
| gi\|WP_162058969.1 | Undibacterium sp. KW1 | Betaproteobacteria | Burkholderiales | Oxalobacteraceae |
| gi\|QGZ42057.1 | Massilia flava | Betaproteobacteria | Burkholderiales | Oxalobacteraceae |
| gi\|WP_057291959.1 | Noviherbaspirillum sp. Root189 | Betaproteobacteria | Burkholderiales | Oxalobacteraceae |
| gi\|WP_025916201.1 | Herminiimonas sp. CN | Betaproteobacteria | Burkholderiales | Oxalobacteraceae |
| gi\|WP_081466440.1 | Collimonas fungivorans | Betaproteobacteria | Burkholderiales | Oxalobacteraceae |
| gi\|TWI70103.1 | Massilia lurida | Betaproteobacteria | Burkholderiales | Oxalobacteraceae |
| gi\|WP_161021397.1 | Duganella sp. FT50W | Betaproteobacteria | Burkholderiales | Oxalobacteraceae |
| gi\|WP_182159962.1 | Duganella sp. LX20W | Betaproteobacteria | Burkholderiales | Oxalobacteraceae |
| gi\|WP_183738882.1 | unclassified Janthinobacterium | Betaproteobacteria | Burkholderiales | Oxalobacteraceae |
| gi\|WP_183381019.1 | unclassified Herbaspirillum | Betaproteobacteria | Burkholderiales | Oxalobacteraceae |
| gi\|WP_154378937.1 | Duganella sp. FT80W | Betaproteobacteria | Burkholderiales | Oxalobacteraceae |
| gi\|WP_094443883.1 | Janthinobacterium sp. PC23-8 | Betaproteobacteria | Burkholderiales | Oxalobacteraceae |
| gi\|WP_196856991.1 | Janthinobacterium sp. CAN_S1 | Betaproteobacteria | Burkholderiales | Oxalobacteraceae |
| gi\|WP_036244422.1 | Massilia sp. BSC265 | Betaproteobacteria | Burkholderiales | Oxalobacteraceae |
| gi\|WP_082491599.1 | Duganella sp. Leaf126 | Betaproteobacteria | Burkholderiales | Oxalobacteraceae |
| gi\|TFW33875.1 | Massilia horti | Betaproteobacteria | Burkholderiales | Oxalobacteraceae |
| gi\|WP_183743914.1 | unclassified Janthinobacterium | Betaproteobacteria | Burkholderiales | Oxalobacteraceae |
| gi\|WP_090182630.1 | unclassified Duganella | Betaproteobacteria | Burkholderiales | Oxalobacteraceae |
| gi\|WP_161084553.1 | Rugamonas sp. FT81W | Betaproteobacteria | Burkholderiales | Oxalobacteraceae |

| | | | | |
|---|---|---|---|---|
| gi\|WP_094443734.1 | Janthinobacterium sp. PC23-8 | Betaproteobacteria | Burkholderiales | Oxalobacteraceae |
| gi\|WP_186880895.1 | Undibacterium sp. CY7W | Betaproteobacteria | Burkholderiales | Oxalobacteraceae |
| gi\|AYR24596.1 | Herbaspirillum rubrisubalbicans | Betaproteobacteria | Burkholderiales | Oxalobacteraceae |
| gi\|WP_167092988.1 | Massilia frigida | Betaproteobacteria | Burkholderiales | Oxalobacteraceae |
| gi\|WP_159698671.1 | Massilia sp. 9I | Betaproteobacteria | Burkholderiales | Oxalobacteraceae |
| gi\|WP_161033688.1 | Duganella fentianensis | Betaproteobacteria | Burkholderiales | Oxalobacteraceae |
| gi\|WP_145881368.1 | Massilia flava | Betaproteobacteria | Burkholderiales | Oxalobacteraceae |
| gi\|WP_144736111.1 | Collimonas arenae | Betaproteobacteria | Burkholderiales | Oxalobacteraceae |
| gi\|WP_017876585.1 | Janthinobacterium sp. CG3 | Betaproteobacteria | Burkholderiales | Oxalobacteraceae |
| gi\|WP_161041188.1 | Pseudoduganella sp. CY13W | Betaproteobacteria | Burkholderiales | Oxalobacteraceae |
| gi\|WP_193686776.1 | Massilia sp. LPB0304 | Betaproteobacteria | Burkholderiales | Oxalobacteraceae |
| gi\|WP_151633559.1 | Noviherbaspirillum aerium | Betaproteobacteria | Burkholderiales | Oxalobacteraceae |
| gi\|SDF57215.1 | Duganella sp. OV458 | Betaproteobacteria | Burkholderiales | Oxalobacteraceae |
| gi\|WP_056337190.1 | Massilia sp. Leaf139 | Betaproteobacteria | Burkholderiales | Oxalobacteraceae |
| gi\|WP_090181395.1 | Duganella sp. OV510 | Betaproteobacteria | Burkholderiales | Oxalobacteraceae |
| gi\|WP_061533710.1 | Collimonas arenae | Betaproteobacteria | Burkholderiales | Oxalobacteraceae |
| gi\|OYO29839.1 | Janthinobacterium sp. PC23-8 | Betaproteobacteria | Burkholderiales | Oxalobacteraceae |
| gi\|WP_188394659.1 | Oxalicibacterium flavum | Betaproteobacteria | Burkholderiales | Oxalobacteraceae |
| gi\|WP_100872841.1 | Janthinobacterium sp. 64 | Betaproteobacteria | Burkholderiales | Oxalobacteraceae |
| gi\|WP_166100085.1 | Duganella aceris | Betaproteobacteria | Burkholderiales | Oxalobacteraceae |
| gi\|WP_183440428.1 | Massilia violacea | Betaproteobacteria | Burkholderiales | Oxalobacteraceae |
| gi\|WP_186897324.1 | Undibacterium sp. CY21W | Betaproteobacteria | Burkholderiales | Oxalobacteraceae |
| gi\|WP_166881848.1 | Massilia mucilaginosa | Betaproteobacteria | Burkholderiales | Oxalobacteraceae |
| gi\|WP_135204750.1 | Duganella callida | Betaproteobacteria | Burkholderiales | Oxalobacteraceae |
| gi\|WP_163962192.1 | Noviherbaspirillum galbum | Betaproteobacteria | Burkholderiales | Oxalobacteraceae |
| gi\|WP_116988117.1 | unclassified Duganella | Betaproteobacteria | Burkholderiales | Oxalobacteraceae |
| gi\|WP_102298675.1 | Janthinobacterium sp. AD80 | Betaproteobacteria | Burkholderiales | Oxalobacteraceae |
| gi\|WP_083438864.1 | Herbaspirillum autotrophicum | Betaproteobacteria | Burkholderiales | Oxalobacteraceae |
| gi\|SNT11187.1 | Noviherbaspirillum humi | Betaproteobacteria | Burkholderiales | Oxalobacteraceae |
| gi\|WP_130188186.1 | Massilia lutea | Betaproteobacteria | Burkholderiales | Oxalobacteraceae |
| gi\|WP_186913714.1 | Undibacterium jejuense | Betaproteobacteria | Burkholderiales | Oxalobacteraceae |
| gi\|WP_108441663.1 | Glaciimonas sp. PCH181 | Betaproteobacteria | Burkholderiales | Oxalobacteraceae |
| gi\|WP_081926099.1 | Massilia sp. LC238 | Betaproteobacteria | Burkholderiales | Oxalobacteraceae |
| gi\|SFD59486.1 | Massilia yuzhufengensis | Betaproteobacteria | Burkholderiales | Oxalobacteraceae |
| gi\|WP_161079006.1 | Duganella sp. CY15W | Betaproteobacteria | Burkholderiales | Oxalobacteraceae |
| gi\|WP_155710702.1 | Massilia dura | Betaproteobacteria | Burkholderiales | Oxalobacteraceae |
| gi\|WP_176348064.1 | Massilia sp. BJB1822 | Betaproteobacteria | Burkholderiales | Oxalobacteraceae |
| gi\|WP_160989134.1 | Duganella sp. FT94W | Betaproteobacteria | Burkholderiales | Oxalobacteraceae |
| gi\|RZT10620.1 | Duganella sp. BK701 | Betaproteobacteria | Burkholderiales | Oxalobacteraceae |
| gi\|WP_098497866.1 | Collimonas sp. PA-H2 | Betaproteobacteria | Burkholderiales | Oxalobacteraceae |
| gi\|WP_161041365.1 | Pseudoduganella sp. CY13W | Betaproteobacteria | Burkholderiales | Oxalobacteraceae |
| gi\|WP_167238121.1 | Massilia genomosp. 1 | Betaproteobacteria | Burkholderiales | Oxalobacteraceae |
| gi\|WP_162042114.1 | Undibacterium sp. YM2 | Betaproteobacteria | Burkholderiales | Oxalobacteraceae |

| | | | | |
|---|---|---|---|---|
| gi\|WP_008448257.1 | Janthinobacterium sp. HH01 | Betaproteobacteria | Burkholderiales | Oxalobacteraceae |
| gi\|WP_099414821.1 | Janthinobacterium sp. BJB412 | Betaproteobacteria | Burkholderiales | Oxalobacteraceae |
| gi\|WP_161096938.1 | Rugamonas sp. FT82W | Betaproteobacteria | Burkholderiales | Oxalobacteraceae |
| gi\|WP_126075803.1 | Massilia atriviolacea | Betaproteobacteria | Burkholderiales | Oxalobacteraceae |
| gi\|WP_126769906.1 | Undibacterium piscinae | Betaproteobacteria | Burkholderiales | Oxalobacteraceae |
| gi\|WP_034380466.1 | Herbaspirillum sp. CF444 | Betaproteobacteria | Burkholderiales | Oxalobacteraceae |
| gi\|WP_126072852.1 | Massilia atriviolacea | Betaproteobacteria | Burkholderiales | Oxalobacteraceae |
| gi\|WP_116990768.1 | unclassified Duganella | Betaproteobacteria | Burkholderiales | Oxalobacteraceae |
| gi\|WP_186885425.1 | Undibacterium sp. FT79W | Betaproteobacteria | Burkholderiales | Oxalobacteraceae |
| gi\|WP_034349387.1 | Noviherbaspirillum massiliense | Betaproteobacteria | Burkholderiales | Oxalobacteraceae |
| gi\|WP_159696088.1 | Massilia sp. 9I | Betaproteobacteria | Burkholderiales | Oxalobacteraceae |
| gi\|WP_152248044.1 | Janthinobacterium sp. FT58W | Betaproteobacteria | Burkholderiales | Oxalobacteraceae |
| gi\|WP_078032723.1 | Massilia sp. KIM | Betaproteobacteria | Burkholderiales | Oxalobacteraceae |
| gi\|WP_161052102.1 | Duganella sp. FT134W | Betaproteobacteria | Burkholderiales | Oxalobacteraceae |
| gi\|WP_189355856.1 | Undibacterium squillarum | Betaproteobacteria | Burkholderiales | Oxalobacteraceae |
| gi\|WP_092436557.1 | Collimonas sp. OK607 | Betaproteobacteria | Burkholderiales | Oxalobacteraceae |
| gi\|WP_177196801.1 | Duganella sp. CF517 | Betaproteobacteria | Burkholderiales | Oxalobacteraceae |
| gi\|WP_130189011.1 | Massilia lutea | Betaproteobacteria | Burkholderiales | Oxalobacteraceae |
| gi\|WP_099878295.1 | Massilia violaceinigra | Betaproteobacteria | Burkholderiales | Oxalobacteraceae |
| gi\|WP_161054368.1 | Duganella levis | Betaproteobacteria | Burkholderiales | Oxalobacteraceae |
| gi\|WP_061533678.1 | Collimonas arenae | Betaproteobacteria | Burkholderiales | Oxalobacteraceae |
| gi\|WP_162058970.1 | Undibacterium sp. KW1 | Betaproteobacteria | Burkholderiales | Oxalobacteraceae |
| gi\|WP_107140492.1 | Massilia armeniaca | Betaproteobacteria | Burkholderiales | Oxalobacteraceae |
| gi\|WP_065307629.1 | Janthinobacterium psychrotolerans | Betaproteobacteria | Burkholderiales | Oxalobacteraceae |
| gi\|WP_092417021.1 | Collimonas sp. OK307 | Betaproteobacteria | Burkholderiales | Oxalobacteraceae |
| gi\|WP_194722197.1 | Noviherbaspirillum malthae | Betaproteobacteria | Burkholderiales | Oxalobacteraceae |
| gi\|WP_176650569.1 | Rugamonas sp. SG757 | Betaproteobacteria | Burkholderiales | Oxalobacteraceae |
| gi\|WP_098493969.1 | Collimonas sp. PA-H2 | Betaproteobacteria | Burkholderiales | Oxalobacteraceae |
| gi\|WP_020654536.1 | Massilia niastensis | Betaproteobacteria | Burkholderiales | Oxalobacteraceae |
| gi\|WP_136220153.1 | Massilia sp. Mn16-1_5 | Betaproteobacteria | Burkholderiales | Oxalobacteraceae |
| gi\|WP_137316908.1 | Massilia umbonata | Betaproteobacteria | Burkholderiales | Oxalobacteraceae |
| gi\|WP_175048162.1 | Rugamonas sp. FT81W | Betaproteobacteria | Burkholderiales | Oxalobacteraceae |
| gi\|WP_099419087.1 | Janthinobacterium sp. BJB412 | Betaproteobacteria | Burkholderiales | Oxalobacteraceae |
| gi\|WP_166105972.1 | Duganella aceris | Betaproteobacteria | Burkholderiales | Oxalobacteraceae |
| gi\|WP_176650337.1 | Rugamonas sp. SG757 | Betaproteobacteria | Burkholderiales | Oxalobacteraceae |
| gi\|WP_094445851.1 | Janthinobacterium sp. PC23-8 | Betaproteobacteria | Burkholderiales | Oxalobacteraceae |
| gi\|WP_154133269.1 | unclassified Herbaspirillum | Betaproteobacteria | Burkholderiales | Oxalobacteraceae |
| gi\|WP_084320008.1 | Herbaspirillum huttiense | Betaproteobacteria | Burkholderiales | Oxalobacteraceae |
| gi\|WP_183108155.1 | Massilia sp. Dwa41.01b | Betaproteobacteria | Burkholderiales | Oxalobacteraceae |
| gi\|WP_161993179.1 | Lacisediminimonas profundi | Betaproteobacteria | Burkholderiales | Oxalobacteraceae |
| gi\|WP_183549687.1 | Massilia aurea | Betaproteobacteria | Burkholderiales | Oxalobacteraceae |
| gi\|WP_034294489.1 | Herbaspirillum sp. RV1423 | Betaproteobacteria | Burkholderiales | Oxalobacteraceae |
| gi\|CDG83654.1 | Janthinobacterium agaricidamnosum NBRC 102515 = DSM 9628 | Betaproteobacteria | Burkholderiales | Oxalobacteraceae |

| | | | | |
|---|---|---|---|---|
| gi\|WP_161035246.1 | Duganella fentianensis | Betaproteobacteria | Burkholderiales | Oxalobacteraceae |
| gi\|WP_112938799.1 | Massilia sp. YMA4 | Betaproteobacteria | Burkholderiales | Oxalobacteraceae |
| gi\|WP_099914755.1 | Massilia psychrophila | Betaproteobacteria | Burkholderiales | Oxalobacteraceae |
| gi\|KAF1042739.1 | Herbaspirillum frisingense | Betaproteobacteria | Burkholderiales | Oxalobacteraceae |
| gi\|CUI03361.1 | Janthinobacterium sp. CG23_2 | Betaproteobacteria | Burkholderiales | Oxalobacteraceae |
| gi\|WP_154356930.1 | Duganella rivi | Betaproteobacteria | Burkholderiales | Oxalobacteraceae |
| gi\|WP_184235603.1 | Massilia timonae | Betaproteobacteria | Burkholderiales | Oxalobacteraceae |
| gi\|WP_179672574.1 | Duganella sp. 1224 | Betaproteobacteria | Burkholderiales | Oxalobacteraceae |
| gi\|WP_161027447.1 | Massilia guangdongensis | Betaproteobacteria | Burkholderiales | Oxalobacteraceae |
| gi\|WP_154372889.1 | Duganella sp. FT80W | Betaproteobacteria | Burkholderiales | Oxalobacteraceae |
| gi\|WP_070246067.1 | Duganella phyllosphaerae | Betaproteobacteria | Burkholderiales | Oxalobacteraceae |
| gi\|WP_126127907.1 | Undibacterium parvum | Betaproteobacteria | Burkholderiales | Oxalobacteraceae |
| gi\|WP_186948530.1 | Undibacterium sp. CY18W | Betaproteobacteria | Burkholderiales | Oxalobacteraceae |
| gi\|PXX39990.1 | Undibacterium pigrum | Betaproteobacteria | Burkholderiales | Oxalobacteraceae |
| gi\|WP_110255060.1 | Undibacterium pigrum | Betaproteobacteria | Burkholderiales | Oxalobacteraceae |
| gi\|WP_119737153.1 | Herbaspirillum sp. K2R10-39 | Betaproteobacteria | Burkholderiales | Oxalobacteraceae |
| gi\|WP_182213586.1 | Duganella sp. FT3S | Betaproteobacteria | Burkholderiales | Oxalobacteraceae |
| gi\|WP_186955307.1 | Undibacterium sp. NL8W | Betaproteobacteria | Burkholderiales | Oxalobacteraceae |
| gi\|WP_183442683.1 | Massilia violacea | Betaproteobacteria | Burkholderiales | Oxalobacteraceae |
| gi\|WP_161079390.1 | Duganella sp. CY15W | Betaproteobacteria | Burkholderiales | Oxalobacteraceae |
| gi\|WP_144770279.1 | Herbaspirillum sp. SJZ099 | Betaproteobacteria | Burkholderiales | Oxalobacteraceae |
| gi\|WP_082221296.1 | Herbaspirillum chlorophenolicum | Betaproteobacteria | Burkholderiales | Oxalobacteraceae |
| gi\|WP_176648490.1 | Duganella sp. SG902 | Betaproteobacteria | Burkholderiales | Oxalobacteraceae |
| gi\|WP_090178746.1 | unclassified Duganella | Betaproteobacteria | Burkholderiales | Oxalobacteraceae |
| gi\|WP_137172245.1 | Massilia sp. HP4 | Betaproteobacteria | Burkholderiales | Oxalobacteraceae |
| gi\|WP_186952262.1 | Undibacterium sp. NL8W | Betaproteobacteria | Burkholderiales | Oxalobacteraceae |
| gi\|WP_135202042.1 | Duganella callida | Betaproteobacteria | Burkholderiales | Oxalobacteraceae |
| gi\|PUA20499.1 | Glaciimonas sp. PCH181 | Betaproteobacteria | Burkholderiales | Oxalobacteraceae |
| gi\|WP_008120300.1 | Herbaspirillum sp. YR522 | Betaproteobacteria | Burkholderiales | Oxalobacteraceae |
| gi\|TDK68672.1 | Sapientia aquatica | Betaproteobacteria | Burkholderiales | Oxalobacteraceae |
| gi\|WP_147936664.1 | Massilia sp. GEM5 | Betaproteobacteria | Burkholderiales | Oxalobacteraceae |
| gi\|WP_152882183.1 | Duganella sp. FT27W | Betaproteobacteria | Burkholderiales | Oxalobacteraceae |
| gi\|RYF21473.1 | Oxalobacteraceae bacterium | Betaproteobacteria | Burkholderiales | Oxalobacteraceae |
| gi\|WP_084416543.1 | Massilia alkalitolerans | Betaproteobacteria | Burkholderiales | Oxalobacteraceae |
| gi\|WP_186913719.1 | Undibacterium jejuense | Betaproteobacteria | Burkholderiales | Oxalobacteraceae |
| gi\|WP_186948145.1 | Undibacterium sp. CY18W | Betaproteobacteria | Burkholderiales | Oxalobacteraceae |
| gi\|WP_197034919.1 | Herbaspirillum sp. RV1423 | Betaproteobacteria | Burkholderiales | Oxalobacteraceae |
| gi\|WP_186863714.1 | Undibacterium sp. FT31W | Betaproteobacteria | Burkholderiales | Oxalobacteraceae |
| gi\|WP_124452023.1 | Paucibacter sp. KBW04 | Betaproteobacteria | Burkholderiales | Paucibacter |
| gi\|WP_124454914.1 | Paucibacter sp. KBW04 | Betaproteobacteria | Burkholderiales | Paucibacter |
| gi\|WP_124454936.1 | Paucibacter sp. KBW04 | Betaproteobacteria | Burkholderiales | Paucibacter |
| gi\|TDP71243.1 | Paucibacter toxinivorans | Betaproteobacteria | Burkholderiales | Paucibacter |
| gi\|WP_058718492.1 | Paucibacter sp. KCTC 42545 | Betaproteobacteria | Burkholderiales | Paucibacter |

| | | | | |
|---|---|---|---|---|
| gi\|WP_124454968.1 | Paucibacter sp. KBW04 | Betaproteobacteria | Burkholderiales | Paucibacter |
| gi\|TXC62160.1 | Piscinibacter aquaticus | Betaproteobacteria | Burkholderiales | Piscinibacter |
| gi\|WP_128000758.1 | Piscinibacter defluvii | Betaproteobacteria | Burkholderiales | Piscinibacter |
| gi\|TXC62134.1 | Piscinibacter aquaticus | Betaproteobacteria | Burkholderiales | Piscinibacter |
| gi\|WP_182661490.1 | Piscinibacter sp. SJAQ100 | Betaproteobacteria | Burkholderiales | Piscinibacter |
| gi\|WP_182663810.1 | Piscinibacter sp. SJAQ100 | Betaproteobacteria | Burkholderiales | Piscinibacter |
| gi\|TXC66947.1 | Piscinibacter aquaticus | Betaproteobacteria | Burkholderiales | Piscinibacter |
| gi\|WP_182662864.1 | Piscinibacter sp. SJAQ100 | Betaproteobacteria | Burkholderiales | Piscinibacter |
| gi\|WP_056806127.1 | unclassified Rhizobacter | Betaproteobacteria | Burkholderiales | Rhizobacter |
| gi\|WP_085749822.1 | Rhizobacter gummiphilus | Betaproteobacteria | Burkholderiales | Rhizobacter |
| gi\|WP_169669288.1 | Rhizobacter sp. SG490 | Betaproteobacteria | Burkholderiales | Rhizobacter |
| gi\|WP_083525952.1 | Roseateles depolymerans | Betaproteobacteria | Burkholderiales | Roseateles |
| gi\|WP_088451618.1 | Roseateles terrae | Betaproteobacteria | Burkholderiales | Roseateles |
| gi\|WP_058933724.1 | Roseateles depolymerans | Betaproteobacteria | Burkholderiales | Roseateles |
| gi\|WP_088449448.1 | Roseateles terrae | Betaproteobacteria | Burkholderiales | Roseateles |
| gi\|WP_143074051.1 | Roseateles sp. YR242 | Betaproteobacteria | Burkholderiales | Roseateles |
| gi\|WP_141100858.1 | Roseateles aquatilis | Betaproteobacteria | Burkholderiales | Roseateles |
| gi\|WP_164963571.1 | Rubrivivax sp. JA1026 | Betaproteobacteria | Burkholderiales | Rubrivivax |
| gi\|WP_196887490.1 | Rubrivivax gelatinosus | Betaproteobacteria | Burkholderiales | Rubrivivax |
| gi\|ODU10399.1 | Rubrivivax sp. SCN 71-131 | Betaproteobacteria | Burkholderiales | Rubrivivax |
| gi\|RZL02223.1 | Rubrivivax sp. | Betaproteobacteria | Burkholderiales | Rubrivivax |
| gi\|RVT51427.1 | Rubrivivax albus | Betaproteobacteria | Burkholderiales | Rubrivivax |
| gi\|RZI84896.1 | Rubrivivax sp. | Betaproteobacteria | Burkholderiales | Rubrivivax |
| gi\|WP_132646531.1 | Rubrivivax gelatinosus | Betaproteobacteria | Burkholderiales | Rubrivivax |
| gi\|WP_043784128.1 | Rubrivivax gelatinosus | Betaproteobacteria | Burkholderiales | Rubrivivax |
| gi\|WP_051632198.1 | Sphaerotilus natans | Betaproteobacteria | Burkholderiales | Sphaerotilus |
| gi\|WP_179635916.1 | Sphaerotilus montanus | Betaproteobacteria | Burkholderiales | Sphaerotilus |
| gi\|WP_133596046.1 | Tepidicella xavieri | Betaproteobacteria | Burkholderiales | Tepidicella |
| gi\|WP_180682922.1 | Tepidicella baoligensis | Betaproteobacteria | Burkholderiales | Tepidicella |
| gi\|WP_180682920.1 | Tepidicella baoligensis | Betaproteobacteria | Burkholderiales | Tepidicella |
| gi\|WP_043700633.1 | Tepidimonas taiwanensis | Betaproteobacteria | Burkholderiales | Tepidimonas |
| gi\|TSE29491.1 | Tepidimonas charontis | Betaproteobacteria | Burkholderiales | Tepidimonas |
| gi\|WP_185975033.1 | Tepidimonas thermarum | Betaproteobacteria | Burkholderiales | Tepidimonas |
| gi\|WP_082955436.1 | Tepidimonas fonticaldi | Betaproteobacteria | Burkholderiales | Tepidimonas |
| gi\|WP_185970649.1 | Tepidimonas sediminis | Betaproteobacteria | Burkholderiales | Tepidimonas |
| gi\|TSE21386.1 | Tepidimonas alkaliphilus | Betaproteobacteria | Burkholderiales | Tepidimonas |
| gi\|WP_082668345.1 | Tepidimonas taiwanensis | Betaproteobacteria | Burkholderiales | Tepidimonas |
| gi\|WP_143889630.1 | Tepidimonas alkaliphilus | Betaproteobacteria | Burkholderiales | Tepidimonas |
| gi\|TCS94578.1 | Tepidimonas ignava | Betaproteobacteria | Burkholderiales | Tepidimonas |
| gi\|OYU27763.1 | Burkholderiales bacterium PBB2 | Betaproteobacteria | Burkholderiales | |
| gi\|OYU29046.1 | Burkholderiales bacterium PBB2 | Betaproteobacteria | Burkholderiales | |
| gi\|RTL44221.1 | Burkholderiales bacterium | Betaproteobacteria | Burkholderiales | |
| gi\|OGA82381.1 | Burkholderiales bacterium RIFCSPHIGHO2_01_FULL_63_240 | Betaproteobacteria | Burkholderiales | |

| | | | | |
|---|---|---|---|---|
| gi\|OGB15922.1 | Burkholderiales bacterium RIFCSPLOWO2_02_FULL_67_64 | Betaproteobacteria | Burkholderiales | |
| gi\|OGA75887.1 | Burkholderiales bacterium GWE1_65_30 | Betaproteobacteria | Burkholderiales | |
| gi\|RTL29742.1 | Burkholderiales bacterium | Betaproteobacteria | Burkholderiales | |
| gi\|RPH67384.1 | Burkholderiales bacterium | Betaproteobacteria | Burkholderiales | |
| gi\|OGA81649.1 | Burkholderiales bacterium RIFCSPHIGHO2_01_FULL_63_240 | Betaproteobacteria | Burkholderiales | |
| gi\|OYU74626.1 | Burkholderiales bacterium PBB5 | Betaproteobacteria | Burkholderiales | |
| gi\|OYT85553.1 | Burkholderiales bacterium PBB6 | Betaproteobacteria | Burkholderiales | |
| gi\|OJX06329.1 | Burkholderiales bacterium 70-64 | Betaproteobacteria | Burkholderiales | |
| gi\|TAG69861.1 | Burkholderiales bacterium | Betaproteobacteria | Burkholderiales | |
| gi\|WP_035933172.1 | Burkholderiaceae | Betaproteobacteria | Burkholderiales | |
| gi\|OYY65265.1 | Burkholderiales bacterium 28-67-8 | Betaproteobacteria | Burkholderiales | |
| gi\|TNF58289.1 | Burkholderiales bacterium | Betaproteobacteria | Burkholderiales | |
| gi\|EHR73551.1 | Burkholderiales bacterium JOSHI_001 | Betaproteobacteria | Burkholderiales | |
| gi\|OYT98595.1 | Burkholderiales bacterium PBB1 | Betaproteobacteria | Burkholderiales | |
| gi\|OGB53080.1 | Burkholderiales bacterium RIFOXYD12_FULL_59_19 | Betaproteobacteria | Burkholderiales | |
| gi\|OJX31803.1 | Burkholderiales bacterium 68-12 | Betaproteobacteria | Burkholderiales | |
| gi\|EHR69429.1 | Burkholderiales bacterium JOSHI_001 | Betaproteobacteria | Burkholderiales | |
| gi\|OYV01006.1 | Burkholderiales bacterium PBB5 | Betaproteobacteria | Burkholderiales | |
| gi\|RTL18232.1 | Burkholderiales bacterium | Betaproteobacteria | Burkholderiales | |
| gi\|OGB30066.1 | Burkholderiales bacterium RIFCSPLOWO2_02_FULL_66_35 | Betaproteobacteria | Burkholderiales | |
| gi\|RPH67423.1 | Burkholderiales bacterium | Betaproteobacteria | Burkholderiales | |
| gi\|RTL18228.1 | Burkholderiales bacterium | Betaproteobacteria | Burkholderiales | |
| gi\|RTL31403.1 | Burkholderiales bacterium | Betaproteobacteria | Burkholderiales | |
| gi\|RTL18238.1 | Burkholderiales bacterium | Betaproteobacteria | Burkholderiales | |
| gi\|OYU27758.1 | Burkholderiales bacterium PBB2 | Betaproteobacteria | Burkholderiales | |
| gi\|OGB72329.1 | Burkholderiales bacterium RIFOXYC12_FULL_65_23 | Betaproteobacteria | Burkholderiales | |
| gi\|OGB05093.1 | Burkholderiales bacterium RIFCSPHIGHO2_12_FULL_63_20 | Betaproteobacteria | Burkholderiales | |
| gi\|OJX07574.1 | Burkholderiales bacterium 70-64 | Betaproteobacteria | Burkholderiales | |
| gi\|WP_156863106.1 | Casimicrobium huifangae | Betaproteobacteria | Casimicrobiaceae | Casimicrobium |
| gi\|WP_137938093.1 | Chitinivorax sp. B | Betaproteobacteria | Chitinivorax | unclassified Chitinivora |
| gi\|WP_137939466.1 | Chitinivorax sp. B | Betaproteobacteria | Chitinivorax | unclassified Chitinivora |
| gi\|WP_184041404.1 | Chitinivorax tropicus | Betaproteobacteria | Chitinivorax | |
| gi\|WP_026262865.1 | Chitiniphilus shinanonensis | Betaproteobacteria | Neisseriales | Chromobacteriaceae |
| gi\|WP_188704023.1 | Silvimonas iriomotensis | Betaproteobacteria | Neisseriales | Chromobacteriaceae |
| gi\|WP_188697799.1 | Silvimonas amylolytica | Betaproteobacteria | Neisseriales | Chromobacteriaceae |
| gi\|WP_084090514.1 | Andreprevotia lacus | Betaproteobacteria | Neisseriales | Chromobacteriaceae |
| gi\|WP_136772293.1 | Chitiniphilus eburneus | Betaproteobacteria | Neisseriales | Chromobacteriaceae |
| gi\|KAF0813755.1 | Andreprevotia sp. IGB-42 | Betaproteobacteria | Neisseriales | Chromobacteriaceae |
| gi\|ASM76636.1 | Vitreoscilla filiformis | Betaproteobacteria | Neisseriales | Neisseriaceae |
| gi\|WP_013029989.1 | Sideroxydans lithotrophicus | Betaproteobacteria | Nitrosomonadales | Gallionellaceae |
| gi\|ROH87198.1 | Pseudomethylobacillus aquaticus | Betaproteobacteria | Nitrosomonadales | Methylophilaceae |
| gi\|WP_067266913.1 | Methylovorus sp. MM2 | Betaproteobacteria | Nitrosomonadales | Methylophilaceae |
| gi\|WP_052661126.1 | Candidatus Methylopumilus turicensis | Betaproteobacteria | Nitrosomonadales | Methylophilaceae |

| | | | | |
|---|---|---|---|---|
| gi\|EUJ09795.1 | Methylophilaceae bacterium 11 | Betaproteobacteria | Nitrosomonadales | Methylophilaceae |
| gi\|WP_020167798.1 | Methylotenera | Betaproteobacteria | Nitrosomonadales | Methylophilaceae |
| gi\|GBL31825.1 | Methylophilaceae bacterium | Betaproteobacteria | Nitrosomonadales | Methylophilaceae |
| gi\|PCI59334.1 | Methylophilaceae bacterium | Betaproteobacteria | Nitrosomonadales | Methylophilaceae |
| gi\|PPC94488.1 | Methylotenera sp. | Betaproteobacteria | Nitrosomonadales | Methylophilaceae |
| gi\|WP_158497428.1 | Methylophilus sp. OH31 | Betaproteobacteria | Nitrosomonadales | Methylophilaceae |
| gi\|WP_029146837.1 | Methylophilus sp. 5 | Betaproteobacteria | Nitrosomonadales | Methylophilaceae |
| gi\|PPC94498.1 | Methylotenera sp. | Betaproteobacteria | Nitrosomonadales | Methylophilaceae |
| gi\|WP_046487682.1 | Candidatus Methylopumilus planktonicus | Betaproteobacteria | Nitrosomonadales | Methylophilaceae |
| gi\|WP_055827365.1 | unclassified Methylophilus | Betaproteobacteria | Nitrosomonadales | Methylophilaceae |
| gi\|WP_020182723.1 | unclassified Methylotenera | Betaproteobacteria | Nitrosomonadales | Methylophilaceae |
| gi\|WP_140002697.1 | Methylophilus medardicus | Betaproteobacteria | Nitrosomonadales | Methylophilaceae |
| gi\|WP_018230142.1 | Methyloversatilis universalis | Betaproteobacteria | Nitrosomonadales | Sterolibacteriaceae |
| gi\|WP_018411597.1 | Methyloversatilis thermotolerans | Betaproteobacteria | Nitrosomonadales | Sterolibacteriaceae |
| gi\|EGK73562.1 | Methyloversatilis universalis FAM5 | Betaproteobacteria | Nitrosomonadales | Sterolibacteriaceae |
| gi\|OYY48476.1 | Methylophilales bacterium 28-44-11 | Betaproteobacteria | Nitrosomonadales | |
| gi\|SIQ58056.1 | Aromatoleum tolulyticum | Betaproteobacteria | Rhodocyclales | Rhodocyclaceae |
| gi\|WP_168953661.1 | Aromatoleum aromaticum | Betaproteobacteria | Rhodocyclales | Rhodocyclaceae |
| gi\|WP_169260689.1 | Aromatoleum diolicum | Betaproteobacteria | Rhodocyclales | Rhodocyclaceae |
| gi\|AUL99697.1 | Rhodocyclaceae bacterium | Betaproteobacteria | Rhodocyclales | Rhodocyclaceae |
| gi\|WP_184414907.1 | Rhodocyclus tenuis | Betaproteobacteria | Rhodocyclales | Rhodocyclaceae |
| gi\|AJP47770.1 | Rugosibacter aromaticivorans | Betaproteobacteria | Rhodocyclales | Rhodocyclaceae |
| gi\|WP_183632782.1 | Niveibacterium umoris | Betaproteobacteria | Rhodocyclales | Rhodocyclaceae |
| gi\|WP_026688131.1 | Azovibrio restrictus | Betaproteobacteria | Rhodocyclales | Rhodocyclaceae |
| gi\|WP_172202311.1 | Niveibacterium sp. COAC-50 | Betaproteobacteria | Rhodocyclales | Rhodocyclaceae |
| gi\|OHC61696.1 | Rhodocyclales bacterium GWA2_65_19 | Betaproteobacteria | Rhodocyclales | unclassified Rhodocycla |
| gi\|OHC68299.1 | Rhodocyclales bacterium RIFCSPLOWO2_02_FULL_63_24 | Betaproteobacteria | Rhodocyclales | unclassified Rhodocycla |
| gi\|WP_011765888.1 | Azoarcus olearius | Betaproteobacteria | Rhodocyclales | Zoogloeaceae |
| gi\|BAL24361.1 | Azoarcus sp. KH32C | Betaproteobacteria | Rhodocyclales | Zoogloeaceae |
| gi\|WP_141018560.1 | Azoarcus sp. DD4 | Betaproteobacteria | Rhodocyclales | Zoogloeaceae |
| gi\|WP_043746796.1 | Thauera sp. SWB20 | Betaproteobacteria | Rhodocyclales | Zoogloeaceae |
| gi\|WP_002940923.1 | Thauera sp. 27 | Betaproteobacteria | Rhodocyclales | Zoogloeaceae |
| gi\|WP_002936577.1 | Thauera sp. 27 | Betaproteobacteria | Rhodocyclales | Zoogloeaceae |
| gi\|WP_028792260.1 | Thauera linaloolentis | Betaproteobacteria | Rhodocyclales | Zoogloeaceae |
| gi\|WP_094267991.1 | Thauera propionica | Betaproteobacteria | Rhodocyclales | Zoogloeaceae |
| gi\|WP_004365913.1 | Thauera phenylacetica | Betaproteobacteria | Rhodocyclales | Zoogloeaceae |
| gi\|WP_107221634.1 | Thauera aromatica | Betaproteobacteria | Rhodocyclales | Zoogloeaceae |
| gi\|WP_068807292.1 | Thauera phenolivorans | Betaproteobacteria | Rhodocyclales | Zoogloeaceae |
| gi\|WP_168941243.1 | Azoarcus communis | Betaproteobacteria | Rhodocyclales | Zoogloeaceae |
| gi\|WP_136385116.1 | Azoarcus rhizosphaerae | Betaproteobacteria | Rhodocyclales | Zoogloeaceae |
| gi\|WP_021250616.1 | Thauera terpenica | Betaproteobacteria | Rhodocyclales | Zoogloeaceae |
| gi\|WP_108975498.1 | Azoarcus communis | Betaproteobacteria | Rhodocyclales | Zoogloeaceae |
| gi\|TDN48108.1 | Azoarcus indigens | Betaproteobacteria | Rhodocyclales | Zoogloeaceae |

| | | | | |
|---|---|---|---|---|
| gi\|THF67256.1 | Azoarcus nasutitermitis | Betaproteobacteria | Rhodocyclales | Zoogloeaceae |
| gi\|WP_075147558.1 | Thauera chlorobenzoica | Betaproteobacteria | Rhodocyclales | Zoogloeaceae |
| gi\|WP_187717606.1 | Thauera sp. CAU 1555 | Betaproteobacteria | Rhodocyclales | Zoogloeaceae |
| gi\|WP_168989604.1 | Azoarcus taiwanensis | Betaproteobacteria | Rhodocyclales | Zoogloeaceae |
| gi\|PTD98165.1 | Thauera sp. D20 | Betaproteobacteria | Rhodocyclales | Zoogloeaceae |
| gi\|WP_136346521.1 | Azoarcus nasutitermitis | Betaproteobacteria | Rhodocyclales | Zoogloeaceae |
| gi\|WP_004339014.1 | Thauera linaloolentis | Betaproteobacteria | Rhodocyclales | Zoogloeaceae |
| gi\|WP_114649711.1 | Thauera hydrothermalis | Betaproteobacteria | Rhodocyclales | Zoogloeaceae |
| gi\|WP_187717610.1 | Thauera sp. CAU 1555 | Betaproteobacteria | Rhodocyclales | Zoogloeaceae |
| gi\|WP_141018080.1 | Azoarcus sp. DD4 | Betaproteobacteria | Rhodocyclales | Zoogloeaceae |
| gi\|WP_136346523.1 | Azoarcus nasutitermitis | Betaproteobacteria | Rhodocyclales | Zoogloeaceae |
| gi\|WP_169152940.1 | Azoarcus sp. TTM-91 | Betaproteobacteria | Rhodocyclales | Zoogloeaceae |
| gi\|WP_096446429.1 | Thauera sp. K11 | Betaproteobacteria | Rhodocyclales | Zoogloeaceae |
| gi\|WP_173767198.1 | Azoarcus sp. M9-3-2 | Betaproteobacteria | Rhodocyclales | Zoogloeaceae |
| gi\|QDF97508.1 | Azoarcus sp. DD4 | Betaproteobacteria | Rhodocyclales | Zoogloeaceae |
| gi\|WP_169149817.1 | Azoarcus sp. TTM-91 | Betaproteobacteria | Rhodocyclales | Zoogloeaceae |
| gi\|WP_018610353.1 | Uliginosibacterium gangwonense | Betaproteobacteria | Rhodocyclales | Zoogloeaceae |
| gi\|PKO59351.1 | Betaproteobacteria bacterium HGW-Betaproteobacteria-19 | Betaproteobacteria | | |
| gi\|KPF48170.1 | beta proteobacterium AAP65 | Betaproteobacteria | | |
| gi\|PKO59341.1 | Betaproteobacteria bacterium HGW-Betaproteobacteria-19 | Betaproteobacteria | | |
| gi\|TMG99601.1 | Betaproteobacteria bacterium | Betaproteobacteria | | |
| gi\|PRY99332.1 | beta proteobacterium MWH-P2sevCIIIb | Betaproteobacteria | | |
| gi\|PKO67258.1 | Betaproteobacteria bacterium HGW-Betaproteobacteria-16 | Betaproteobacteria | | |
| gi\|TMG85304.1 | Betaproteobacteria bacterium | Betaproteobacteria | | |
| gi\|OAI52156.1 | Betaproteobacteria bacterium SCGC AG-212-J23 | Betaproteobacteria | | |
| gi\|TMH63601.1 | Betaproteobacteria bacterium | Betaproteobacteria | | |
| gi\|TMH28417.1 | Betaproteobacteria bacterium | Betaproteobacteria | | |
| gi\|WP_094201249.1 | Oceanimonas doudoroffii | Gammaproteobacteria | Aeromonadales | Aeromonadaceae |
| gi\|PSJ44474.1 | Zobellella endophytica | Gammaproteobacteria | Aeromonadales | Aeromonadaceae |
| gi\|WP_165855923.1 | Marinobacter sp. JSM 1782161 | Gammaproteobacteria | Alteromonadales | Alteromonadaceae |
| gi\|WP_121206455.1 | Marinobacter hydrocarbonoclasticus | Gammaproteobacteria | Alteromonadales | Alteromonadaceae |
| gi\|WP_123635898.1 | Marinobacter sp. R17 | Gammaproteobacteria | Alteromonadales | Alteromonadaceae |
| gi\|WP_004580561.1 | Marinobacter nanhaiticus | Gammaproteobacteria | Alteromonadales | Alteromonadaceae |
| gi\|WP_138437567.1 | Marinobacter shengliensis | Gammaproteobacteria | Alteromonadales | Alteromonadaceae |
| gi\|WP_178380779.1 | Marinobacter sp. C18 | Gammaproteobacteria | Alteromonadales | Alteromonadaceae |
| gi\|WP_168203191.1 | Marinobacter fonticola | Gammaproteobacteria | Alteromonadales | Alteromonadaceae |
| gi\|WP_136548779.1 | Hydrocarboniclastica marina | Gammaproteobacteria | Alteromonadales | Alteromonadaceae |
| gi\|WP_012638513.1 | Thioalkalivibrio sulfidiphilus | Gammaproteobacteria | Chromatiales | Ectothiorhodospiraceae |
| gi\|WP_116302778.1 | Alkalilimnicola ehrlichii | Gammaproteobacteria | Chromatiales | Ectothiorhodospiraceae |
| gi\|WP_132924724.1 | Sodalis sp. 159R | Gammaproteobacteria | Enterobacterales | Bruguierivoracaceae |
| gi\|WP_131865920.1 | Biostraticola tofi | Gammaproteobacteria | Enterobacterales | Bruguierivoracaceae |
| gi\|WP_111741394.1 | Leminorella richardii | Gammaproteobacteria | Enterobacterales | Budviciaceae |
| gi\|WP_022548345.1 | Plautia stali symbiont | Gammaproteobacteria | Enterobacterales | Enterobacteriaceae |

| | | | | |
|---|---|---|---|---|
| gi\|STE15749.1 | Escherichia coli | Gammaproteobacteria | Enterobacterales | Enterobacteriaceae |
| gi\|WP_061707521.1 | Enterobacter timonensis | Gammaproteobacteria | Enterobacterales | Enterobacteriaceae |
| gi\|SUX60525.1 | Citrobacter koseri | Gammaproteobacteria | Enterobacterales | Enterobacteriaceae |
| gi\|ABP61750.1 | Enterobacter sp. 638 | Gammaproteobacteria | Enterobacterales | Enterobacteriaceae |
| gi\|AML39077.1 | Klebsiella aerogenes | Gammaproteobacteria | Enterobacterales | Enterobacteriaceae |
| gi\|WP_064374013.1 | Klebsiella oxytoca | Gammaproteobacteria | Enterobacterales | Enterobacteriaceae |
| gi\|WP_103950181.1 | Lelliottia | Gammaproteobacteria | Enterobacterales | Enterobacteriaceae |
| gi\|KLV66025.1 | Citrobacter sp. MGH106 | Gammaproteobacteria | Enterobacterales | Enterobacteriaceae |
| gi\|WP_086626519.1 | Enterobacter hormaechei | Gammaproteobacteria | Enterobacterales | Enterobacteriaceae |
| gi\|WP_161617585.1 | Yokenella regensburgei | Gammaproteobacteria | Enterobacterales | Enterobacteriaceae |
| gi\|WP_097164669.1 | Enterobacter sp. CC120223-11 | Gammaproteobacteria | Enterobacterales | Enterobacteriaceae |
| gi\|WP_182240489.1 | Klebsiella sp. RHBSTW-00215 | Gammaproteobacteria | Enterobacterales | Enterobacteriaceae |
| gi\|WP_110512058.1 | Scandinavium goeteborgense | Gammaproteobacteria | Enterobacterales | Enterobacteriaceae |
| gi\|WP_090465037.1 | Enterobacter sp. kpr-6 | Gammaproteobacteria | Enterobacterales | Enterobacteriaceae |
| gi\|WP_142470317.1 | Klebsiella pasteurii | Gammaproteobacteria | Enterobacterales | Enterobacteriaceae |
| gi\|KMV35403.1 | Franconibacter pulveris | Gammaproteobacteria | Enterobacterales | Enterobacteriaceae |
| gi\|WP_072571528.1 | Enterobacter sp. SA187 | Gammaproteobacteria | Enterobacterales | Enterobacteriaceae |
| gi\|WP_142465741.1 | Klebsiella spallanzanii | Gammaproteobacteria | Enterobacterales | Enterobacteriaceae |
| gi\|WP_041146787.1 | Raoultella ornithinolytica | Gammaproteobacteria | Enterobacterales | Enterobacteriaceae |
| gi\|WP_126509445.1 | Raoultella ornithinolytica | Gammaproteobacteria | Enterobacterales | Enterobacteriaceae |
| gi\|WP_192478508.1 | Citrobacter amalonaticus | Gammaproteobacteria | Enterobacterales | Enterobacteriaceae |
| gi\|WP_064564436.1 | Kosakonia oryzae | Gammaproteobacteria | Enterobacterales | Enterobacteriaceae |
| gi\|WP_142486766.1 | Leclercia adecarboxylata | Gammaproteobacteria | Enterobacterales | Enterobacteriaceae |
| gi\|BBV64796.1 | Klebsiella sp. STW0522-44 | Gammaproteobacteria | Enterobacterales | Enterobacteriaceae |
| gi\|WP_154681583.1 | Klebsiella oxytoca | Gammaproteobacteria | Enterobacterales | Enterobacteriaceae |
| gi\|KNC06045.1 | Klebsiella sp. RIT-PI-d | Gammaproteobacteria | Enterobacterales | Enterobacteriaceae |
| gi\|WP_149461728.1 | Pseudocitrobacter sp. 73 | Gammaproteobacteria | Enterobacterales | Enterobacteriaceae |
| gi\|WP_110277458.1 | Klebsiella oxytoca | Gammaproteobacteria | Enterobacterales | Enterobacteriaceae |
| gi\|WP_161660932.1 | Atlantibacter hermannii | Gammaproteobacteria | Enterobacterales | Enterobacteriaceae |
| gi\|WP_044711903.1 | Citrobacter freundii | Gammaproteobacteria | Enterobacterales | Enterobacteriaceae |
| gi\|QLK61991.1 | Enterobacteriaceae bacterium Kacie_13 | Gammaproteobacteria | Enterobacterales | Enterobacteriaceae |
| gi\|WP_062741724.1 | [Enterobacter] lignolyticus | Gammaproteobacteria | Enterobacterales | Enterobacteriaceae |
| gi\|WP_121265572.1 | Enterobacter sp. R1(2018) | Gammaproteobacteria | Enterobacterales | Enterobacteriaceae |
| gi\|WP_123349958.1 | unclassified Enterobacter | Gammaproteobacteria | Enterobacterales | Enterobacteriaceae |
| gi\|OAT24617.1 | Buttiauxella ferragutiae ATCC 51602 | Gammaproteobacteria | Enterobacterales | Enterobacteriaceae |
| gi\|BBQ82702.1 | Klebsiella sp. WP3-W18-ESBL-02 | Gammaproteobacteria | Enterobacterales | Enterobacteriaceae |
| gi\|VDZ82079.1 | Kluyvera intermedia | Gammaproteobacteria | Enterobacterales | Enterobacteriaceae |
| gi\|KGB02884.1 | Enterobacteriaceae bacterium ATCC 29904 | Gammaproteobacteria | Enterobacterales | Enterobacteriaceae |
| gi\|WP_123915032.1 | Citrobacter europaeus | Gammaproteobacteria | Enterobacterales | Enterobacteriaceae |
| gi\|WP_034813585.1 | Enterobacter cloacae | Gammaproteobacteria | Enterobacterales | Enterobacteriaceae |
| gi\|WP_081653630.1 | Metakosakonia massiliensis | Gammaproteobacteria | Enterobacterales | Enterobacteriaceae |
| gi\|WP_165463485.1 | Citrobacter freundii | Gammaproteobacteria | Enterobacterales | Enterobacteriaceae |
| gi\|WP_061493615.1 | Kosakonia oryzendophytica | Gammaproteobacteria | Enterobacterales | Enterobacteriaceae |

| | | | | |
|---|---|---|---|---|
| gi\|WP_032983496.1 | Cronobacter malonaticus | Gammaproteobacteria | Enterobacterales | Enterobacteriaceae |
| gi\|WP_082022442.1 | Enterobacter sp. Bisph1 | Gammaproteobacteria | Enterobacterales | Enterobacteriaceae |
| gi\|KSY31794.1 | Citrobacter sp. 50677481 | Gammaproteobacteria | Enterobacterales | Enterobacteriaceae |
| gi\|VDZ74679.1 | Atlantibacter hermannii | Gammaproteobacteria | Enterobacterales | Enterobacteriaceae |
| gi\|OAT55241.1 | Kluyvera georgiana ATCC 51603 | Gammaproteobacteria | Enterobacterales | Enterobacteriaceae |
| gi\|WP_165501944.1 | Kosakonia quasisacchari | Gammaproteobacteria | Enterobacterales | Enterobacteriaceae |
| gi\|VFS63826.1 | Kluyvera cryocrescens | Gammaproteobacteria | Enterobacterales | Enterobacteriaceae |
| gi\|WP_124023108.1 | Buttiauxella warmboldiae | Gammaproteobacteria | Enterobacterales | Enterobacteriaceae |
| gi\|WP_039293084.1 | Cedecea neteri | Gammaproteobacteria | Enterobacterales | Enterobacteriaceae |
| gi\|WP_110876859.1 | Franconibacter helveticus | Gammaproteobacteria | Enterobacterales | Enterobacteriaceae |
| gi\|KEA52246.1 | Mangrovibacter sp. MFB070 | Gammaproteobacteria | Enterobacterales | Enterobacteriaceae |
| gi\|WP_172731001.1 | Pluralibacter gergoviae | Gammaproteobacteria | Enterobacterales | Enterobacteriaceae |
| gi\|VDR30317.1 | Raoultella terrigena | Gammaproteobacteria | Enterobacterales | Enterobacteriaceae |
| gi\|WP_138099316.1 | Jejubacter calystegiae | Gammaproteobacteria | Enterobacterales | Enterobacteriaceae |
| gi\|ASG62891.1 | Kluyvera genomosp. 3 | Gammaproteobacteria | Enterobacterales | Enterobacteriaceae |
| gi\|AUP76406.1 | Enterobacter sp. EA-1 | Gammaproteobacteria | Enterobacterales | Enterobacteriaceae |
| gi\|WP_130099291.1 | Siccibacter turicensis | Gammaproteobacteria | Enterobacterales | Enterobacteriaceae |
| gi\|WP_086499451.1 | Pluralibacter gergoviae | Gammaproteobacteria | Enterobacterales | Enterobacteriaceae |
| gi\|WP_114262800.1 | Klebsiella pneumoniae | Gammaproteobacteria | Enterobacterales | Enterobacteriaceae |
| gi\|WP_075203725.1 | Citrobacter koseri | Gammaproteobacteria | Enterobacterales | Enterobacteriaceae |
| gi\|WP_039295443.1 | Cedecea | Gammaproteobacteria | Enterobacterales | Enterobacteriaceae |
| gi\|WP_039898482.1 | Cedecea | Gammaproteobacteria | Enterobacterales | Enterobacteriaceae |
| gi\|WP_082031763.1 | Citrobacter | Gammaproteobacteria | Enterobacterales | Enterobacteriaceae |
| gi\|SFD19347.1 | Kosakonia oryzae | Gammaproteobacteria | Enterobacterales | Enterobacteriaceae |
| gi\|WP_064569327.1 | Klebsiella aerogenes | Gammaproteobacteria | Enterobacterales | Enterobacteriaceae |
| gi\|AOV15511.1 | Klebsiella sp. LTGPAF-6F | Gammaproteobacteria | Enterobacterales | Enterobacteriaceae |
| gi\|KHJ68471.1 | Pantoea rodasii | Gammaproteobacteria | Enterobacterales | Erwiniaceae |
| gi\|KJV35228.1 | Pantoea sp. SM3 | Gammaproteobacteria | Enterobacterales | Erwiniaceae |
| gi\|WP_120455994.1 | Kalamiella piersonii | Gammaproteobacteria | Enterobacterales | Erwiniaceae |
| gi\|WP_133842425.1 | Erwinia rhapontici | Gammaproteobacteria | Enterobacterales | Erwiniaceae |
| gi\|QGY31513.1 | Pantoea cypripedii | Gammaproteobacteria | Enterobacterales | Erwiniaceae |
| gi\|WP_110331991.1 | Pantoea sp. JKS000250 | Gammaproteobacteria | Enterobacterales | Erwiniaceae |
| gi\|WP_040113679.1 | Pantoea | Gammaproteobacteria | Enterobacterales | Erwiniaceae |
| gi\|CCF12026.1 | Pantoea ananatis LMG 5342 | Gammaproteobacteria | Enterobacterales | Erwiniaceae |
| gi\|WP_017800976.1 | Erwinia toletana | Gammaproteobacteria | Enterobacterales | Erwiniaceae |
| gi\|WP_163638822.1 | Pantoea agglomerans | Gammaproteobacteria | Enterobacterales | Erwiniaceae |
| gi\|WP_052901575.1 | Erwinia iniecta | Gammaproteobacteria | Enterobacterales | Erwiniaceae |
| gi\|WP_123802325.1 | Pantoea sp. RIT388 | Gammaproteobacteria | Enterobacterales | Erwiniaceae |
| gi\|WP_145891349.1 | Pantoea dispersa | Gammaproteobacteria | Enterobacterales | Erwiniaceae |
| gi\|EXU74429.1 | Erwinia mallotivora | Gammaproteobacteria | Enterobacterales | Erwiniaceae |
| gi\|WP_034950256.1 | Erwinia oleae | Gammaproteobacteria | Enterobacterales | Erwiniaceae |
| gi\|WP_094119074.1 | Pantoea conspicua | Gammaproteobacteria | Enterobacterales | Erwiniaceae |
| gi\|QKJ87136.1 | Erwiniaceae bacterium PD-1 | Gammaproteobacteria | Enterobacterales | Erwiniaceae |

| | | | | |
|---|---|---|---|---|
| gi\|WP_081141819.1 | Pantoea latae | Gammaproteobacteria | Enterobacterales | Erwiniaceae |
| gi\|WP_147200508.1 | Pantoea sp. CCBC3-3-1 | Gammaproteobacteria | Enterobacterales | Erwiniaceae |
| gi\|WP_072055931.1 | Tatumella morbirosei | Gammaproteobacteria | Enterobacterales | Erwiniaceae |
| gi\|WP_193406167.1 | Mixta mediterraneensis | Gammaproteobacteria | Enterobacterales | Erwiniaceae |
| gi\|WP_046289294.1 | Pantoea | Gammaproteobacteria | Enterobacterales | Erwiniaceae |
| gi\|WP_103061260.1 | Mixta theicola | Gammaproteobacteria | Enterobacterales | Erwiniaceae |
| gi\|CAX58097.1 | Erwinia billingiae Eb661 | Gammaproteobacteria | Enterobacterales | Erwiniaceae |
| gi\|WP_111207164.1 | Pantoea sp. ARC270 | Gammaproteobacteria | Enterobacterales | Erwiniaceae |
| gi\|WP_158782749.1 | Pantoea sp. BAV 3049 | Gammaproteobacteria | Enterobacterales | Erwiniaceae |
| gi\|TDS67801.1 | Pantoea sp. PNA 14-12 | Gammaproteobacteria | Enterobacterales | Erwiniaceae |
| gi\|WP_156287964.1 | Erwinia sp. J780 | Gammaproteobacteria | Enterobacterales | Erwiniaceae |
| gi\|WP_187485943.1 | Erwinia gerundensis | Gammaproteobacteria | Enterobacterales | Erwiniaceae |
| gi\|WP_017801779.1 | Erwinia toletana | Gammaproteobacteria | Enterobacterales | Erwiniaceae |
| gi\|WP_075183084.1 | Pantoea sp. 1.19 | Gammaproteobacteria | Enterobacterales | Erwiniaceae |
| gi\|WP_125290429.1 | Erwinia sp. 198 | Gammaproteobacteria | Enterobacterales | Erwiniaceae |
| gi\|WP_067704617.1 | Erwinia sp. ErVv1 | Gammaproteobacteria | Enterobacterales | Erwiniaceae |
| gi\|WP_105594736.1 | Pantoea coffeiphila | Gammaproteobacteria | Enterobacterales | Erwiniaceae |
| gi\|WP_130835814.1 | Erwinia mediterraneensis | Gammaproteobacteria | Enterobacterales | Erwiniaceae |
| gi\|WP_051434224.1 | Phaseolibacter flectens | Gammaproteobacteria | Enterobacterales | Erwiniaceae |
| gi\|WP_171149938.1 | Erwinia sp. JH02 | Gammaproteobacteria | Enterobacterales | Erwiniaceae |
| gi\|QHM73880.1 | Mixta intestinalis | Gammaproteobacteria | Enterobacterales | Erwiniaceae |
| gi\|WP_152322711.1 | Erwinia endophytica | Gammaproteobacteria | Enterobacterales | Erwiniaceae |
| gi\|WP_168428371.1 | Erwinia amylovora | Gammaproteobacteria | Enterobacterales | Erwiniaceae |
| gi\|WP_124234461.1 | Erwinia psidii | Gammaproteobacteria | Enterobacterales | Erwiniaceae |
| gi\|WP_017348734.1 | Pantoea sp. A4 | Gammaproteobacteria | Enterobacterales | Erwiniaceae |
| gi\|WP_167017124.1 | Pantoea sp. Acro-835 | Gammaproteobacteria | Enterobacterales | Erwiniaceae |
| gi\|PRD15486.1 | Pantoea coffeiphila | Gammaproteobacteria | Enterobacterales | Erwiniaceae |
| gi\|WP_177173132.1 | Rosenbergiella nectarea | Gammaproteobacteria | Enterobacterales | Erwiniaceae |
| gi\|WP_099754099.1 | Pantoea Psp39-30 | Gammaproteobacteria | Enterobacterales | Erwiniaceae |
| gi\|WP_192841080.1 | Pantoea sp. A4 | Gammaproteobacteria | Enterobacterales | Erwiniaceae |
| gi\|WP_190296477.1 | Mixta calida | Gammaproteobacteria | Enterobacterales | Erwiniaceae |
| gi\|WP_167373281.1 | Pantoea alhagi | Gammaproteobacteria | Enterobacterales | Erwiniaceae |
| gi\|WP_034895815.1 | Erwinia typographi | Gammaproteobacteria | Enterobacterales | Erwiniaceae |
| gi\|CBJ46452.1 | Erwinia amylovora ATCC 49946 | Gammaproteobacteria | Enterobacterales | Erwiniaceae |
| gi\|WP_048697774.1 | Erwinia piriflorinigrans | Gammaproteobacteria | Enterobacterales | Erwiniaceae |
| gi\|WP_023655097.1 | Erwinia piriflorinigrans | Gammaproteobacteria | Enterobacterales | Erwiniaceae |
| gi\|WP_137268576.1 | Erwinia persicina | Gammaproteobacteria | Enterobacterales | Erwiniaceae |
| gi\|TDT02413.1 | Erwinia rhapontici | Gammaproteobacteria | Enterobacterales | Erwiniaceae |
| gi\|WP_078001331.1 | Izhakiella australiensis | Gammaproteobacteria | Enterobacterales | Erwiniaceae |
| gi\|WP_193406228.1 | Mixta mediterraneensis | Gammaproteobacteria | Enterobacterales | Erwiniaceae |
| gi\|WP_160250803.1 | Mixta theicola | Gammaproteobacteria | Enterobacterales | Erwiniaceae |
| gi\|WP_087489758.1 | Tatumella citrea | Gammaproteobacteria | Enterobacterales | Erwiniaceae |
| gi\|WP_012440351.1 | Erwinia tasmaniensis | Gammaproteobacteria | Enterobacterales | Erwiniaceae |

| | | | | |
|---|---|---|---|---|
| gi\|WP_092677667.1 | Rosenbergiella nectarea | Gammaproteobacteria | Enterobacterales | Erwiniaceae |
| gi\|WP_051124374.1 | Erwinia tracheiphila | Gammaproteobacteria | Enterobacterales | Erwiniaceae |
| gi\|WP_191933084.1 | Erwinia persicina | Gammaproteobacteria | Enterobacterales | Erwiniaceae |
| gi\|WP_154325535.1 | Pantoea sp. 201603H | Gammaproteobacteria | Enterobacterales | Erwiniaceae |
| gi\|WP_034892104.1 | Erwinia typographi | Gammaproteobacteria | Enterobacterales | Erwiniaceae |
| gi\|WP_157725262.1 | Tatumella sp. TA1 | Gammaproteobacteria | Enterobacterales | Erwiniaceae |
| gi\|WP_104951623.1 | Mixta calida | Gammaproteobacteria | Enterobacterales | Erwiniaceae |
| gi\|WP_158239892.1 | Erwinia sp. B116 | Gammaproteobacteria | Enterobacterales | Erwiniaceae |
| gi\|WP_152540239.1 | Pantoea sp. IMH | Gammaproteobacteria | Enterobacterales | Erwiniaceae |
| gi\|WP_123333178.1 | Erwinia sp. JUb26 | Gammaproteobacteria | Enterobacterales | Erwiniaceae |
| gi\|WP_188474503.1 | Hafnia psychrotolerans | Gammaproteobacteria | Enterobacterales | Hafniaceae |
| gi\|ETS33373.1 | Photorhabdus khanii NC19 | Gammaproteobacteria | Enterobacterales | Morganellaceae |
| gi\|SCZ55504.1 | Photorhabdus luminescens | Gammaproteobacteria | Enterobacterales | Morganellaceae |
| gi\|OTA19922.1 | Xenorhabdus beddingii | Gammaproteobacteria | Enterobacterales | Morganellaceae |
| gi\|WP_081990776.1 | Xenorhabdus nematophila | Gammaproteobacteria | Enterobacterales | Morganellaceae |
| gi\|CDH21567.1 | Xenorhabdus bovienii str. kraussei Quebec | Gammaproteobacteria | Enterobacterales | Morganellaceae |
| gi\|WP_092508758.1 | Xenorhabdus mauleonii | Gammaproteobacteria | Enterobacterales | Morganellaceae |
| gi\|KLU15747.1 | Xenorhabdus griffiniae | Gammaproteobacteria | Enterobacterales | Morganellaceae |
| gi\|TYP03620.1 | Xenorhabdus doucetiae | Gammaproteobacteria | Enterobacterales | Morganellaceae |
| gi\|CDG21391.1 | Xenorhabdus poinarii G6 | Gammaproteobacteria | Enterobacterales | Morganellaceae |
| gi\|WP_084616090.1 | Xenorhabdus szentirmaii | Gammaproteobacteria | Enterobacterales | Morganellaceae |
| gi\|WP_187129526.1 | Providencia sp. JUb39 | Gammaproteobacteria | Enterobacterales | Morganellaceae |
| gi\|PHM74224.1 | Xenorhabdus kozodoii | Gammaproteobacteria | Enterobacterales | Morganellaceae |
| gi\|WP_068445799.1 | Providencia heimbachae | Gammaproteobacteria | Enterobacterales | Morganellaceae |
| gi\|WP_081989012.1 | Xenorhabdus | Gammaproteobacteria | Enterobacterales | Morganellaceae |
| gi\|WP_086109689.1 | Xenorhabdus vietnamensis | Gammaproteobacteria | Enterobacterales | Morganellaceae |
| gi\|APC12773.1 | Providencia rettgeri | Gammaproteobacteria | Enterobacterales | Morganellaceae |
| gi\|WP_154622000.1 | unclassified Providencia | Gammaproteobacteria | Enterobacterales | Morganellaceae |
| gi\|WP_102139968.1 | Providencia | Gammaproteobacteria | Enterobacterales | Morganellaceae |
| gi\|WP_102780717.1 | Providencia stuartii | Gammaproteobacteria | Enterobacterales | Morganellaceae |
| gi\|WP_039855158.1 | Providencia rustigianii | Gammaproteobacteria | Enterobacterales | Morganellaceae |
| gi\|WP_008910279.1 | Providencia burhodogranariea | Gammaproteobacteria | Enterobacterales | Morganellaceae |
| gi\|WP_081335833.1 | Providencia stuartii | Gammaproteobacteria | Enterobacterales | Morganellaceae |
| gi\|WP_137741639.1 | Pectobacterium polonicum | Gammaproteobacteria | Enterobacterales | Pectobacteriaceae |
| gi\|WP_072009181.1 | Pectobacterium brasiliense | Gammaproteobacteria | Enterobacterales | Pectobacteriaceae |
| gi\|WP_072034320.1 | Pectobacterium fontis | Gammaproteobacteria | Enterobacterales | Pectobacteriaceae |
| gi\|WP_129711737.1 | Pectobacterium zantedeschiae | Gammaproteobacteria | Enterobacterales | Pectobacteriaceae |
| gi\|WP_132454623.1 | Samsonia erythrinae | Gammaproteobacteria | Enterobacterales | Pectobacteriaceae |
| gi\|WP_109053435.1 | Brenneria roseae | Gammaproteobacteria | Enterobacterales | Pectobacteriaceae |
| gi\|WP_136168040.1 | Brenneria sp. CFCC 11842 | Gammaproteobacteria | Enterobacterales | Pectobacteriaceae |
| gi\|WP_172289200.1 | Brenneria sp. hezel4-2-4 | Gammaproteobacteria | Enterobacterales | Pectobacteriaceae |
| gi\|WP_077245877.1 | Dickeya dadantii | Gammaproteobacteria | Enterobacterales | Pectobacteriaceae |
| gi\|WP_071601345.1 | Dickeya chrysanthemi | Gammaproteobacteria | Enterobacterales | Pectobacteriaceae |

| | | | | |
|---|---|---|---|---|
| gi\|WP_103415900.1 | Dickeya dianthicola | Gammaproteobacteria | Enterobacterales | Pectobacteriaceae |
| gi\|WP_123252296.1 | Dickeya undicola | Gammaproteobacteria | Enterobacterales | Pectobacteriaceae |
| gi\|WP_095833837.1 | Brenneria goodwinii | Gammaproteobacteria | Enterobacterales | Pectobacteriaceae |
| gi\|WP_074384638.1 | Dickeya | Gammaproteobacteria | Enterobacterales | Pectobacteriaceae |
| gi\|KAA9002030.1 | Affinibrenneria salicis | Gammaproteobacteria | Enterobacterales | Pectobacteriaceae |
| gi\|WP_125259022.1 | Dickeya lacustris | Gammaproteobacteria | Enterobacterales | Pectobacteriaceae |
| gi\|WP_067486393.1 | Dickeya | Gammaproteobacteria | Enterobacterales | Pectobacteriaceae |
| gi\|QDX29692.1 | Dickeya poaceiphila | Gammaproteobacteria | Enterobacterales | Pectobacteriaceae |
| gi\|WP_168365945.1 | Dickeya zeae | Gammaproteobacteria | Enterobacterales | Pectobacteriaceae |
| gi\|CAG75746.1 | Pectobacterium atrosepticum SCRI1043 | Gammaproteobacteria | Enterobacterales | Pectobacteriaceae |
| gi\|WP_085685997.1 | Lonsdalea | Gammaproteobacteria | Enterobacterales | Pectobacteriaceae |
| gi\|WP_094099739.1 | Lonsdalea iberica | Gammaproteobacteria | Enterobacterales | Pectobacteriaceae |
| gi\|SEA13638.1 | Lonsdalea quercina | Gammaproteobacteria | Enterobacterales | Pectobacteriaceae |
| gi\|ACS85636.1 | Dickeya paradisiaca Ech703 | Gammaproteobacteria | Enterobacterales | Pectobacteriaceae |
| gi\|WP_085650897.1 | Lonsdalea britannica | Gammaproteobacteria | Enterobacterales | Pectobacteriaceae |
| gi\|PKH25373.1 | Enterobacterales bacterium CwR94 | Gammaproteobacteria | Enterobacterales | unclassified Enterobact |
| gi\|GBU12727.1 | Enterobacterales bacterium | Gammaproteobacteria | Enterobacterales | unclassified Enterobact |
| gi\|WP_104921232.1 | Rahnella sp. ERMR1:05 | Gammaproteobacteria | Enterobacterales | Yersiniaceae |
| gi\|WP_112151220.1 | Rahnella | Gammaproteobacteria | Enterobacterales | Yersiniaceae |
| gi\|WP_050763115.1 | Serratia odorifera | Gammaproteobacteria | Enterobacterales | Yersiniaceae |
| gi\|WP_042839628.1 | Yersinia aldovae | Gammaproteobacteria | Enterobacterales | Yersiniaceae |
| gi\|WP_054872860.1 | Yersinia bercovieri | Gammaproteobacteria | Enterobacterales | Yersiniaceae |
| gi\|WP_130380744.1 | Serratia grimesii | Gammaproteobacteria | Enterobacterales | Yersiniaceae |
| gi\|WP_145603983.1 | Yersinia intermedia | Gammaproteobacteria | Enterobacterales | Yersiniaceae |
| gi\|WP_073970232.1 | Serratia ficaria | Gammaproteobacteria | Enterobacterales | Yersiniaceae |
| gi\|WP_050152342.1 | Yersinia frederiksenii | Gammaproteobacteria | Enterobacterales | Yersiniaceae |
| gi\|WP_080987196.1 | Yersinia mollaretii | Gammaproteobacteria | Enterobacterales | Yersiniaceae |
| gi\|WP_056782133.1 | Serratia sp. Leaf51 | Gammaproteobacteria | Enterobacterales | Yersiniaceae |
| gi\|WP_195312397.1 | Serratia marcescens | Gammaproteobacteria | Enterobacterales | Yersiniaceae |
| gi\|WP_017490293.1 | Rouxiella badensis | Gammaproteobacteria | Enterobacterales | Yersiniaceae |
| gi\|WP_159680308.1 | Yersinia canariae | Gammaproteobacteria | Enterobacterales | Yersiniaceae |
| gi\|WP_101826117.1 | Chimaeribacter coloradensis | Gammaproteobacteria | Enterobacterales | Yersiniaceae |
| gi\|AKE09610.1 | Serratia liquefaciens | Gammaproteobacteria | Enterobacterales | Yersiniaceae |
| gi\|QCR36961.1 | Nissabacter sp. SGAir0207 | Gammaproteobacteria | Enterobacterales | Yersiniaceae |
| gi\|SQJ14922.1 | Serratia rubidaea | Gammaproteobacteria | Enterobacterales | Yersiniaceae |
| gi\|WP_140471477.1 | Ewingella americana | Gammaproteobacteria | Enterobacterales | Yersiniaceae |
| gi\|WP_025376622.1 | Yersinia enterocolitica | Gammaproteobacteria | Enterobacterales | Yersiniaceae |
| gi\|PLR44733.1 | Chimaeribacter arupi | Gammaproteobacteria | Enterobacterales | Yersiniaceae |
| gi\|CFQ40140.1 | Yersinia aleksiciae | Gammaproteobacteria | Enterobacterales | Yersiniaceae |
| gi\|WP_037431391.1 | Serratia plymuthica | Gammaproteobacteria | Enterobacterales | Yersiniaceae |
| gi\|SNY83627.1 | Serratia sp. JKS000199 | Gammaproteobacteria | Enterobacterales | Yersiniaceae |
| gi\|SUI49246.1 | Serratia marcescens | Gammaproteobacteria | Enterobacterales | Yersiniaceae |
| gi\|WP_147882062.1 | Serratia marcescens | Gammaproteobacteria | Enterobacterales | Yersiniaceae |

| | | | | |
|---|---|---|---|---|
| gi\|ANI30454.1 | Yersinia entomophaga | Gammaproteobacteria | Enterobacterales | Yersiniaceae |
| gi\|WP_082026996.1 | Serratia symbiotica | Gammaproteobacteria | Enterobacterales | Yersiniaceae |
| gi\|AHK19002.1 | Yersinia similis | Gammaproteobacteria | Enterobacterales | Yersiniaceae |
| gi\|WP_009636415.1 | Serratia sp. M24T3 | Gammaproteobacteria | Enterobacterales | Yersiniaceae |
| gi\|EEP99988.1 | Yersinia ruckeri ATCC 29473 | Gammaproteobacteria | Enterobacterales | Yersiniaceae |
| gi\|WP_050879265.1 | Yersinia frederiksenii | Gammaproteobacteria | Enterobacterales | Yersiniaceae |
| gi\|WP_152554915.1 | Serratia | Gammaproteobacteria | Enterobacterales | Yersiniaceae |
| gi\|WP_195314833.1 | Serratia marcescens | Gammaproteobacteria | Enterobacterales | Yersiniaceae |
| gi\|WP_061795296.1 | Serratia | Gammaproteobacteria | Enterobacterales | Yersiniaceae |
| gi\|WP_084983130.1 | Rouxiella silvae | Gammaproteobacteria | Enterobacterales | Yersiniaceae |
| gi\|WP_169401131.1 | Rouxiella aceris | Gammaproteobacteria | Enterobacterales | Yersiniaceae |
| gi\|WP_122016596.1 | Serratia marcescens | Gammaproteobacteria | Enterobacterales | Yersiniaceae |
| gi\|WP_071988463.1 | Serratia sp. M24T3 | Gammaproteobacteria | Enterobacterales | Yersiniaceae |
| gi\|WP_021016087.1 | Serratia sp. ATCC 39006 | Gammaproteobacteria | Enterobacterales | Yersiniaceae |
| gi\|GAK27632.1 | Serratia liquefaciens FK01 | Gammaproteobacteria | Enterobacterales | Yersiniaceae |
| gi\|WP_045047413.1 | Rouxiella chamberiensis | Gammaproteobacteria | Enterobacterales | Yersiniaceae |
| gi\|WP_116727192.1 | Serratia sp. S1B | Gammaproteobacteria | Enterobacterales | Yersiniaceae |
| gi\|WP_048914709.1 | Erwiniaceae | Gammaproteobacteria | Enterobacterales | |
| gi\|WP_157725398.1 | Erwiniaceae | Gammaproteobacteria | Enterobacterales | |
| gi\|WP_094168992.1 | Enterobacteriaceae | Gammaproteobacteria | Enterobacterales | |
| gi\|TXH05318.1 | Sinobacteraceae bacterium | Gammaproteobacteria | Nevskiales | Sinobacteraceae |
| gi\|WP_107939584.1 | Stenotrophobium rhamnosiphilum | Gammaproteobacteria | Nevskiales | Sinobacteraceae |
| gi\|WP_143383683.1 | Fontimonas thermophila | Gammaproteobacteria | Nevskiales | Sinobacteraceae |
| gi\|WP_090134503.1 | Kushneria avicenniae | Gammaproteobacteria | Oceanospirillales | Halomonadaceae |
| gi\|WP_108841727.1 | Kushneria phyllosphaerae | Gammaproteobacteria | Oceanospirillales | Halomonadaceae |
| gi\|WP_189517369.1 | Kushneria pakistanensis | Gammaproteobacteria | Oceanospirillales | Halomonadaceae |
| gi\|WP_019952807.1 | Kushneria aurantia | Gammaproteobacteria | Oceanospirillales | Halomonadaceae |
| gi\|WP_175070446.1 | Halomonas taeanensis | Gammaproteobacteria | Oceanospirillales | Halomonadaceae |
| gi\|TZG41488.1 | Halomonas eurihalina | Gammaproteobacteria | Oceanospirillales | Halomonadaceae |
| gi\|WP_110641564.1 | Salinicola sp. CPA57 | Gammaproteobacteria | Oceanospirillales | Halomonadaceae |
| gi\|WP_183388268.1 | Halomonas organivorans | Gammaproteobacteria | Oceanospirillales | Halomonadaceae |
| gi\|WP_168709184.1 | Halomonas borealis | Gammaproteobacteria | Oceanospirillales | Halomonadaceae |
| gi\|WP_075563784.1 | Salinicola | Gammaproteobacteria | Oceanospirillales | Halomonadaceae |
| gi\|WP_110649792.1 | Salinicola peritrichatus | Gammaproteobacteria | Oceanospirillales | Halomonadaceae |
| gi\|WP_089730860.1 | Halomonas muralis | Gammaproteobacteria | Oceanospirillales | Halomonadaceae |
| gi\|WP_146742484.1 | Halomonas taeanensis | Gammaproteobacteria | Oceanospirillales | Halomonadaceae |
| gi\|WP_087720220.1 | Salinicola salarius | Gammaproteobacteria | Oceanospirillales | Halomonadaceae |
| gi\|GEK72601.1 | Halomonas halophila | Gammaproteobacteria | Oceanospirillales | Halomonadaceae |
| gi\|WP_110708835.1 | Salinicola sp. CR57 | Gammaproteobacteria | Oceanospirillales | Halomonadaceae |
| gi\|WP_168709025.1 | Halomonas niordiana | Gammaproteobacteria | Oceanospirillales | Halomonadaceae |
| gi\|WP_064122350.1 | Halotalea alkalilenta | Gammaproteobacteria | Oceanospirillales | Halomonadaceae |
| gi\|WP_110656182.1 | Salinicola halimionae | Gammaproteobacteria | Oceanospirillales | Halomonadaceae |
| gi\|WP_168380891.1 | Halomonas sp. EAR18 | Gammaproteobacteria | Oceanospirillales | Halomonadaceae |

| | | | | |
|---|---|---|---|---|
| gi\|WP_083861799.1 | Halomonas sp. KM-1 | Gammaproteobacteria | Oceanospirillales | Halomonadaceae |
| gi\|KAA0012654.1 | Halomonas sp. L5 | Gammaproteobacteria | Oceanospirillales | Halomonadaceae |
| gi\|WP_129140888.1 | Halomonas coralii | Gammaproteobacteria | Oceanospirillales | Halomonadaceae |
| gi\|WP_086621657.1 | Kushneria konosiri | Gammaproteobacteria | Oceanospirillales | Halomonadaceae |
| gi\|WP_108448011.1 | Halomonas sp. BN3-1 | Gammaproteobacteria | Oceanospirillales | Halomonadaceae |
| gi\|WP_083933030.1 | Halomonas lutea | Gammaproteobacteria | Oceanospirillales | Halomonadaceae |
| gi\|WP_083970158.1 | Halomonas sp. S2151 | Gammaproteobacteria | Oceanospirillales | Halomonadaceae |
| gi\|WP_021818295.1 | Halomonas huangheensis | Gammaproteobacteria | Oceanospirillales | Halomonadaceae |
| gi\|WP_035592897.1 | Halomonas | Gammaproteobacteria | Oceanospirillales | Halomonadaceae |
| gi\|WP_073435930.1 | Halomonas cupida | Gammaproteobacteria | Oceanospirillales | Halomonadaceae |
| gi\|WP_016418449.1 | Halomonas anticariensis | Gammaproteobacteria | Oceanospirillales | Halomonadaceae |
| gi\|WP_157959069.1 | Salinicola endophyticus | Gammaproteobacteria | Oceanospirillales | Halomonadaceae |
| gi\|WP_110685137.1 | Salinicola aestuarinus | Gammaproteobacteria | Oceanospirillales | Halomonadaceae |
| gi\|WP_157958859.1 | Salinicola | Gammaproteobacteria | Oceanospirillales | Halomonadaceae |
| gi\|WP_177223467.1 | Halomonas xianhensis | Gammaproteobacteria | Oceanospirillales | Halomonadaceae |
| gi\|WP_137079981.1 | Halomonas caseinilytica | Gammaproteobacteria | Oceanospirillales | Halomonadaceae |
| gi\|WP_104202188.1 | Halomonas saliphila | Gammaproteobacteria | Oceanospirillales | Halomonadaceae |
| gi\|WP_149286206.1 | Halomonas sp. Y2R2 | Gammaproteobacteria | Oceanospirillales | Halomonadaceae |
| gi\|WP_189442955.1 | Salinicola rhizosphaerae | Gammaproteobacteria | Oceanospirillales | Halomonadaceae |
| gi\|WP_090134656.1 | Kushneria avicenniae | Gammaproteobacteria | Oceanospirillales | Halomonadaceae |
| gi\|WP_192527161.1 | Halomonas sp. FME16 | Gammaproteobacteria | Oceanospirillales | Halomonadaceae |
| gi\|WP_165942923.1 | Marinomonas sp. JHZ-47 | Gammaproteobacteria | Oceanospirillales | Oceanospirillaceae |
| gi\|CUB05060.1 | Marinomonas fungiae | Gammaproteobacteria | Oceanospirillales | Oceanospirillaceae |
| gi\|WP_132291438.1 | Marinobacterium mangrovicola | Gammaproteobacteria | Oceanospirillales | Oceanospirillaceae |
| gi\|WP_114413535.1 | Marinomonas foliarum | Gammaproteobacteria | Oceanospirillales | Oceanospirillaceae |
| gi\|WP_084545992.1 | Marinomonas profundimaris | Gammaproteobacteria | Oceanospirillales | Oceanospirillaceae |
| gi\|WP_111639010.1 | Marinomonas shanghaiensis | Gammaproteobacteria | Oceanospirillales | Oceanospirillaceae |
| gi\|WP_168822152.1 | Marinomonas sp. M1K-6 | Gammaproteobacteria | Oceanospirillales | Oceanospirillaceae |
| gi\|WP_067095488.1 | Marinomonas atlantica | Gammaproteobacteria | Oceanospirillales | Oceanospirillaceae |
| gi\|WP_083766327.1 | Marinomonas sp. MWYL1 | Gammaproteobacteria | Oceanospirillales | Oceanospirillaceae |
| gi\|WP_133011349.1 | Marinomonas sp. JHZ-47 | Gammaproteobacteria | Oceanospirillales | Oceanospirillaceae |
| gi\|WP_012069623.1 | Marinomonas sp. MWYL1 | Gammaproteobacteria | Oceanospirillales | Oceanospirillaceae |
| gi\|WP_133003778.1 | Marinomonas sp. KMM3893 | Gammaproteobacteria | Oceanospirillales | Oceanospirillaceae |
| gi\|AEF55044.1 | Marinomonas posidonica IVIA-Po-181 | Gammaproteobacteria | Oceanospirillales | Oceanospirillaceae |
| gi\|WP_113917613.1 | Marinomonas rhizomae | Gammaproteobacteria | Oceanospirillales | Oceanospirillaceae |
| gi\|WP_176335522.1 | Marinomonas primoryensis | Gammaproteobacteria | Oceanospirillales | Oceanospirillaceae |
| gi\|WP_111606065.1 | Marinomonas arctica | Gammaproteobacteria | Oceanospirillales | Oceanospirillaceae |
| gi\|RNF49667.1 | Marinomonas hwangdonensis | Gammaproteobacteria | Oceanospirillales | Oceanospirillaceae |
| gi\|WP_191595040.1 | Marinomonas colpomeniae | Gammaproteobacteria | Oceanospirillales | Oceanospirillaceae |
| gi\|WP_063333334.1 | Marinomonas sp. TW1 | Gammaproteobacteria | Oceanospirillales | Oceanospirillaceae |
| gi\|WP_067016555.1 | Marinomonas spartinae | Gammaproteobacteria | Oceanospirillales | Oceanospirillaceae |
| gi\|WP_139116540.1 | Terasakiispira papahanaumokuakeensis | Gammaproteobacteria | Oceanospirillales | Oceanospirillales incert sedis |
| gi\|WP_065618068.1 | Gilliamella apicola | Gammaproteobacteria | Orbales | Orbaceae |

| | | | | |
|---|---|---|---|---|
| gi\|CAG68591.1 | Acinetobacter baylyi ADP1 | Gammaproteobacteria | Pseudomonadales | Moraxellaceae |
| gi\|WP_191012944.1 | Acinetobacter seifertii | Gammaproteobacteria | Pseudomonadales | Moraxellaceae |
| gi\|WP_111885208.1 | Acinetobacter sp. CFCC 11171 | Gammaproteobacteria | Pseudomonadales | Moraxellaceae |
| gi\|WP_109441437.1 | Acinetobacter haemolyticus | Gammaproteobacteria | Pseudomonadales | Moraxellaceae |
| gi\|WP_044102552.1 | Acinetobacter pittii | Gammaproteobacteria | Pseudomonadales | Moraxellaceae |
| gi\|WP_055415851.1 | Acinetobacter soli | Gammaproteobacteria | Pseudomonadales | Moraxellaceae |
| gi\|WP_034595322.1 | Acinetobacter sp. CIP-A165 | Gammaproteobacteria | Pseudomonadales | Moraxellaceae |
| gi\|WP_131322484.1 | Acinetobacter sp. ANC 4178 | Gammaproteobacteria | Pseudomonadales | Moraxellaceae |
| gi\|OJU75629.1 | Acinetobacter sp. 39-4 | Gammaproteobacteria | Pseudomonadales | Moraxellaceae |
| gi\|WP_174560311.1 | Acinetobacter bouvetii | Gammaproteobacteria | Pseudomonadales | Moraxellaceae |
| gi\|WP_142770096.1 | Acinetobacter tandoii | Gammaproteobacteria | Pseudomonadales | Moraxellaceae |
| gi\|WP_153373038.1 | Acinetobacter wanghuae | Gammaproteobacteria | Pseudomonadales | Moraxellaceae |
| gi\|GGA29965.1 | Acinetobacter modestus | Gammaproteobacteria | Pseudomonadales | Moraxellaceae |
| gi\|WP_159138879.1 | Acinetobacter lwoffii | Gammaproteobacteria | Pseudomonadales | Moraxellaceae |
| gi\|WP_004944206.1 | Acinetobacter soli | Gammaproteobacteria | Pseudomonadales | Moraxellaceae |
| gi\|WP_068885789.1 | Acinetobacter celticus | Gammaproteobacteria | Pseudomonadales | Moraxellaceae |
| gi\|WP_067731744.1 | Acinetobacter sp. NCu2D-2 | Gammaproteobacteria | Pseudomonadales | Moraxellaceae |
| gi\|WP_151708788.1 | Acinetobacter brisouii | Gammaproteobacteria | Pseudomonadales | Moraxellaceae |
| gi\|WP_086213987.1 | Acinetobacter sp. ANC 3813 | Gammaproteobacteria | Pseudomonadales | Moraxellaceae |
| gi\|WP_086200303.1 | Acinetobacter sp. ANC 4169 | Gammaproteobacteria | Pseudomonadales | Moraxellaceae |
| gi\|WP_086164838.1 | Acinetobacter sp. ANC 4654 | Gammaproteobacteria | Pseudomonadales | Moraxellaceae |
| gi\|WP_120375757.1 | Acinetobacter | Gammaproteobacteria | Pseudomonadales | Moraxellaceae |
| gi\|EEY85646.1 | Acinetobacter radioresistens SH164 | Gammaproteobacteria | Pseudomonadales | Moraxellaceae |
| gi\|WP_099338022.1 | Acinetobacter sp. LoGeW2-3 | Gammaproteobacteria | Pseudomonadales | Moraxellaceae |
| gi\|WP_196076406.1 | Acinetobacter | Gammaproteobacteria | Pseudomonadales | Moraxellaceae |
| gi\|ESK45186.1 | Acinetobacter oleivorans CIP 110421 | Gammaproteobacteria | Pseudomonadales | Moraxellaceae |
| gi\|ENV70964.1 | Acinetobacter towneri DSM 14962 = CIP 107472 | Gammaproteobacteria | Pseudomonadales | Moraxellaceae |
| gi\|WP_042128043.1 | Pseudomonas japonica | Gammaproteobacteria | Pseudomonadales | Pseudomonadaceae |
| gi\|WP_055101167.1 | Pseudomonas endophytica | Gammaproteobacteria | Pseudomonadales | Pseudomonadaceae |
| gi\|WP_177105415.1 | Pseudomonas gingeri | Gammaproteobacteria | Pseudomonadales | Pseudomonadaceae |
| gi\|WP_075804705.1 | Pseudomonas putida | Gammaproteobacteria | Pseudomonadales | Pseudomonadaceae |
| gi\|PXX76248.1 | Pseudomonas sp. LAMO17WK12:I9 | Gammaproteobacteria | Pseudomonadales | Pseudomonadaceae |
| gi\|WP_110971387.1 | Pseudomonas huaxiensis | Gammaproteobacteria | Pseudomonadales | Pseudomonadaceae |
| gi\|WP_011104228.1 | Pseudomonas syringae group genomosp. 3 | Gammaproteobacteria | Pseudomonadales | Pseudomonadaceae |
| gi\|WP_191485853.1 | Pseudomonas sp. FEN | Gammaproteobacteria | Pseudomonadales | Pseudomonadaceae |
| gi\|WP_161719781.1 | Pseudomonas sp. Fl4BN2 | Gammaproteobacteria | Pseudomonadales | Pseudomonadaceae |
| gi\|WP_046810047.1 | Pseudomonas psychrophila | Gammaproteobacteria | Pseudomonadales | Pseudomonadaceae |
| gi\|WP_102881962.1 | Pseudomonas protegens | Gammaproteobacteria | Pseudomonadales | Pseudomonadaceae |
| gi\|WP_060481707.1 | Pseudomonas sp. NBRC 111119 | Gammaproteobacteria | Pseudomonadales | Pseudomonadaceae |
| gi\|WP_146426531.1 | Pseudomonas saxonica | Gammaproteobacteria | Pseudomonadales | Pseudomonadaceae |
| gi\|WP_180274397.1 | Pseudomonas viridiflava | Gammaproteobacteria | Pseudomonadales | Pseudomonadaceae |
| gi\|WP_048382691.1 | Pseudomonas | Gammaproteobacteria | Pseudomonadales | Pseudomonadaceae |
| gi\|WP_038615109.1 | Pseudomonas alkylphenolica | Gammaproteobacteria | Pseudomonadales | Pseudomonadaceae |

| | | | | |
|---|---|---|---|---|
| gi\|WP_158461086.1 | Pseudomonas fluorescens | Gammaproteobacteria | Pseudomonadales | Pseudomonadaceae |
| gi\|WP_050978674.1 | Pseudomonas fuscovaginae | Gammaproteobacteria | Pseudomonadales | Pseudomonadaceae |
| gi\|WP_028695549.1 | Pseudomonas cremoricolorata | Gammaproteobacteria | Pseudomonadales | Pseudomonadaceae |
| gi\|WP_194286255.1 | Pseudomonas helleri | Gammaproteobacteria | Pseudomonadales | Pseudomonadaceae |
| gi\|WP_094990109.1 | Pseudomonas lundensis | Gammaproteobacteria | Pseudomonadales | Pseudomonadaceae |
| gi\|WP_136916798.1 | Pseudomonas putida | Gammaproteobacteria | Pseudomonadales | Pseudomonadaceae |
| gi\|WP_120266342.1 | Pseudomonas sp. TMW 2.1634 | Gammaproteobacteria | Pseudomonadales | Pseudomonadaceae |
| gi\|WP_119146212.1 | Pseudomonas reidholzensis | Gammaproteobacteria | Pseudomonadales | Pseudomonadaceae |
| gi\|WP_169909242.1 | Pseudomonas proteolytica | Gammaproteobacteria | Pseudomonadales | Pseudomonadaceae |
| gi\|WP_159412294.1 | Pseudomonas putida | Gammaproteobacteria | Pseudomonadales | Pseudomonadaceae |
| gi\|WP_177073590.1 | Pseudomonas gingeri | Gammaproteobacteria | Pseudomonadales | Pseudomonadaceae |
| gi\|WP_087499542.1 | Pseudomonas sp. SID14000 | Gammaproteobacteria | Pseudomonadales | Pseudomonadaceae |
| gi\|WP_196130142.1 | Pseudomonas fulva | Gammaproteobacteria | Pseudomonadales | Pseudomonadaceae |
| gi\|WP_084858778.1 | Pseudomonas putida | Gammaproteobacteria | Pseudomonadales | Pseudomonadaceae |
| gi\|GFM82950.1 | Pseudomonas cichorii | Gammaproteobacteria | Pseudomonadales | Pseudomonadaceae |
| gi\|WP_177083287.1 | Pseudomonas | Gammaproteobacteria | Pseudomonadales | Pseudomonadaceae |
| gi\|PCE22918.1 | Pseudomonas acidophila | Gammaproteobacteria | Pseudomonadales | Pseudomonadaceae |
| gi\|WP_169916621.1 | Pseudomonas sp. WS 5051 | Gammaproteobacteria | Pseudomonadales | Pseudomonadaceae |
| gi\|WP_054062172.1 | Pseudomonas fuscovaginae | Gammaproteobacteria | Pseudomonadales | Pseudomonadaceae |
| gi\|WP_080520264.1 | Pseudomonas tolaasii | Gammaproteobacteria | Pseudomonadales | Pseudomonadaceae |
| gi\|WP_084920884.1 | Pseudomonas | Gammaproteobacteria | Pseudomonadales | Pseudomonadaceae |
| gi\|WP_029613675.1 | Pseudomonas parafulva | Gammaproteobacteria | Pseudomonadales | Pseudomonadaceae |
| gi\|WP_100632893.1 | Pseudomonas | Gammaproteobacteria | Pseudomonadales | Pseudomonadaceae |
| gi\|WP_116550677.1 | Pseudomonas sp. SDI | Gammaproteobacteria | Pseudomonadales | Pseudomonadaceae |
| gi\|WP_087881671.1 | Pseudomonas floridensis | Gammaproteobacteria | Pseudomonadales | Pseudomonadaceae |
| gi\|POF90934.1 | Pseudomonas putida | Gammaproteobacteria | Pseudomonadales | Pseudomonadaceae |
| gi\|WP_163934914.1 | Pseudomonas laurentiana | Gammaproteobacteria | Pseudomonadales | Pseudomonadaceae |
| gi\|RML58637.1 | Pseudomonas amygdali pv. morsprunorum | Gammaproteobacteria | Pseudomonadales | Pseudomonadaceae |
| gi\|WP_196162599.1 | Pseudomonas guariconensis | Gammaproteobacteria | Pseudomonadales | Pseudomonadaceae |
| gi\|KAF2408362.1 | Pseudomonas antarctica | Gammaproteobacteria | Pseudomonadales | Pseudomonadaceae |
| gi\|WP_176569752.1 | Pseudomonas eucalypticola | Gammaproteobacteria | Pseudomonadales | Pseudomonadaceae |
| gi\|WP_108239496.1 | unclassified Pseudomonas | Gammaproteobacteria | Pseudomonadales | Pseudomonadaceae |
| gi\|WP_123329685.1 | Pseudomonas chlororaphis | Gammaproteobacteria | Pseudomonadales | Pseudomonadaceae |
| gi\|RZI75899.1 | Pseudomonas sp. | Gammaproteobacteria | Pseudomonadales | Pseudomonadaceae |
| gi\|WP_186605891.1 | Pseudomonas lurida | Gammaproteobacteria | Pseudomonadales | Pseudomonadaceae |
| gi\|WP_123402365.1 | Pseudomonas frederiksbergensis | Gammaproteobacteria | Pseudomonadales | Pseudomonadaceae |
| gi\|WP_084709945.1 | Pseudomonas sp. StFLB209 | Gammaproteobacteria | Pseudomonadales | Pseudomonadaceae |
| gi\|WP_053931579.1 | Pseudomonas coronafaciens | Gammaproteobacteria | Pseudomonadales | Pseudomonadaceae |
| gi\|WP_028620206.1 | Pseudomonas sp. Ant30-3 | Gammaproteobacteria | Pseudomonadales | Pseudomonadaceae |
| gi\|AGL84442.1 | Pseudomonas protegens CHA0 | Gammaproteobacteria | Pseudomonadales | Pseudomonadaceae |
| gi\|KPY66692.1 | Pseudomonas syringae pv. spinaceae | Gammaproteobacteria | Pseudomonadales | Pseudomonadaceae |
| gi\|WP_080482209.1 | Pseudomonas syringae | Gammaproteobacteria | Pseudomonadales | Pseudomonadaceae |
| gi\|WP_003457951.1 | Pseudomonas furukawaii | Gammaproteobacteria | Pseudomonadales | Pseudomonadaceae |

| | | | | |
|---|---|---|---|---|
| gi\|WP_043230071.1 | Pseudomonas sp. CF161 | Gammaproteobacteria | Pseudomonadales | Pseudomonadaceae |
| gi\|WP_008368675.1 | Pseudomonas sp. M47T1 | Gammaproteobacteria | Pseudomonadales | Pseudomonadaceae |
| gi\|KTC40440.1 | Pseudomonas sp. ABAC61 | Gammaproteobacteria | Pseudomonadales | Pseudomonadaceae |
| gi\|WP_169895290.1 | Pseudomonas poae | Gammaproteobacteria | Pseudomonadales | Pseudomonadaceae |
| gi\|WP_166357747.1 | Pseudomonas sp. PS24 | Gammaproteobacteria | Pseudomonadales | Pseudomonadaceae |
| gi\|SFL53168.1 | Rugamonas rubra | Gammaproteobacteria | Pseudomonadales | Pseudomonadaceae |
| gi\|WP_110951880.1 | Pseudomonas bohemica | Gammaproteobacteria | Pseudomonadales | Pseudomonadaceae |
| gi\|WP_050507954.1 | Pseudomonas syringae | Gammaproteobacteria | Pseudomonadales | Pseudomonadaceae |
| gi\|PCE22851.1 | Pseudomonas acidophila | Gammaproteobacteria | Pseudomonadales | Pseudomonadaceae |
| gi\|WP_081563925.1 | Pseudomonas sp. Bc-h | Gammaproteobacteria | Pseudomonadales | Pseudomonadaceae |
| gi\|WP_145136830.1 | Pseudomonas duriflava | Gammaproteobacteria | Pseudomonadales | Pseudomonadaceae |
| gi\|WP_196176084.1 | Pseudomonas fulva | Gammaproteobacteria | Pseudomonadales | Pseudomonadaceae |
| gi\|GFM68635.1 | Pseudomonas cichorii | Gammaproteobacteria | Pseudomonadales | Pseudomonadaceae |
| gi\|WP_007250950.1 | Pseudomonas syringae group | Gammaproteobacteria | Pseudomonadales | Pseudomonadaceae |
| gi\|WP_181095447.1 | Pseudomonas entomophila | Gammaproteobacteria | Pseudomonadales | Pseudomonadaceae |
| gi\|WP_133217254.1 | Pseudomonas sp. H9 | Gammaproteobacteria | Pseudomonadales | Pseudomonadaceae |
| gi\|WP_064675682.1 | Pseudomonas | Gammaproteobacteria | Pseudomonadales | Pseudomonadaceae |
| gi\|WP_181114497.1 | Pseudomonas viridiflava | Gammaproteobacteria | Pseudomonadales | Pseudomonadaceae |
| gi\|WP_028626980.1 | Pseudomonas | Gammaproteobacteria | Pseudomonadales | Pseudomonadaceae |
| gi\|WP_016495470.1 | Pseudomonas resinovorans | Gammaproteobacteria | Pseudomonadales | Pseudomonadaceae |
| gi\|WP_175387661.1 | Pseudomonas sp. C2B4 | Gammaproteobacteria | Pseudomonadales | Pseudomonadaceae |
| gi\|WP_122725380.1 | Pseudomonas viridiflava | Gammaproteobacteria | Pseudomonadales | Pseudomonadaceae |
| gi\|WP_123335153.1 | Pseudomonas chlororaphis | Gammaproteobacteria | Pseudomonadales | Pseudomonadaceae |
| gi\|WP_181130789.1 | Pseudomonas capeferrum | Gammaproteobacteria | Pseudomonadales | Pseudomonadaceae |
| gi\|TDV85480.1 | Pseudomonas mandelii | Gammaproteobacteria | Pseudomonadales | Pseudomonadaceae |
| gi\|WP_095109428.1 | Pseudomonas sp. Irchel 3E20 | Gammaproteobacteria | Pseudomonadales | Pseudomonadaceae |
| gi\|SEJ53350.1 | Pseudomonas sp. NFR16 | Gammaproteobacteria | Pseudomonadales | Pseudomonadaceae |
| gi\|WP_084229659.1 | unclassified Pseudomonas | Gammaproteobacteria | Pseudomonadales | Pseudomonadaceae |
| gi\|WP_181121538.1 | Pseudomonas japonica | Gammaproteobacteria | Pseudomonadales | Pseudomonadaceae |
| gi\|WP_177145547.1 | Pseudomonas gingeri | Gammaproteobacteria | Pseudomonadales | Pseudomonadaceae |
| gi\|WP_092408307.1 | Pseudomonas sp. NFACC02 | Gammaproteobacteria | Pseudomonadales | Pseudomonadaceae |
| gi\|WP_110618105.1 | Pseudomonas sp. OV467 | Gammaproteobacteria | Pseudomonadales | Pseudomonadaceae |
| gi\|WP_110951092.1 | Pseudomonas bohemica | Gammaproteobacteria | Pseudomonadales | Pseudomonadaceae |
| gi\|WP_150806883.1 | Pseudomonas fluorescens | Gammaproteobacteria | Pseudomonadales | Pseudomonadaceae |
| gi\|WP_102668591.1 | Pseudomonas sp. GW456-11-11-14-LB1 | Gammaproteobacteria | Pseudomonadales | Pseudomonadaceae |
| gi\|SDH61201.1 | Pseudomonas panipatensis | Gammaproteobacteria | Pseudomonadales | Pseudomonadaceae |
| gi\|WP_045060349.1 | Pseudomonas sp. ES3-33 | Gammaproteobacteria | Pseudomonadales | Pseudomonadaceae |
| gi\|WP_162863730.1 | Pseudomonas viridiflava | Gammaproteobacteria | Pseudomonadales | Pseudomonadaceae |
| gi\|WP_160108996.1 | Pseudomonas sp. IzPS43_3003 | Gammaproteobacteria | Pseudomonadales | Pseudomonadaceae |
| gi\|WP_083213929.1 | Pseudomonas sp. 35 E 8 | Gammaproteobacteria | Pseudomonadales | Pseudomonadaceae |
| gi\|WP_175653993.1 | Pseudomonas sp. Marseille-P9899 | Gammaproteobacteria | Pseudomonadales | Pseudomonadaceae |
| gi\|WP_177108902.1 | Pseudomonas gingeri | Gammaproteobacteria | Pseudomonadales | Pseudomonadaceae |
| gi\|CDF82426.1 | Pseudomonas knackmussii B13 | Gammaproteobacteria | Pseudomonadales | Pseudomonadaceae |

| | | | | |
|---|---|---|---|---|
| gi\|WP_192055715.1 | Pseudomonas sp. CFBP 8758 | Gammaproteobacteria | Pseudomonadales | Pseudomonadaceae |
| gi\|WP_038615105.1 | Pseudomonas alkylphenolica | Gammaproteobacteria | Pseudomonadales | Pseudomonadaceae |
| gi\|WP_103103107.1 | Pseudomonas sp. LFM046 | Gammaproteobacteria | Pseudomonadales | Pseudomonadaceae |
| gi\|WP_084315086.1 | Pseudomonas jinjuensis | Gammaproteobacteria | Pseudomonadales | Pseudomonadaceae |
| gi\|SCX64922.1 | Pseudomonas sp. NFACC32-1 | Gammaproteobacteria | Pseudomonadales | Pseudomonadaceae |
| gi\|WP_164708592.1 | Pseudomonas viridiflava | Gammaproteobacteria | Pseudomonadales | Pseudomonadaceae |
| gi\|WP_110966134.1 | Pseudomonas putida | Gammaproteobacteria | Pseudomonadales | Pseudomonadaceae |
| gi\|SDX89861.1 | Pseudomonas kuykendallii | Gammaproteobacteria | Pseudomonadales | Pseudomonadaceae |
| gi\|WP_052028504.1 | Pseudomonas syringae | Gammaproteobacteria | Pseudomonadales | Pseudomonadaceae |
| gi\|WP_042946418.1 | Pseudomonas extremaustralis | Gammaproteobacteria | Pseudomonadales | Pseudomonadaceae |
| gi\|WP_092312594.1 | Pseudomonas saponiphila | Gammaproteobacteria | Pseudomonadales | Pseudomonadaceae |
| gi\|WP_038411895.1 | Pseudomonas cremoricolorata | Gammaproteobacteria | Pseudomonadales | Pseudomonadaceae |
| gi\|WP_192307775.1 | Pseudomonas sp. PDM04 | Gammaproteobacteria | Pseudomonadales | Pseudomonadaceae |
| gi\|AHC36358.1 | Pseudomonas sp. TKP | Gammaproteobacteria | Pseudomonadales | Pseudomonadaceae |
| gi\|WP_020481086.1 | Pseudomonas fuscovaginae | Gammaproteobacteria | Pseudomonadales | Pseudomonadaceae |
| gi\|WP_189395442.1 | Pseudomonas laurentiana | Gammaproteobacteria | Pseudomonadales | Pseudomonadaceae |
| gi\|WP_036998167.1 | Pseudomonas | Gammaproteobacteria | Pseudomonadales | Pseudomonadaceae |
| gi\|WP_068828533.1 | Pseudomonas sp. BMS12 | Gammaproteobacteria | Pseudomonadales | Pseudomonadaceae |
| gi\|WP_009394303.1 | Pseudomonas putida | Gammaproteobacteria | Pseudomonadales | Pseudomonadaceae |
| gi\|WP_133750860.1 | Pseudomonas sp. LP_7_YM | Gammaproteobacteria | Pseudomonadales | Pseudomonadaceae |
| gi\|WP_105753980.1 | Pseudomonas | Gammaproteobacteria | Pseudomonadales | Pseudomonadaceae |
| gi\|WP_121136385.1 | Pseudomonas asplenii | Gammaproteobacteria | Pseudomonadales | Pseudomonadaceae |
| gi\|VVO01663.1 | Pseudomonas fluorescens | Gammaproteobacteria | Pseudomonadales | Pseudomonadaceae |
| gi\|OWJ91598.1 | Pseudomonas sp. A46 | Gammaproteobacteria | Pseudomonadales | Pseudomonadaceae |
| gi\|WP_119955947.1 | Pseudomonas sp. K1S02-6 | Gammaproteobacteria | Pseudomonadales | Pseudomonadaceae |
| gi\|WP_166595955.1 | Pseudomonas sp. SLFW | Gammaproteobacteria | Pseudomonadales | Pseudomonadaceae |
| gi\|WP_169916640.1 | Pseudomonas sp. WS 5051 | Gammaproteobacteria | Pseudomonadales | Pseudomonadaceae |
| gi\|WP_119953742.1 | Pseudomonas sp. K1S02-6 | Gammaproteobacteria | Pseudomonadales | Pseudomonadaceae |
| gi\|WP_003439179.1 | Pseudomonas | Gammaproteobacteria | Pseudomonadales | Pseudomonadaceae |
| gi\|WP_166357745.1 | Pseudomonas sp. PS24 | Gammaproteobacteria | Pseudomonadales | Pseudomonadaceae |
| gi\|KAF1032015.1 | Pseudomonas sp. | Gammaproteobacteria | Pseudomonadales | Pseudomonadaceae |
| gi\|WP_168082815.1 | Pseudomonas | Gammaproteobacteria | Pseudomonadales | Pseudomonadaceae |
| gi\|SEI20604.1 | Pseudomonas fuscovaginae | Gammaproteobacteria | Pseudomonadales | Pseudomonadaceae |
| gi\|WP_166568698.1 | Pseudomonas sp. R5(2019) | Gammaproteobacteria | Pseudomonadales | Pseudomonadaceae |
| gi\|KRW67666.1 | Pseudomonas sp. TTU2014-096BSC | Gammaproteobacteria | Pseudomonadales | Pseudomonadaceae |
| gi\|WP_193682375.1 | Pseudomonas lopnurensis | Gammaproteobacteria | Pseudomonadales | Pseudomonadaceae |
| gi\|SED80824.1 | Pseudomonas coleopterorum | Gammaproteobacteria | Pseudomonadales | Pseudomonadaceae |
| gi\|WP_110595748.1 | Pseudomonas | Gammaproteobacteria | Pseudomonadales | Pseudomonadaceae |
| gi\|WP_123329689.1 | Pseudomonas chlororaphis | Gammaproteobacteria | Pseudomonadales | Pseudomonadaceae |
| gi\|WP_073262080.1 | Pseudomonas punonensis | Gammaproteobacteria | Pseudomonadales | Pseudomonadaceae |
| gi\|SDU29038.1 | Pseudomonas guangdongensis | Gammaproteobacteria | Pseudomonadales | Pseudomonadaceae |
| gi\|WP_019341043.1 | Pseudomonas stutzeri | Gammaproteobacteria | Pseudomonadales | Pseudomonadaceae |
| gi\|WP_138407686.1 | Pseudomonas nosocomialis | Gammaproteobacteria | Pseudomonadales | Pseudomonadaceae |

| | | | | |
|---|---|---|---|---|
| gi\|WP_137822400.1 | Pseudomonas sp. D(2018) | Gammaproteobacteria | Pseudomonadales | Pseudomonadaceae |
| gi\|WP_178119408.1 | Pseudomonas lalkuanensis | Gammaproteobacteria | Pseudomonadales | Pseudomonadaceae |
| gi\|QHC99364.1 | Pseudomonas sp. S04 | Gammaproteobacteria | Pseudomonadales | Pseudomonadaceae |
| gi\|WP_180984082.1 | Pseudomonas stutzeri | Gammaproteobacteria | Pseudomonadales | Pseudomonadaceae |
| gi\|WP_157883339.1 | Pseudomonas sp. ATCC 13867 | Gammaproteobacteria | Pseudomonadales | Pseudomonadaceae |
| gi\|PYG16112.1 | Pseudomonas sp. OV286 | Gammaproteobacteria | Pseudomonadales | Pseudomonadaceae |
| gi\|WP_149411095.1 | unclassified Pseudomonas | Gammaproteobacteria | Pseudomonadales | Pseudomonadaceae |
| gi\|WP_128606439.1 | Pseudomonas sp. ERMR1:02 | Gammaproteobacteria | Pseudomonadales | Pseudomonadaceae |
| gi\|ABP80656.1 | Pseudomonas stutzeri A1501 | Gammaproteobacteria | Pseudomonadales | Pseudomonadaceae |
| gi\|WP_177124254.1 | Pseudomonas gingeri | Gammaproteobacteria | Pseudomonadales | Pseudomonadaceae |
| gi\|WP_123333901.1 | Pseudomonas chlororaphis | Gammaproteobacteria | Pseudomonadales | Pseudomonadaceae |
| gi\|WP_141123505.1 | Pseudomonas veronii | Gammaproteobacteria | Pseudomonadales | Pseudomonadaceae |
| gi\|TRX73892.1 | Pseudomonas sp. DMKU_BBB3-04 | Gammaproteobacteria | Pseudomonadales | Pseudomonadaceae |
| gi\|WP_188865572.1 | Pseudomonas asuensis | Gammaproteobacteria | Pseudomonadales | Pseudomonadaceae |
| gi\|QFU13394.1 | Pseudomonas sp. THAF7b | Gammaproteobacteria | Pseudomonadales | Pseudomonadaceae |
| gi\|QHD06961.1 | Pseudomonas sp. R76 | Gammaproteobacteria | Pseudomonadales | Pseudomonadaceae |
| gi\|OXS21107.1 | Pseudomonas fluorescens | Gammaproteobacteria | Pseudomonadales | Pseudomonadaceae |
| gi\|WP_108094616.1 | unclassified Pseudomonas | Gammaproteobacteria | Pseudomonadales | Pseudomonadaceae |
| gi\|PAU52248.1 | Pseudomonas indica | Gammaproteobacteria | Pseudomonadales | Pseudomonadaceae |
| gi\|WP_150787621.1 | Pseudomonas fluorescens | Gammaproteobacteria | Pseudomonadales | Pseudomonadaceae |
| gi\|KPX74337.1 | Pseudomonas syringae pv. maculicola | Gammaproteobacteria | Pseudomonadales | Pseudomonadaceae |
| gi\|WP_187682317.1 | Pseudomonas lurida | Gammaproteobacteria | Pseudomonadales | Pseudomonadaceae |
| gi\|VVN49183.1 | Pseudomonas fluorescens | Gammaproteobacteria | Pseudomonadales | Pseudomonadaceae |
| gi\|WP_075932275.1 | unclassified Pseudomonas | Gammaproteobacteria | Pseudomonadales | Pseudomonadaceae |
| gi\|WP_084319225.1 | Pseudomonas migulae | Gammaproteobacteria | Pseudomonadales | Pseudomonadaceae |
| gi\|WP_156430112.1 | Pseudomonas agarici | Gammaproteobacteria | Pseudomonadales | Pseudomonadaceae |
| gi\|WP_019582254.1 | Pseudomonas mandelii | Gammaproteobacteria | Pseudomonadales | Pseudomonadaceae |
| gi\|WP_181130787.1 | Pseudomonas capeferrum | Gammaproteobacteria | Pseudomonadales | Pseudomonadaceae |
| gi\|WP_181101662.1 | Pseudomonas | Gammaproteobacteria | Pseudomonadales | Pseudomonadaceae |
| gi\|WP_158461088.1 | Pseudomonas fluorescens | Gammaproteobacteria | Pseudomonadales | Pseudomonadaceae |
| gi\|SFI90004.1 | Pseudomonas guineae | Gammaproteobacteria | Pseudomonadales | Pseudomonadaceae |
| gi\|ROM77759.1 | Pseudomonas brassicacearum | Gammaproteobacteria | Pseudomonadales | Pseudomonadaceae |
| gi\|WP_028630106.1 | Pseudomonas resinovorans | Gammaproteobacteria | Pseudomonadales | Pseudomonadaceae |
| gi\|WP_150644387.1 | Pseudomonas fluorescens | Gammaproteobacteria | Pseudomonadales | Pseudomonadaceae |
| gi\|WP_074861989.1 | Pseudomonas agarici | Gammaproteobacteria | Pseudomonadales | Pseudomonadaceae |
| gi\|WP_173178151.1 | Pseudomonas sp. TUM18999 | Gammaproteobacteria | Pseudomonadales | Pseudomonadaceae |
| gi\|WP_186685733.1 | Pseudomonas sp. RW8P3 | Gammaproteobacteria | Pseudomonadales | Pseudomonadaceae |
| gi\|WP_084380930.1 | Pseudomonas mucidolens | Gammaproteobacteria | Pseudomonadales | Pseudomonadaceae |
| gi\|WP_187673107.1 | Pseudomonas carbonaria | Gammaproteobacteria | Pseudomonadales | Pseudomonadaceae |
| gi\|SEM88612.1 | Pseudomonas sp. ok272 | Gammaproteobacteria | Pseudomonadales | Pseudomonadaceae |
| gi\|WP_016493286.1 | Pseudomonas resinovorans | Gammaproteobacteria | Pseudomonadales | Pseudomonadaceae |
| gi\|WP_095944218.1 | Pseudomonas sp. ACN8 | Gammaproteobacteria | Pseudomonadales | Pseudomonadaceae |
| gi\|WP_150346886.1 | Pseudomonas | Gammaproteobacteria | Pseudomonadales | Pseudomonadaceae |

| | | | | |
|---|---|---|---|---|
| gi\|WP_099237359.1 | Pseudomonas sp. ICMP 460 | Gammaproteobacteria | Pseudomonadales | Pseudomonadaceae |
| gi\|WP_122314100.1 | Pseudomonas cichorii | Gammaproteobacteria | Pseudomonadales | Pseudomonadaceae |
| gi\|WP_042128038.1 | Pseudomonas japonica | Gammaproteobacteria | Pseudomonadales | Pseudomonadaceae |
| gi\|SFP28399.1 | Pseudomonas sagittaria | Gammaproteobacteria | Pseudomonadales | Pseudomonadaceae |
| gi\|WP_045196971.1 | unclassified Pseudomonas | Gammaproteobacteria | Pseudomonadales | Pseudomonadaceae |
| gi\|WP_192068957.1 | Pseudomonas coleopterorum | Gammaproteobacteria | Pseudomonadales | Pseudomonadaceae |
| gi\|WP_190424948.1 | Pseudomonas typographi | Gammaproteobacteria | Pseudomonadales | Pseudomonadaceae |
| gi\|WP_179058098.1 | Pseudomonas taiwanensis | Gammaproteobacteria | Pseudomonadales | Pseudomonadaceae |
| gi\|WP_186555553.1 | Pseudomonas sp. SWRI10 | Gammaproteobacteria | Pseudomonadales | Pseudomonadaceae |
| gi\|WP_090343148.1 | Pseudomonas guariconensis | Gammaproteobacteria | Pseudomonadales | Pseudomonadaceae |
| gi\|CDZ93561.1 | Pseudomonas saudiphocaensis | Gammaproteobacteria | Pseudomonadales | Pseudomonadaceae |
| gi\|WP_081006023.1 | Pseudomonas fuscovaginae | Gammaproteobacteria | Pseudomonadales | Pseudomonadaceae |
| gi\|WP_161759953.1 | unclassified Pseudomonas | Gammaproteobacteria | Pseudomonadales | Pseudomonadaceae |
| gi\|WP_190832068.1 | Pseudomonas sp. JM0905a | Gammaproteobacteria | Pseudomonadales | Pseudomonadaceae |
| gi\|WP_185267985.1 | Pseudomonas xiamenensis | Gammaproteobacteria | Pseudomonadales | Pseudomonadaceae |
| gi\|SFM69503.1 | Rugamonas rubra | Gammaproteobacteria | Pseudomonadales | Pseudomonadaceae |
| gi\|WP_058603509.1 | Pseudomonas | Gammaproteobacteria | Pseudomonadales | Pseudomonadaceae |
| gi\|WP_150804138.1 | Pseudomonas fluorescens | Gammaproteobacteria | Pseudomonadales | Pseudomonadaceae |
| gi\|WP_172149963.1 | Pseudomonas sp. LAM-KW06 | Gammaproteobacteria | Pseudomonadales | Pseudomonadaceae |
| gi\|WP_122453514.1 | Pseudomonas viridiflava | Gammaproteobacteria | Pseudomonadales | Pseudomonadaceae |
| gi\|WP_064388722.1 | Pseudomonas sp. RIT-PI-r | Gammaproteobacteria | Pseudomonadales | Pseudomonadaceae |
| gi\|WP_108239506.1 | unclassified Pseudomonas | Gammaproteobacteria | Pseudomonadales | Pseudomonadaceae |
| gi\|WP_084596162.1 | Pseudomonas massiliensis | Gammaproteobacteria | Pseudomonadales | Pseudomonadaceae |
| gi\|WP_123589180.1 | Pseudomonas fluorescens | Gammaproteobacteria | Pseudomonadales | Pseudomonadaceae |
| gi\|WP_056839392.1 | Pseudomonas sp. Leaf127 | Gammaproteobacteria | Pseudomonadales | Pseudomonadaceae |
| gi\|WP_191485851.1 | Pseudomonas sp. FEN | Gammaproteobacteria | Pseudomonadales | Pseudomonadaceae |
| gi\|WP_146426532.1 | Pseudomonas saxonica | Gammaproteobacteria | Pseudomonadales | Pseudomonadaceae |
| gi\|WP_196176087.1 | Pseudomonas fulva | Gammaproteobacteria | Pseudomonadales | Pseudomonadaceae |
| gi\|WP_122315264.1 | Pseudomonas cichorii | Gammaproteobacteria | Pseudomonadales | Pseudomonadaceae |
| gi\|WP_090902910.1 | Azotobacter beijerinckii | Gammaproteobacteria | Pseudomonadales | Pseudomonadaceae |
| gi\|WP_133324559.1 | Pseudomonas putida group | Gammaproteobacteria | Pseudomonadales | Pseudomonadaceae |
| gi\|WP_085582754.1 | unclassified Pseudomonas | Gammaproteobacteria | Pseudomonadales | Pseudomonadaceae |
| gi\|RMP59128.1 | Pseudomonas marginalis pv. marginalis | Gammaproteobacteria | Pseudomonadales | Pseudomonadaceae |
| gi\|WP_083329851.1 | Pseudomonas argentinensis | Gammaproteobacteria | Pseudomonadales | Pseudomonadaceae |
| gi\|WP_022642041.1 | Pseudomonas | Gammaproteobacteria | Pseudomonadales | Pseudomonadaceae |
| gi\|WP_055136703.1 | Pseudomonas corrugata | Gammaproteobacteria | Pseudomonadales | Pseudomonadaceae |
| gi\|RMO62107.1 | Pseudomonas marginalis pv. marginalis | Gammaproteobacteria | Pseudomonadales | Pseudomonadaceae |
| gi\|WP_116552621.1 | Pseudomonas sp. SDI | Gammaproteobacteria | Pseudomonadales | Pseudomonadaceae |
| gi\|WP_105642401.1 | Pseudomonas sp. MYb187 | Gammaproteobacteria | Pseudomonadales | Pseudomonadaceae |
| gi\|WP_145190122.1 | Pseudomonas sp. URMO17WK12:I11 | Gammaproteobacteria | Pseudomonadales | Pseudomonadaceae |
| gi\|RBL67106.1 | Pseudomonas sp. MWU13-2625 | Gammaproteobacteria | Pseudomonadales | Pseudomonadaceae |
| gi\|RZI87823.1 | Pseudomonas sp. | Gammaproteobacteria | Pseudomonadales | Pseudomonadaceae |
| gi\|WP_028238917.1 | Pseudomonas azotifigens | Gammaproteobacteria | Pseudomonadales | Pseudomonadaceae |

| | | | | |
|---|---|---|---|---|
| gi\|WP_123345058.1 | Pseudomonas brassicacearum | Gammaproteobacteria | Pseudomonadales | Pseudomonadaceae |
| gi\|ESW58412.1 | Pseudomonas fluorescens BBc6R8 | Gammaproteobacteria | Pseudomonadales | Pseudomonadaceae |
| gi\|WP_042730025.1 | Pseudomonas fluorescens | Gammaproteobacteria | Pseudomonadales | Pseudomonadaceae |
| gi\|WP_172792110.1 | Pseudomonas sp. B14-6 | Gammaproteobacteria | Pseudomonadales | Pseudomonadaceae |
| gi\|WP_188982192.1 | Pseudomonas matsuisoli | Gammaproteobacteria | Pseudomonadales | Pseudomonadaceae |
| gi\|WP_083392435.1 | Pseudomonas bauzanensis | Gammaproteobacteria | Pseudomonadales | Pseudomonadaceae |
| gi\|WP_131173249.1 | Pseudomonas dryadis | Gammaproteobacteria | Pseudomonadales | Pseudomonadaceae |
| gi\|WP_102052028.1 | Pseudomonas sp. FFUP_PS_473 | Gammaproteobacteria | Pseudomonadales | Pseudomonadaceae |
| gi\|WP_179113304.1 | Pseudomonas sp. ABC1 | Gammaproteobacteria | Pseudomonadales | Pseudomonadaceae |
| gi\|WP_087515682.1 | Pseudomonas sp. M30-35 | Gammaproteobacteria | Pseudomonadales | Pseudomonadaceae |
| gi\|WP_122251058.1 | Pseudomonas marginalis | Gammaproteobacteria | Pseudomonadales | Pseudomonadaceae |
| gi\|WP_152225316.1 | Pseudomonas sp. SCB32 | Gammaproteobacteria | Pseudomonadales | Pseudomonadaceae |
| gi\|WP_020297485.1 | Pseudomonas sp. CF161 | Gammaproteobacteria | Pseudomonadales | Pseudomonadaceae |
| gi\|WP_090262411.1 | Pseudomonas panipatensis | Gammaproteobacteria | Pseudomonadales | Pseudomonadaceae |
| gi\|WP_093419202.1 | unclassified Pseudomonas | Gammaproteobacteria | Pseudomonadales | Pseudomonadaceae |
| gi\|WP_095109430.1 | Pseudomonas sp. Irchel 3E20 | Gammaproteobacteria | Pseudomonadales | Pseudomonadaceae |
| gi\|WP_169428671.1 | Pseudomonas fluorescens | Gammaproteobacteria | Pseudomonadales | Pseudomonadaceae |
| gi\|WP_184682338.1 | Pseudomonas fluvialis | Gammaproteobacteria | Pseudomonadales | Pseudomonadaceae |
| gi\|WP_122372507.1 | Pseudomonas cichorii | Gammaproteobacteria | Pseudomonadales | Pseudomonadaceae |
| gi\|WP_122164454.1 | Pseudomonas zhaodongensis | Gammaproteobacteria | Pseudomonadales | Pseudomonadaceae |
| gi\|WP_021445551.1 | Pseudomonas sp. EGD-AK9 | Gammaproteobacteria | Pseudomonadales | Pseudomonadaceae |
| gi\|WP_136491593.1 | Pseudomonas sp. A-1 | Gammaproteobacteria | Pseudomonadales | Pseudomonadaceae |
| gi\|WP_069899650.1 | Pseudomonas | Gammaproteobacteria | Pseudomonadales | Pseudomonadaceae |
| gi\|WP_116550675.1 | Pseudomonas sp. SDI | Gammaproteobacteria | Pseudomonadales | Pseudomonadaceae |
| gi\|WP_131188970.1 | Pseudomonas kirkiae | Gammaproteobacteria | Pseudomonadales | Pseudomonadaceae |
| gi\|PZP22837.1 | Pseudomonas kuykendallii | Gammaproteobacteria | Pseudomonadales | Pseudomonadaceae |
| gi\|WP_192209955.1 | Pseudomonas sp. PDM22 | Gammaproteobacteria | Pseudomonadales | Pseudomonadaceae |
| gi\|WP_160343642.1 | Pseudomonas sp. R-22-3w-18 | Gammaproteobacteria | Pseudomonadales | Pseudomonadaceae |
| gi\|WP_103457790.1 | Pseudomonas stutzeri | Gammaproteobacteria | Pseudomonadales | Pseudomonadaceae |
| gi\|WP_103400128.1 | Pseudomonas sp. FW300-N1A1 | Gammaproteobacteria | Pseudomonadales | Pseudomonadaceae |
| gi\|WP_191487923.1 | Pseudomonas sp. FEN | Gammaproteobacteria | Pseudomonadales | Pseudomonadaceae |
| gi\|RZI70110.1 | Pseudomonas sp. | Gammaproteobacteria | Pseudomonadales | Pseudomonadaceae |
| gi\|WP_045425289.1 | Pseudomonas stutzeri | Gammaproteobacteria | Pseudomonadales | Pseudomonadaceae |
| gi\|WP_166650976.1 | Pseudomonas sp. LP_7_YM | Gammaproteobacteria | Pseudomonadales | Pseudomonadaceae |
| gi\|WP_012700871.1 | Azotobacter vinelandii | Gammaproteobacteria | Pseudomonadales | Pseudomonadaceae |
| gi\|WP_079203330.1 | Pseudomonas sp. CC6-YY-74 | Gammaproteobacteria | Pseudomonadales | Pseudomonadaceae |
| gi\|WP_165670644.1 | Pseudomonas otitidis | Gammaproteobacteria | Pseudomonadales | Pseudomonadaceae |
| gi\|WP_135291425.1 | Pseudomonas kairouanensis | Gammaproteobacteria | Pseudomonadales | Pseudomonadaceae |
| gi\|WP_010488867.1 | Pseudomonas sp. S9 | Gammaproteobacteria | Pseudomonadales | Pseudomonadaceae |
| gi\|WP_149087614.1 | Pseudomonas prosekii | Gammaproteobacteria | Pseudomonadales | Pseudomonadaceae |
| gi\|WP_179526759.1 | Pseudomonas composti | Gammaproteobacteria | Pseudomonadales | Pseudomonadaceae |
| gi\|WP_119892118.1 | Pseudomonas sp. K2W31S-8 | Gammaproteobacteria | Pseudomonadales | Pseudomonadaceae |
| gi\|SDS83843.1 | Pseudomonas oryzae | Gammaproteobacteria | Pseudomonadales | Pseudomonadaceae |

| | | | | |
|---|---|---|---|---|
| gi\|WP_043310789.1 | Pseudomonas sp. ML96 | Gammaproteobacteria | Pseudomonadales | Pseudomonadaceae |
| gi\|QEY62622.1 | Pseudomonas lalkuanensis | Gammaproteobacteria | Pseudomonadales | Pseudomonadaceae |
| gi\|WP_070880748.1 | Pseudomonas seleniipraecipitans | Gammaproteobacteria | Pseudomonadales | Pseudomonadaceae |
| gi\|WP_042935138.1 | Pseudomonas gingeri | Gammaproteobacteria | Pseudomonadales | Pseudomonadaceae |
| gi\|WP_076426794.1 | Pseudomonas alcaligenes | Gammaproteobacteria | Pseudomonadales | Pseudomonadaceae |
| gi\|WP_099526187.1 | Pseudomonas sediminis | Gammaproteobacteria | Pseudomonadales | Pseudomonadaceae |
| gi\|SDS95935.1 | Pseudomonas litoralis | Gammaproteobacteria | Pseudomonadales | Pseudomonadaceae |
| gi\|WP_070886926.1 | Pseudomonas argentinensis | Gammaproteobacteria | Pseudomonadales | Pseudomonadaceae |
| gi\|WP_192101823.1 | Pseudomonas syringae | Gammaproteobacteria | Pseudomonadales | Pseudomonadaceae |
| gi\|WP_039802456.1 | Azotobacter chroococcum | Gammaproteobacteria | Pseudomonadales | Pseudomonadaceae |
| gi\|WP_053155523.1 | Pseudomonas sp. Pf153 | Gammaproteobacteria | Pseudomonadales | Pseudomonadaceae |
| gi\|OXM41437.1 | Pseudomonas fluvialis | Gammaproteobacteria | Pseudomonadales | Pseudomonadaceae |
| gi\|WP_083350074.1 | Pseudomonas umsongensis | Gammaproteobacteria | Pseudomonadales | Pseudomonadaceae |
| gi\|WP_177103252.1 | Pseudomonas gingeri | Gammaproteobacteria | Pseudomonadales | Pseudomonadaceae |
| gi\|WP_090445481.1 | Pseudomonas benzenivorans | Gammaproteobacteria | Pseudomonadales | Pseudomonadaceae |
| gi\|WP_081563995.1 | Pseudomonas sp. Bc-h | Gammaproteobacteria | Pseudomonadales | Pseudomonadaceae |
| gi\|WP_110725462.1 | unclassified Pseudomonas | Gammaproteobacteria | Pseudomonadales | Pseudomonadaceae |
| gi\|KAF1011300.1 | Pseudomonas fluorescens | Gammaproteobacteria | Pseudomonadales | Pseudomonadaceae |
| gi\|WP_192101825.1 | Pseudomonas syringae | Gammaproteobacteria | Pseudomonadales | Pseudomonadaceae |
| gi\|KAF1031284.1 | Pseudomonas sp. | Gammaproteobacteria | Pseudomonadales | Pseudomonadaceae |
| gi\|WP_193074657.1 | Pseudomonas sp. FME51 | Gammaproteobacteria | Pseudomonadales | Pseudomonadaceae |
| gi\|WP_148926294.1 | Pseudomonas stutzeri | Gammaproteobacteria | Pseudomonadales | Pseudomonadaceae |
| gi\|WP_120994128.1 | Pseudomonas urumqiensis | Gammaproteobacteria | Pseudomonadales | Pseudomonadaceae |
| gi\|WP_166590698.1 | Pseudomonas sp. BC115LW | Gammaproteobacteria | Pseudomonadales | Pseudomonadaceae |
| gi\|WP_011533586.1 | Pseudomonas entomophila | Gammaproteobacteria | Pseudomonadales | Pseudomonadaceae |
| gi\|WP_150712339.1 | Pseudomonas fluorescens | Gammaproteobacteria | Pseudomonadales | Pseudomonadaceae |
| gi\|KAF1068878.1 | Pseudomonas citronellolis | Gammaproteobacteria | Pseudomonadales | Pseudomonadaceae |
| gi\|WP_076584335.1 | Pseudomonas alcaligenes | Gammaproteobacteria | Pseudomonadales | Pseudomonadaceae |
| gi\|WP_172433740.1 | Pseudomonas otitidis | Gammaproteobacteria | Pseudomonadales | Pseudomonadaceae |
| gi\|WP_083183952.1 | Pseudomonas floridensis | Gammaproteobacteria | Pseudomonadales | Pseudomonadaceae |
| gi\|WP_065895731.1 | Pseudomonas | Gammaproteobacteria | Pseudomonadales | Pseudomonadaceae |
| gi\|WP_011912458.1 | Pseudomonas stutzeri | Gammaproteobacteria | Pseudomonadales | Pseudomonadaceae |
| gi\|WP_182832548.1 | Pseudomonas sp. SR9 | Gammaproteobacteria | Pseudomonadales | Pseudomonadaceae |
| gi\|SFQ05594.1 | Pseudomonas borbori | Gammaproteobacteria | Pseudomonadales | Pseudomonadaceae |
| gi\|WP_090502191.1 | Pseudomonas borbori | Gammaproteobacteria | Pseudomonadales | Pseudomonadaceae |
| gi\|WP_116887480.1 | Pseudomonas parafulva | Gammaproteobacteria | Pseudomonadales | Pseudomonadaceae |
| gi\|WP_183087770.1 | Pseudomonas sp. UL070 | Gammaproteobacteria | Pseudomonadales | Pseudomonadaceae |
| gi\|WP_188390290.1 | Pseudomonas fluvialis | Gammaproteobacteria | Pseudomonadales | Pseudomonadaceae |
| gi\|WP_058069271.1 | Pseudomonas sp. TTU2014-080ASC | Gammaproteobacteria | Pseudomonadales | Pseudomonadaceae |
| gi\|WP_045490603.1 | Pseudomonas sp. StFLB209 | Gammaproteobacteria | Pseudomonadales | Pseudomonadaceae |
| gi\|WP_125862296.1 | Pseudomonas xanthomarina | Gammaproteobacteria | Pseudomonadales | Pseudomonadaceae |
| gi\|WP_104728715.1 | Pseudomonas oleovorans | Gammaproteobacteria | Pseudomonadales | Pseudomonadaceae |
| gi\|WP_090243516.1 | Pseudomonas guineae | Gammaproteobacteria | Pseudomonadales | Pseudomonadaceae |

| | | | | |
|---|---|---|---|---|
| gi\|WP_139199050.1 | Pseudomonas panipatensis | Gammaproteobacteria | Pseudomonadales | Pseudomonadaceae |
| gi\|WP_099235648.1 | Pseudomonas sp. ICMP 460 | Gammaproteobacteria | Pseudomonadales | Pseudomonadaceae |
| gi\|WP_108107959.1 | Pseudomonas mangrovi | Gammaproteobacteria | Pseudomonadales | Pseudomonadaceae |
| gi\|WP_042551780.1 | Pseudomonas | Gammaproteobacteria | Pseudomonadales | Pseudomonadaceae |
| gi\|WP_153919439.1 | Pseudomonas sp. JG-B | Gammaproteobacteria | Pseudomonadales | Pseudomonadaceae |
| gi\|WP_095941539.1 | Pseudomonas sp. HAR-UPW-AIA-41 | Gammaproteobacteria | Pseudomonadales | Pseudomonadaceae |
| gi\|WP_179113314.1 | Pseudomonas sp. ABC1 | Gammaproteobacteria | Pseudomonadales | Pseudomonadaceae |
| gi\|WP_079203332.1 | Pseudomonas sp. CC6-YY-74 | Gammaproteobacteria | Pseudomonadales | Pseudomonadaceae |
| gi\|WP_122459431.1 | Pseudomonas viridiflava | Gammaproteobacteria | Pseudomonadales | Pseudomonadaceae |
| gi\|KPX27570.1 | Pseudomonas syringae pv. delphinii | Gammaproteobacteria | Pseudomonadales | Pseudomonadaceae |
| gi\|WP_090445473.1 | Pseudomonas benzenivorans | Gammaproteobacteria | Pseudomonadales | Pseudomonadaceae |
| gi\|WP_093465605.1 | Pseudomonas sp. NFR16 | Gammaproteobacteria | Pseudomonadales | Pseudomonadaceae |
| gi\|WP_188982191.1 | Pseudomonas matsuisoli | Gammaproteobacteria | Pseudomonadales | Pseudomonadaceae |
| gi\|TQL05588.1 | Pseudomonas sp. SLBN-26 | Gammaproteobacteria | Pseudomonadales | Pseudomonadaceae |
| gi\|WP_146180698.1 | unclassified Pseudomonas | Gammaproteobacteria | Pseudomonadales | Pseudomonadaceae |
| gi\|WP_153326547.1 | Pseudomonas helleri | Gammaproteobacteria | Pseudomonadales | Pseudomonadaceae |
| gi\|WP_173180111.1 | Pseudomonas sp. TUM18999 | Gammaproteobacteria | Pseudomonadales | Pseudomonadaceae |
| gi\|WP_147170963.1 | Pseudomonas sp. SJZ079 | Gammaproteobacteria | Pseudomonadales | Pseudomonadaceae |
| gi\|WP_165594223.1 | Pseudomonas stutzeri | Gammaproteobacteria | Pseudomonadales | Pseudomonadaceae |
| gi\|WP_192318502.1 | Pseudomonas sp. PDM16 | Gammaproteobacteria | Pseudomonadales | Pseudomonadaceae |
| gi\|WP_061240906.1 | Pseudomonas composti | Gammaproteobacteria | Pseudomonadales | Pseudomonadaceae |
| gi\|WP_057008855.1 | Pseudomonas trivialis | Gammaproteobacteria | Pseudomonadales | Pseudomonadaceae |
| gi\|WP_159890285.1 | Pseudomonas sp. LD120 | Gammaproteobacteria | Pseudomonadales | Pseudomonadaceae |
| gi\|WP_026012989.1 | Pseudomonas agarici | Gammaproteobacteria | Pseudomonadales | Pseudomonadaceae |
| gi\|WP_150644385.1 | Pseudomonas fluorescens | Gammaproteobacteria | Pseudomonadales | Pseudomonadaceae |
| gi\|WP_170049133.1 | Pseudomonas sp. WS 5011 | Gammaproteobacteria | Pseudomonadales | Pseudomonadaceae |
| gi\|KHO65445.1 | Pseudomonas flexibilis | Gammaproteobacteria | Pseudomonadales | Pseudomonadaceae |
| gi\|WP_076423589.1 | Pseudomonas alcaligenes | Gammaproteobacteria | Pseudomonadales | Pseudomonadaceae |
| gi\|WP_179111714.1 | Pseudomonas sp. ABC1 | Gammaproteobacteria | Pseudomonadales | Pseudomonadaceae |
| gi\|WP_122840579.1 | Pseudomonas viridiflava | Gammaproteobacteria | Pseudomonadales | Pseudomonadaceae |
| gi\|WP_024309508.1 | Pseudomonas sp. P818 | Gammaproteobacteria | Pseudomonadales | Pseudomonadaceae |
| gi\|PZP24914.1 | Pseudomonas kuykendallii | Gammaproteobacteria | Pseudomonadales | Pseudomonadaceae |
| gi\|SFQ87050.1 | Pseudomonas formosensis | Gammaproteobacteria | Pseudomonadales | Pseudomonadaceae |
| gi\|WP_122772214.1 | Pseudomonas viridiflava | Gammaproteobacteria | Pseudomonadales | Pseudomonadaceae |
| gi\|WP_137973980.1 | Pseudomonas sp. F(2018) | Gammaproteobacteria | Pseudomonadales | Pseudomonadaceae |
| gi\|WP_183088828.1 | Pseudomonas sp. UL070 | Gammaproteobacteria | Pseudomonadales | Pseudomonadaceae |
| gi\|WP_044499182.1 | Pseudomonas saudimassiliensis | Gammaproteobacteria | Pseudomonadales | Pseudomonadaceae |
| gi\|WP_193681099.1 | Pseudomonas lopnurensis | Gammaproteobacteria | Pseudomonadales | Pseudomonadaceae |
| gi\|WP_153326434.1 | Pseudomonas | Gammaproteobacteria | Pseudomonadales | Pseudomonadaceae |
| gi\|WP_172150292.1 | Pseudomonas sp. LAM-KW06 | Gammaproteobacteria | Pseudomonadales | Pseudomonadaceae |
| gi\|WP_080890638.1 | Pseudomonas stutzeri | Gammaproteobacteria | Pseudomonadales | Pseudomonadaceae |
| gi\|WP_125861343.1 | Pseudomonas entomophila | Gammaproteobacteria | Pseudomonadales | Pseudomonadaceae |
| gi\|WP_153327631.1 | Pseudomonas helleri | Gammaproteobacteria | Pseudomonadales | Pseudomonadaceae |

| | | | | |
|---|---|---|---|---|
| gi\|WP_131188412.1 | Pseudomonas kirkiae | Gammaproteobacteria | Pseudomonadales | Pseudomonadaceae |
| gi\|WP_133774444.1 | Pseudomonas graminis | Gammaproteobacteria | Pseudomonadales | Pseudomonadaceae |
| gi\|SDH85395.1 | Pseudomonas panipatensis | Gammaproteobacteria | Pseudomonadales | Pseudomonadaceae |
| gi\|WP_192331285.1 | Pseudomonas sp. PDM14 | Gammaproteobacteria | Pseudomonadales | Pseudomonadaceae |
| gi\|WP_081711604.1 | Pseudomonas alcaligenes | Gammaproteobacteria | Pseudomonadales | Pseudomonadaceae |
| gi\|WP_157825252.1 | Pseudomonas pharmacofabricae | Gammaproteobacteria | Pseudomonadales | Pseudomonadaceae |
| gi\|WP_111264246.1 | Pseudomonas sp. 57B-090624 | Gammaproteobacteria | Pseudomonadales | Pseudomonadaceae |
| gi\|WP_099526191.1 | Pseudomonas sediminis | Gammaproteobacteria | Pseudomonadales | Pseudomonadaceae |
| gi\|WP_160089036.1 | Pseudomonas sp. 9AZ | Gammaproteobacteria | Pseudomonadales | Pseudomonadaceae |
| gi\|WP_179554902.1 | Pseudomonas oleovorans | Gammaproteobacteria | Pseudomonadales | Pseudomonadaceae |
| gi\|WP_088192954.1 | Pseudomonas sp. A46 | Gammaproteobacteria | Pseudomonadales | Pseudomonadaceae |
| gi\|WP_133539616.1 | Thiopseudomonas denitrificans | Gammaproteobacteria | Pseudomonadales | Pseudomonadaceae |
| gi\|WP_119687215.1 | Pseudomonas putida | Gammaproteobacteria | Pseudomonadales | Pseudomonadaceae |
| gi\|WP_187803855.1 | Pseudomonas alcaligenes | Gammaproteobacteria | Pseudomonadales | Pseudomonadaceae |
| gi\|WP_090416059.1 | Pseudomonas jinjuensis | Gammaproteobacteria | Pseudomonadales | Pseudomonadaceae |
| gi\|WP_166571818.1 | Pseudomonas sp. R5(2019) | Gammaproteobacteria | Pseudomonadales | Pseudomonadaceae |
| gi\|WP_196473526.1 | Pseudomonas sp. LMG 31766 | Gammaproteobacteria | Pseudomonadales | Pseudomonadaceae |
| gi\|WP_105645756.1 | Pseudomonas sp. MYb185 | Gammaproteobacteria | Pseudomonadales | Pseudomonadaceae |
| gi\|WP_110973246.1 | Pseudomonas huaxiensis | Gammaproteobacteria | Pseudomonadales | Pseudomonadaceae |
| gi\|WP_188982190.1 | Pseudomonas matsuisoli | Gammaproteobacteria | Pseudomonadales | Pseudomonadaceae |
| gi\|WP_108486022.1 | unclassified Pseudomonas | Gammaproteobacteria | Pseudomonadales | Pseudomonadaceae |
| gi\|WP_112898382.1 | Pseudomonas | Gammaproteobacteria | Pseudomonadales | Pseudomonadaceae |
| gi\|WP_159994908.1 | Pseudomonas | Gammaproteobacteria | Pseudomonadales | Pseudomonadaceae |
| gi\|WP_082107735.1 | Pseudomonas veronii | Gammaproteobacteria | Pseudomonadales | Pseudomonadaceae |
| gi\|WP_166362534.1 | Pseudomonas sp. PS24 | Gammaproteobacteria | Pseudomonadales | Pseudomonadaceae |
| gi\|WP_083329852.1 | Pseudomonas argentinensis | Gammaproteobacteria | Pseudomonadales | Pseudomonadaceae |
| gi\|EGH95595.1 | Pseudomonas amygdali pv. lachrymans str. M302278 | Gammaproteobacteria | Pseudomonadales | Pseudomonadaceae |
| gi\|WP_102894013.1 | Pseudomonas stutzeri | Gammaproteobacteria | Pseudomonadales | Pseudomonadaceae |
| gi\|WP_173203160.1 | Pseudomonas campi | Gammaproteobacteria | Pseudomonadales | Pseudomonadaceae |
| gi\|WP_110614100.1 | Pseudomonas sp. OV467 | Gammaproteobacteria | Pseudomonadales | Pseudomonadaceae |
| gi\|WP_053527731.1 | Pseudomonas stutzeri | Gammaproteobacteria | Pseudomonadales | Pseudomonadaceae |
| gi\|KPY33959.1 | Pseudomonas syringae pv. primulae | Gammaproteobacteria | Pseudomonadales | Pseudomonadaceae |
| gi\|WP_047529603.1 | unclassified Pseudomonas | Gammaproteobacteria | Pseudomonadales | Pseudomonadaceae |
| gi\|WP_168425782.1 | Pseudomonas sp. SST3 | Gammaproteobacteria | Pseudomonadales | Pseudomonadaceae |
| gi\|WP_125859178.1 | Pseudomonas entomophila | Gammaproteobacteria | Pseudomonadales | Pseudomonadaceae |
| gi\|WP_103437625.1 | Pseudomonas putida | Gammaproteobacteria | Pseudomonadales | Pseudomonadaceae |
| gi\|WP_192209951.1 | Pseudomonas sp. PDM22 | Gammaproteobacteria | Pseudomonadales | Pseudomonadaceae |
| gi\|WP_183166079.1 | Azomonas macrocytogenes | Gammaproteobacteria | Pseudomonadales | Pseudomonadaceae |
| gi\|WP_160343644.1 | Pseudomonas sp. R-22-3w-18 | Gammaproteobacteria | Pseudomonadales | Pseudomonadaceae |
| gi\|OYT96989.1 | Pseudomonas sp. PGPPP3 | Gammaproteobacteria | Pseudomonadales | Pseudomonadaceae |
| gi\|WP_102052326.1 | Pseudomonas sp. FFUP_PS_473 | Gammaproteobacteria | Pseudomonadales | Pseudomonadaceae |
| gi\|WP_090404734.1 | Pseudomonas grimontii | Gammaproteobacteria | Pseudomonadales | Pseudomonadaceae |
| gi\|WP_017939754.1 | Pseudomonas thermotolerans | Gammaproteobacteria | Pseudomonadales | Pseudomonadaceae |

| | | | | |
|---|---|---|---|---|
| gi\|SFP05408.1 | Pseudomonas borbori | Gammaproteobacteria | Pseudomonadales | Pseudomonadaceae |
| gi\|WP_196166266.1 | Pseudomonas monteilii | Gammaproteobacteria | Pseudomonadales | Pseudomonadaceae |
| gi\|WP_181418713.1 | Pseudomonas alcaligenes | Gammaproteobacteria | Pseudomonadales | Pseudomonadaceae |
| gi\|WP_109511694.1 | Pseudomonas ovata | Gammaproteobacteria | Pseudomonadales | Pseudomonadaceae |
| gi\|WP_192325967.1 | Pseudomonas sp. PDM14 | Gammaproteobacteria | Pseudomonadales | Pseudomonadaceae |
| gi\|WP_061902786.1 | Pseudomonas alcaligenes | Gammaproteobacteria | Pseudomonadales | Pseudomonadaceae |
| gi\|WP_003292558.1 | Pseudomonas stutzeri | Gammaproteobacteria | Pseudomonadales | Pseudomonadaceae |
| gi\|WP_108487764.1 | unclassified Pseudomonas | Gammaproteobacteria | Pseudomonadales | Pseudomonadaceae |
| gi\|WP_166591613.1 | Pseudomonas | Gammaproteobacteria | Pseudomonadales | Pseudomonadaceae |
| gi\|BCA26911.1 | Pseudomonas otitidis | Gammaproteobacteria | Pseudomonadales | Pseudomonadaceae |
| gi\|WP_153326562.1 | Pseudomonas helleri | Gammaproteobacteria | Pseudomonadales | Pseudomonadaceae |
| gi\|SDV05489.1 | Pseudomonas rhodesiae | Gammaproteobacteria | Pseudomonadales | Pseudomonadaceae |
| gi\|PIA66842.1 | Pseudomonas sediminis | Gammaproteobacteria | Pseudomonadales | Pseudomonadaceae |
| gi\|WP_102052029.1 | Pseudomonas sp. FFUP_PS_473 | Gammaproteobacteria | Pseudomonadales | Pseudomonadaceae |
| gi\|AJE21812.1 | Azotobacter chroococcum NCIMB 8003 | Gammaproteobacteria | Pseudomonadales | Pseudomonadaceae |
| gi\|WP_104739092.1 | Pseudomonas oceani | Gammaproteobacteria | Pseudomonadales | Pseudomonadaceae |
| gi\|RMR61658.1 | Pseudomonas cichorii | Gammaproteobacteria | Pseudomonadales | Pseudomonadaceae |
| gi\|WP_119142520.1 | Pseudomonas reidholzensis | Gammaproteobacteria | Pseudomonadales | Pseudomonadaceae |
| gi\|SUD78485.1 | Pseudomonas putida | Gammaproteobacteria | Pseudomonadales | Pseudomonadaceae |
| gi\|WP_179111713.1 | Pseudomonas sp. ABC1 | Gammaproteobacteria | Pseudomonadales | Pseudomonadaceae |
| gi\|WP_081672260.1 | Pseudomonas alcaligenes | Gammaproteobacteria | Pseudomonadales | Pseudomonadaceae |
| gi\|WP_102840727.1 | Pseudomonas stutzeri | Gammaproteobacteria | Pseudomonadales | Pseudomonadaceae |
| gi\|OYT94893.1 | Pseudomonas sp. PGPPP3 | Gammaproteobacteria | Pseudomonadales | Pseudomonadaceae |
| gi\|WP_125859188.1 | Pseudomonas entomophila | Gammaproteobacteria | Pseudomonadales | Pseudomonadaceae |
| gi\|WP_127163430.1 | Entomomonas moraniae | Gammaproteobacteria | Pseudomonadales | Pseudomonadaceae |
| gi\|WP_160080455.1 | Pseudomonas sp. 8AS | Gammaproteobacteria | Pseudomonadales | Pseudomonadaceae |
| gi\|WP_159890259.1 | Pseudomonas sp. LD120 | Gammaproteobacteria | Pseudomonadales | Pseudomonadaceae |
| gi\|WP_092389113.1 | Pseudomonas salegens | Gammaproteobacteria | Pseudomonadales | Pseudomonadaceae |
| gi\|WP_090198184.1 | Pseudomonas pohangensis | Gammaproteobacteria | Pseudomonadales | Pseudomonadaceae |
| gi\|WP_181102736.1 | Pseudomonas entomophila | Gammaproteobacteria | Pseudomonadales | Pseudomonadaceae |
| gi\|WP_161492358.1 | Pseudomonas frederiksbergensis | Gammaproteobacteria | Pseudomonadales | Pseudomonadaceae |
| gi\|WP_129932990.1 | Pseudomonas sp. SWI36 | Gammaproteobacteria | Pseudomonadales | Pseudomonadaceae |
| gi\|WP_095940191.1 | Pseudomonas sp. HAR-UPW-AIA-41 | Gammaproteobacteria | Pseudomonadales | Pseudomonadaceae |
| gi\|WP_096137308.1 | Pseudomonas syringae | Gammaproteobacteria | Pseudomonadales | Pseudomonadaceae |
| gi\|ABY99070.1 | Pseudomonas putida GB-1 | Gammaproteobacteria | Pseudomonadales | Pseudomonadaceae |
| gi\|WP_158190072.1 | Pseudomonas stutzeri | Gammaproteobacteria | Pseudomonadales | Pseudomonadaceae |
| gi\|AAN68940.1 | Pseudomonas putida KT2440 | Gammaproteobacteria | Pseudomonadales | Pseudomonadaceae |
| gi\|BBU44432.1 | Pseudomonas putida | Gammaproteobacteria | Pseudomonadales | Pseudomonadaceae |
| gi\|WP_153015415.1 | Ventosimonas gracilis | Gammaproteobacteria | Pseudomonadales | Ventosimonadaceae |
| gi\|WP_079214071.1 | Ventosimonas gracilis | Gammaproteobacteria | Pseudomonadales | Ventosimonadaceae |
| gi\|WP_072956781.1 | Vibrio gazogenes | Gammaproteobacteria | Vibrionales | Vibrionaceae |
| gi\|WP_105901067.1 | Vibrio gangliei | Gammaproteobacteria | Vibrionales | Vibrionaceae |
| gi\|WP_157371894.1 | Vibrio sp. MEBiC08052 | Gammaproteobacteria | Vibrionales | Vibrionaceae |

| | | | | |
|---|---|---|---|---|
| gi\|WP_164711841.1 | Vibrio zhugei | Gammaproteobacteria | Vibrionales | Vibrionaceae |
| gi\|WP_089139374.1 | Vibrio rumoiensis | Gammaproteobacteria | Vibrionales | Vibrionaceae |
| gi\|RCS70771.1 | Vibrio casei | Gammaproteobacteria | Vibrionales | Vibrionaceae |
| gi\|WP_082712215.1 | Vibrio tritonius | Gammaproteobacteria | Vibrionales | Vibrionaceae |
| gi\|WP_077313081.1 | Vibrio palustris | Gammaproteobacteria | Vibrionales | Vibrionaceae |
| gi\|WP_168796993.1 | Vibrio sp. H11 | Gammaproteobacteria | Vibrionales | Vibrionaceae |
| gi\|WP_115497193.1 | Dyella monticola | Gammaproteobacteria | Xanthomonadales | Rhodanobacteraceae |
| gi\|WP_188798358.1 | Dyella nitratireducens | Gammaproteobacteria | Xanthomonadales | Rhodanobacteraceae |
| gi\|EIM02926.1 | Rhodanobacter thiooxydans LCS2 | Gammaproteobacteria | Xanthomonadales | Rhodanobacteraceae |
| gi\|WP_192557536.1 | Dyella sp. 7MK23 | Gammaproteobacteria | Xanthomonadales | Rhodanobacteraceae |
| gi\|RDI97364.1 | Dyella solisilvae | Gammaproteobacteria | Xanthomonadales | Rhodanobacteraceae |
| gi\|WP_192676682.1 | Dyella sp. OAE510 | Gammaproteobacteria | Xanthomonadales | Rhodanobacteraceae |
| gi\|WP_184602245.1 | unclassified Rhodanobacter | Gammaproteobacteria | Xanthomonadales | Rhodanobacteraceae |
| gi\|WP_081500636.1 | Dyella japonica | Gammaproteobacteria | Xanthomonadales | Rhodanobacteraceae |
| gi\|WP_165418441.1 | Dyella sp. DHC06 | Gammaproteobacteria | Xanthomonadales | Rhodanobacteraceae |
| gi\|WP_146203587.1 | Fulvimonas soli | Gammaproteobacteria | Xanthomonadales | Rhodanobacteraceae |
| gi\|WP_090453397.1 | Dyella sp. OK004 | Gammaproteobacteria | Xanthomonadales | Rhodanobacteraceae |
| gi\|ODV15814.1 | Rhodanobacter sp. SCN 68-63 | Gammaproteobacteria | Xanthomonadales | Rhodanobacteraceae |
| gi\|WP_179476649.1 | Rhodanobacter sp. K2T2 | Gammaproteobacteria | Xanthomonadales | Rhodanobacteraceae |
| gi\|WP_128898452.1 | Dyella sp. M7H15-1 | Gammaproteobacteria | Xanthomonadales | Rhodanobacteraceae |
| gi\|WP_126674142.1 | Dyella dinghuensis | Gammaproteobacteria | Xanthomonadales | Rhodanobacteraceae |
| gi\|WP_182529880.1 | Dokdonella fugitiva | Gammaproteobacteria | Xanthomonadales | Rhodanobacteraceae |
| gi\|RUL68651.1 | Dyella choica | Gammaproteobacteria | Xanthomonadales | Rhodanobacteraceae |
| gi\|QAU23800.1 | Dyella sp. M7H15-1 | Gammaproteobacteria | Xanthomonadales | Rhodanobacteraceae |
| gi\|WP_168709582.1 | Rhodanobacter lindaniclasticus | Gammaproteobacteria | Xanthomonadales | Rhodanobacteraceae |
| gi\|WP_133949420.1 | Rhodanobacter sp. TND4FH1 | Gammaproteobacteria | Xanthomonadales | Rhodanobacteraceae |
| gi\|KJV35795.1 | Luteibacter yeojuensis | Gammaproteobacteria | Xanthomonadales | Rhodanobacteraceae |
| gi\|WP_090452713.1 | Dyella sp. OK004 | Gammaproteobacteria | Xanthomonadales | Rhodanobacteraceae |
| gi\|WP_166945470.1 | Luteibacter anthropi | Gammaproteobacteria | Xanthomonadales | Rhodanobacteraceae |
| gi\|WP_157511085.1 | Frateuria sp. Soil773 | Gammaproteobacteria | Xanthomonadales | Rhodanobacteraceae |
| gi\|WP_109126580.1 | Dyella sp. C11 | Gammaproteobacteria | Xanthomonadales | Rhodanobacteraceae |
| gi\|WP_158754975.1 | Dyella sp. S184 | Gammaproteobacteria | Xanthomonadales | Rhodanobacteraceae |
| gi\|WP_185754444.1 | Luteibacter sp. 9135 | Gammaproteobacteria | Xanthomonadales | Rhodanobacteraceae |
| gi\|WP_109126689.1 | Dyella sp. C11 | Gammaproteobacteria | Xanthomonadales | Rhodanobacteraceae |
| gi\|AIF45956.1 | Dyella japonica A8 | Gammaproteobacteria | Xanthomonadales | Rhodanobacteraceae |
| gi\|RDS84185.1 | Dyella psychrodurans | Gammaproteobacteria | Xanthomonadales | Rhodanobacteraceae |
| gi\|EIL86857.1 | Rhodanobacter sp. 115 | Gammaproteobacteria | Xanthomonadales | Rhodanobacteraceae |
| gi\|EIL86832.1 | Rhodanobacter sp. 115 | Gammaproteobacteria | Xanthomonadales | Rhodanobacteraceae |
| gi\|WP_188801252.1 | Dyella caseinilytica | Gammaproteobacteria | Xanthomonadales | Rhodanobacteraceae |
| gi\|WP_184670675.1 | Rhodanobacter sp. A1T4 | Gammaproteobacteria | Xanthomonadales | Rhodanobacteraceae |
| gi\|WP_130618046.1 | Dyella sp. DHC06 | Gammaproteobacteria | Xanthomonadales | Rhodanobacteraceae |
| gi\|WP_157971311.1 | Dyella sp. C9 | Gammaproteobacteria | Xanthomonadales | Rhodanobacteraceae |
| gi\|KAF1006305.1 | Luteibacter sp. | Gammaproteobacteria | Xanthomonadales | Rhodanobacteraceae |

| | | | | |
|---|---|---|---|---|
| gi\|WP_188793832.1 | Dyella nitratireducens | Gammaproteobacteria | Xanthomonadales | Rhodanobacteraceae |
| gi\|WP_111982642.1 | Dyella jiangningensis | Gammaproteobacteria | Xanthomonadales | Rhodanobacteraceae |
| gi\|SHL61964.1 | Rhodanobacter sp. OK091 | Gammaproteobacteria | Xanthomonadales | Rhodanobacteraceae |
| gi\|WP_183421918.1 | Luteibacter sp. Sphag1AF | Gammaproteobacteria | Xanthomonadales | Rhodanobacteraceae |
| gi\|WP_158605339.1 | Dyella sp. YR388 | Gammaproteobacteria | Xanthomonadales | Rhodanobacteraceae |
| gi\|WP_144911786.1 | Luteibacter yeojuensis | Gammaproteobacteria | Xanthomonadales | Rhodanobacteraceae |
| gi\|WP_137915201.1 | Rudaea sp. 3F27F6 | Gammaproteobacteria | Xanthomonadales | Rhodanobacteraceae |
| gi\|WP_068097298.1 | unclassified Rhodanobacter | Gammaproteobacteria | Xanthomonadales | Rhodanobacteraceae |
| gi\|WP_161970926.1 | Aerosticca soli | Gammaproteobacteria | Xanthomonadales | Rhodanobacteraceae |
| gi\|WP_187056955.1 | Dyella sp. G9 | Gammaproteobacteria | Xanthomonadales | Rhodanobacteraceae |
| gi\|AHX11976.1 | Dyella jiangningensis | Gammaproteobacteria | Xanthomonadales | Rhodanobacteraceae |
| gi\|WP_184506145.1 | Rhodanobacter sp. ANJX3 | Gammaproteobacteria | Xanthomonadales | Rhodanobacteraceae |
| gi\|WP_177257429.1 | Luteibacter sp. UNCMF366Tsu5.1 | Gammaproteobacteria | Xanthomonadales | Rhodanobacteraceae |
| gi\|WP_157956558.1 | Dyella sp. C11 | Gammaproteobacteria | Xanthomonadales | Rhodanobacteraceae |
| gi\|QDE37857.1 | Luteibacter pinisoli | Gammaproteobacteria | Xanthomonadales | Rhodanobacteraceae |
| gi\|WP_157971310.1 | Dyella sp. C9 | Gammaproteobacteria | Xanthomonadales | Rhodanobacteraceae |
| gi\|WP_143525755.1 | Rhodanobacter sp. C05 | Gammaproteobacteria | Xanthomonadales | Rhodanobacteraceae |
| gi\|WP_082879341.1 | Luteibacter rhizovicinus | Gammaproteobacteria | Xanthomonadales | Rhodanobacteraceae |
| gi\|WP_166946525.1 | Luteibacter anthropi | Gammaproteobacteria | Xanthomonadales | Rhodanobacteraceae |
| gi\|WP_188798187.1 | Dyella caseinilytica | Gammaproteobacteria | Xanthomonadales | Rhodanobacteraceae |
| gi\|TAM58479.1 | Rhodanobacter sp. | Gammaproteobacteria | Xanthomonadales | Rhodanobacteraceae |
| gi\|WP_147281721.1 | Dyella solisilvae | Gammaproteobacteria | Xanthomonadales | Rhodanobacteraceae |
| gi\|WP_175483735.1 | Frateuria terrea | Gammaproteobacteria | Xanthomonadales | Rhodanobacteraceae |
| gi\|WP_132142697.1 | Luteibacter rhizovicinus | Gammaproteobacteria | Xanthomonadales | Rhodanobacteraceae |
| gi\|WP_158241376.1 | Dyella sp. AD56 | Gammaproteobacteria | Xanthomonadales | Rhodanobacteraceae |
| gi\|WP_157956557.1 | Dyella sp. C11 | Gammaproteobacteria | Xanthomonadales | Rhodanobacteraceae |
| gi\|WP_167257077.1 | unclassified Dyella | Gammaproteobacteria | Xanthomonadales | Rhodanobacteraceae |
| gi\|TAL86195.1 | Rhodanobacter sp. | Gammaproteobacteria | Xanthomonadales | Rhodanobacteraceae |
| gi\|WP_158241388.1 | Dyella sp. AD56 | Gammaproteobacteria | Xanthomonadales | Rhodanobacteraceae |
| gi\|WP_184417519.1 | Rhodanobacter sp. MP7CTX1 | Gammaproteobacteria | Xanthomonadales | Rhodanobacteraceae |
| gi\|WP_183421984.1 | Luteibacter sp. Sphag1AF | Gammaproteobacteria | Xanthomonadales | Rhodanobacteraceae |
| gi\|WP_131994044.1 | Dokdonella fugitiva | Gammaproteobacteria | Xanthomonadales | Rhodanobacteraceae |
| gi\|TCI10265.1 | Dyella soli | Gammaproteobacteria | Xanthomonadales | Rhodanobacteraceae |
| gi\|WP_157971436.1 | Dyella sp. C9 | Gammaproteobacteria | Xanthomonadales | Rhodanobacteraceae |
| gi\|WP_166700083.1 | Luteibacter yeojuensis | Gammaproteobacteria | Xanthomonadales | Rhodanobacteraceae |
| gi\|WP_019467459.1 | Dyella japonica | Gammaproteobacteria | Xanthomonadales | Rhodanobacteraceae |
| gi\|WP_052395100.1 | Oleiagrimonas soli | Gammaproteobacteria | Xanthomonadales | Rhodanobacteraceae |
| gi\|AHX16294.1 | Dyella jiangningensis | Gammaproteobacteria | Xanthomonadales | Rhodanobacteraceae |
| gi\|WP_139351573.1 | Rhodanobacter sp. C06 | Gammaproteobacteria | Xanthomonadales | Rhodanobacteraceae |
| gi\|WP_114241570.1 | Dyella sp. C9 | Gammaproteobacteria | Xanthomonadales | Rhodanobacteraceae |
| gi\|WP_036138520.1 | Luteibacter sp. 9135 | Gammaproteobacteria | Xanthomonadales | Rhodanobacteraceae |
| gi\|WP_131412637.1 | Dyella soli | Gammaproteobacteria | Xanthomonadales | Rhodanobacteraceae |
| gi\|KAF1004467.1 | Luteibacter sp. | Gammaproteobacteria | Xanthomonadales | Rhodanobacteraceae |

| | | | | |
|---|---|---|---|---|
| gi\|WP_192557081.1 | Dyella sp. 7MK23 | Gammaproteobacteria | Xanthomonadales | Rhodanobacteraceae |
| gi\|WP_074547623.1 | Dyella sp. AtDHG13 | Gammaproteobacteria | Xanthomonadales | Rhodanobacteraceae |
| gi\|OZB58347.1 | Xanthomonadales bacterium 15-68-25 | Gammaproteobacteria | Xanthomonadales | unclassified Xanthomon |
| gi\|WP_162204415.1 | Pseudoxanthomonas suwonensis | Gammaproteobacteria | Xanthomonadales | Xanthomonadaceae |
| gi\|WP_190280147.1 | Thermomonas sp. XSG | Gammaproteobacteria | Xanthomonadales | Xanthomonadaceae |
| gi\|WP_192309408.1 | Pseudoxanthomonas sp. PXM02 | Gammaproteobacteria | Xanthomonadales | Xanthomonadaceae |
| gi\|ASR42995.1 | Xanthomonas citri pv. mangiferaeindicae | Gammaproteobacteria | Xanthomonadales | Xanthomonadaceae |
| gi\|WP_192197253.1 | Pseudoxanthomonas sp. PXM04 | Gammaproteobacteria | Xanthomonadales | Xanthomonadaceae |
| gi\|WP_162455695.1 | Pseudoxanthomonas kalamensis | Gammaproteobacteria | Xanthomonadales | Xanthomonadaceae |
| gi\|WP_157074176.1 | Pseudoxanthomonas mexicana | Gammaproteobacteria | Xanthomonadales | Xanthomonadaceae |
| gi\|WP_156383625.1 | Pseudoxanthomonas sp. Root65 | Gammaproteobacteria | Xanthomonadales | Xanthomonadaceae |
| gi\|WP_184410776.1 | Xanthomonas translucens | Gammaproteobacteria | Xanthomonadales | Xanthomonadaceae |
| gi\|WP_187571448.1 | Thermomonas brevis | Gammaproteobacteria | Xanthomonadales | Xanthomonadaceae |
| gi\|RZA34394.1 | Xanthomonadaceae bacterium | Gammaproteobacteria | Xanthomonadales | Xanthomonadaceae |
| gi\|KQZ63601.1 | Lysobacter sp. Root559 | Gammaproteobacteria | Xanthomonadales | Xanthomonadaceae |
| gi\|WP_130523277.1 | unclassified Pseudoxanthomonas | Gammaproteobacteria | Xanthomonadales | Xanthomonadaceae |
| gi\|WP_162310250.1 | Pseudoxanthomonas broegbernensis | Gammaproteobacteria | Xanthomonadales | Xanthomonadaceae |
| gi\|WP_114959790.1 | Thermomonas haemolytica | Gammaproteobacteria | Xanthomonadales | Xanthomonadaceae |
| gi\|WP_169706862.1 | Xanthomonas campestris | Gammaproteobacteria | Xanthomonadales | Xanthomonadaceae |
| gi\|WP_139187963.1 | Pseudoxanthomonas sp. CF385 | Gammaproteobacteria | Xanthomonadales | Xanthomonadaceae |
| gi\|WP_122230260.1 | Pseudoxanthomonas spadix | Gammaproteobacteria | Xanthomonadales | Xanthomonadaceae |
| gi\|WP_079722955.1 | Pseudoxanthomonas indica | Gammaproteobacteria | Xanthomonadales | Xanthomonadaceae |
| gi\|RYD15446.1 | Xanthomonadaceae bacterium | Gammaproteobacteria | Xanthomonadales | Xanthomonadaceae |
| gi\|KAF1692036.1 | Pseudoxanthomonas jiangsuensis | Gammaproteobacteria | Xanthomonadales | Xanthomonadaceae |
| gi\|WP_065470281.1 | Xanthomonas bromi | Gammaproteobacteria | Xanthomonadales | Xanthomonadaceae |
| gi\|GGF94307.1 | Arenimonas maotaiensis | Gammaproteobacteria | Xanthomonadales | Xanthomonadaceae |
| gi\|WP_111267339.1 | Lysobacter maris | Gammaproteobacteria | Xanthomonadales | Xanthomonadaceae |
| gi\|WP_162314935.1 | Pseudoxanthomonas yeongjuensis | Gammaproteobacteria | Xanthomonadales | Xanthomonadaceae |
| gi\|WP_064507789.1 | Xanthomonas floridensis | Gammaproteobacteria | Xanthomonadales | Xanthomonadaceae |
| gi\|WP_039955939.1 | Xanthomonas translucens | Gammaproteobacteria | Xanthomonadales | Xanthomonadaceae |
| gi\|WP_082132394.1 | Luteimonas sp. FCS-9 | Gammaproteobacteria | Xanthomonadales | Xanthomonadaceae |
| gi\|WP_043907954.1 | Xanthomonas | Gammaproteobacteria | Xanthomonadales | Xanthomonadaceae |
| gi\|WP_183644530.1 | unclassified Pseudoxanthomonas | Gammaproteobacteria | Xanthomonadales | Xanthomonadaceae |
| gi\|WP_056880582.1 | Pseudoxanthomonas sp. Root630 | Gammaproteobacteria | Xanthomonadales | Xanthomonadaceae |
| gi\|ELQ12144.1 | Xanthomonas translucens DAR61454 | Gammaproteobacteria | Xanthomonadales | Xanthomonadaceae |
| gi\|WP_019398200.1 | unclassified Pseudoxanthomonas | Gammaproteobacteria | Xanthomonadales | Xanthomonadaceae |
| gi\|WP_189447070.1 | Lysobacter xinjiangensis | Gammaproteobacteria | Xanthomonadales | Xanthomonadaceae |
| gi\|WP_115858365.1 | Lysobacter silvisoli | Gammaproteobacteria | Xanthomonadales | Xanthomonadaceae |
| gi\|WP_104586694.1 | Xanthomonas melonis | Gammaproteobacteria | Xanthomonadales | Xanthomonadaceae |
| gi\|WP_137267711.1 | Luteimonas gilva | Gammaproteobacteria | Xanthomonadales | Xanthomonadaceae |
| gi\|SBV36673.1 | uncultured Stenotrophomonas sp. | Gammaproteobacteria | Xanthomonadales | Xanthomonadaceae |
| gi\|WP_184646616.1 | Xanthomonas arboricola | Gammaproteobacteria | Xanthomonadales | Xanthomonadaceae |
| gi\|WP_152239870.1 | Xanthomonas sp. LMG 12461 | Gammaproteobacteria | Xanthomonadales | Xanthomonadaceae |

| | | | | |
|---|---|---|---|---|
| gi\|WP_184645421.1 | Xanthomonas arboricola | Gammaproteobacteria | Xanthomonadales | Xanthomonadaceae |
| gi\|RMH90922.1 | Lysobacter pythonis | Gammaproteobacteria | Xanthomonadales | Xanthomonadaceae |
| gi\|WP_166636837.1 | Lysobacter terrigena | Gammaproteobacteria | Xanthomonadales | Xanthomonadaceae |
| gi\|WP_045728332.1 | Xanthomonas sp. GPE 39 | Gammaproteobacteria | Xanthomonadales | Xanthomonadaceae |
| gi\|WP_054658204.1 | Stenotrophomonas pictorum | Gammaproteobacteria | Xanthomonadales | Xanthomonadaceae |
| gi\|VXC18703.1 | Luteimonas sp. 9C | Gammaproteobacteria | Xanthomonadales | Xanthomonadaceae |
| gi\|WP_158984567.1 | Lysobacter panacisoli | Gammaproteobacteria | Xanthomonadales | Xanthomonadaceae |
| gi\|WP_167708968.1 | Xanthomonas arboricola | Gammaproteobacteria | Xanthomonadales | Xanthomonadaceae |
| gi\|WP_164081923.1 | Stenotrophomonas maltophilia | Gammaproteobacteria | Xanthomonadales | Xanthomonadaceae |
| gi\|WP_194930138.1 | Lysobacter niastensis | Gammaproteobacteria | Xanthomonadales | Xanthomonadaceae |
| gi\|WP_055821220.1 | Xanthomonas sp. Leaf131 | Gammaproteobacteria | Xanthomonadales | Xanthomonadaceae |
| gi\|WP_144900426.1 | Luteimonas cucumeris | Gammaproteobacteria | Xanthomonadales | Xanthomonadaceae |
| gi\|WP_162125461.1 | Pseudoxanthomonas taiwanensis | Gammaproteobacteria | Xanthomonadales | Xanthomonadaceae |
| gi\|OAG66166.1 | Xanthomonas floridensis | Gammaproteobacteria | Xanthomonadales | Xanthomonadaceae |
| gi\|WP_165782402.1 | Lysobacter silvestris | Gammaproteobacteria | Xanthomonadales | Xanthomonadaceae |
| gi\|WP_003477195.1 | Xanthomonas translucens | Gammaproteobacteria | Xanthomonadales | Xanthomonadaceae |
| gi\|WP_141517630.1 | Lysobacter aestuarii | Gammaproteobacteria | Xanthomonadales | Xanthomonadaceae |
| gi\|WP_166294242.1 | Lysobacter sp. HDW10 | Gammaproteobacteria | Xanthomonadales | Xanthomonadaceae |
| gi\|WP_160954828.1 | Xanthomonas | Gammaproteobacteria | Xanthomonadales | Xanthomonadaceae |
| gi\|WP_142741729.1 | Xanthomonas translucens | Gammaproteobacteria | Xanthomonadales | Xanthomonadaceae |
| gi\|WP_158734000.1 | Lysobacter prati | Gammaproteobacteria | Xanthomonadales | Xanthomonadaceae |
| gi\|QHQ29235.1 | Xanthomonas albilineans | Gammaproteobacteria | Xanthomonadales | Xanthomonadaceae |
| gi\|SBV37381.1 | uncultured Stenotrophomonas sp. | Gammaproteobacteria | Xanthomonadales | Xanthomonadaceae |
| gi\|WP_189375872.1 | Thermomonas carbonis | Gammaproteobacteria | Xanthomonadales | Xanthomonadaceae |
| gi\|GHH47782.1 | [Pseudomonas] boreopolis | Gammaproteobacteria | Xanthomonadales | Xanthomonadaceae |
| gi\|WP_187570655.1 | Thermomonas brevis | Gammaproteobacteria | Xanthomonadales | Xanthomonadaceae |
| gi\|TDK30819.1 | Luteimonas terrae | Gammaproteobacteria | Xanthomonadales | Xanthomonadaceae |
| gi\|WP_166056862.1 | Thermomonas sp. HDW16 | Gammaproteobacteria | Xanthomonadales | Xanthomonadaceae |
| gi\|WP_065469730.1 | Xanthomonas bromi | Gammaproteobacteria | Xanthomonadales | Xanthomonadaceae |
| gi\|RFP60300.1 | Lysobacter sp. WF-2 | Gammaproteobacteria | Xanthomonadales | Xanthomonadaceae |
| gi\|WP_108757030.1 | Stenotrophomonas sp. YAU14A_MKIMI4_1 | Gammaproteobacteria | Xanthomonadales | Xanthomonadaceae |
| gi\|WP_082594976.1 | Stenotrophomonas | Gammaproteobacteria | Xanthomonadales | Xanthomonadaceae |
| gi\|PAK92049.1 | Stenotrophomonas rhizophila | Gammaproteobacteria | Xanthomonadales | Xanthomonadaceae |
| gi\|PJK10582.1 | Xanthomonadaceae bacterium NML95-0200 | Gammaproteobacteria | Xanthomonadales | Xanthomonadaceae |
| gi\|ASR42746.1 | Xanthomonas citri pv. mangiferaeindicae | Gammaproteobacteria | Xanthomonadales | Xanthomonadaceae |
| gi\|KGQ20270.1 | Lysobacter dokdonensis DS-58 | Gammaproteobacteria | Xanthomonadales | Xanthomonadaceae |
| gi\|WP_146313050.1 | Luteimonas wenzhouensis | Gammaproteobacteria | Xanthomonadales | Xanthomonadaceae |
| gi\|WP_082925777.1 | Xanthomonas nasturtii | Gammaproteobacteria | Xanthomonadales | Xanthomonadaceae |
| gi\|WP_078059571.1 | Xanthomonas massiliensis | Gammaproteobacteria | Xanthomonadales | Xanthomonadaceae |
| gi\|WP_129136654.1 | Luteimonas sp. YGD11-2 | Gammaproteobacteria | Xanthomonadales | Xanthomonadaceae |
| gi\|WP_064510318.1 | Xanthomonas floridensis | Gammaproteobacteria | Xanthomonadales | Xanthomonadaceae |
| gi\|WP_185817396.1 | Xanthomonas theicola | Gammaproteobacteria | Xanthomonadales | Xanthomonadaceae |
| gi\|WP_012437146.1 | Xanthomonas campestris | Gammaproteobacteria | Xanthomonadales | Xanthomonadaceae |

| | | | | |
|---|---|---|---|---|
| gi\|WP_130318714.1 | Stenotrophomonas sp. BK441 | Gammaproteobacteria | Xanthomonadales | Xanthomonadaceae |
| gi\|WP_072755682.1 | Thermomonas hydrothermalis | Gammaproteobacteria | Xanthomonadales | Xanthomonadaceae |
| gi\|WP_141483191.1 | Lysobacter maris | Gammaproteobacteria | Xanthomonadales | Xanthomonadaceae |
| gi\|PBJ84078.1 | Xanthomonadaceae bacterium NML93-0399 | Gammaproteobacteria | Xanthomonadales | Xanthomonadaceae |
| gi\|KRG40176.1 | Stenotrophomonas panacihumi | Gammaproteobacteria | Xanthomonadales | Xanthomonadaceae |
| gi\|WP_088060991.1 | Xanthomonas fragariae | Gammaproteobacteria | Xanthomonadales | Xanthomonadaceae |
| gi\|BBO50577.1 | Stenotrophomonas maltophilia | Gammaproteobacteria | Xanthomonadales | Xanthomonadaceae |
| gi\|WP_072755493.1 | Thermomonas hydrothermalis | Gammaproteobacteria | Xanthomonadales | Xanthomonadaceae |
| gi\|WP_115514668.1 | Xanthomonas | Gammaproteobacteria | Xanthomonadales | Xanthomonadaceae |
| gi\|WP_157029055.1 | Lysobacter soli | Gammaproteobacteria | Xanthomonadales | Xanthomonadaceae |
| gi\|WP_011269546.1 | Xanthomonas campestris | Gammaproteobacteria | Xanthomonadales | Xanthomonadaceae |
| gi\|PJK14921.1 | Xanthomonadaceae bacterium NML07-0707 | Gammaproteobacteria | Xanthomonadales | Xanthomonadaceae |
| gi\|TXH68398.1 | Xanthomonadaceae bacterium | Gammaproteobacteria | Xanthomonadales | Xanthomonadaceae |
| gi\|WP_144890245.1 | Luteimonas granuli | Gammaproteobacteria | Xanthomonadales | Xanthomonadaceae |
| gi\|WP_132999328.1 | Luteimonas arsenica | Gammaproteobacteria | Xanthomonadales | Xanthomonadaceae |
| gi\|WP_099820855.1 | Stenotrophomonas sp. LMG 10879 | Gammaproteobacteria | Xanthomonadales | Xanthomonadaceae |
| gi\|WP_043957317.1 | Lysobacter sp. A03 | Gammaproteobacteria | Xanthomonadales | Xanthomonadaceae |
| gi\|WP_158601676.1 | Lysobacter pythonis | Gammaproteobacteria | Xanthomonadales | Xanthomonadaceae |
| gi\|KHL58393.1 | Xanthomonas cannabis pv. cannabis | Gammaproteobacteria | Xanthomonadales | Xanthomonadaceae |
| gi\|WP_027081965.1 | Lysobacter sp. URHA0019 | Gammaproteobacteria | Xanthomonadales | Xanthomonadaceae |
| gi\|WP_192287484.1 | Stenotrophomonas sp. STM01 | Gammaproteobacteria | Xanthomonadales | Xanthomonadaceae |

### Table S2. Plasmids used in this study

| | Plasmid name | Description |
|---|---|---|
| | pSS1129 | Suicide plasmid for homologous recombination in *Bordetella* |
| | pQC2123 | Suicide plasmid carrying *lacZ* for chromosomal transcriptional fusions in *B. pertussis* |
| | pRM1 | Suicide plasmid for genetic insertion in the *ure* locus of *Bordetella pertussis* |
| Allelic exchange | pSS1129-Δ*bp2923* | Plasmid carrying the regions flanking *bp2923* for its deletion |
| | pSS1129-Δ*bfrG* | Plasmid carrying the regions flanking *bfrG* for its deletion |
| | pSS1129-*bp2923*-OCU | Plasmid carrying *bp2923*-OCU and its flanking regions for insertion in BP*Δ2923* by homologous recombination |
| | pRM1-*bp2923* | Plasmid carrying *bp2923* flanked by *ure* locus regions for complementation in *trans* |
| Gene KO | pFUS2-efp | Suicide plasmid carrying a 316-bp fragment of *efp* used to knock out this gene |
| Chromosomal translational *lacZ* fusions | pQC2123-*bp2923-lacZ* | Translational fusion between the first 10 codons of *bp2923* and *lacZ*; contains a 631-bp EcoRI-XhoI fragment in pQC2123 for homologous recombination |
| | pQC2123-*bfrG-lacZ* | Translational fusion between the first 10 codons of *bfrG* and *lacZ*; contains a 1225-bp EcoRI-XhoI fragment in pQC2123 for homologous recombination |
| | pQC2123-*bp2921-lacZ* | Translational fusion between the first 10 codons of bp2921 and *lacZ*; contains a 635-bp EcoRI-XhoI fragment in pQC2123, for homologous recombination |
| | pUC57-*cruR* wt | Plasmid carrying a 797-bp EcoRI-XhoI fragment encompassing part of *bp2924,* the intergenic region between it and *cruR, cruR* and the 10 first codons of *bfrG,* used to introduce mutations in *bp2923* |
| | pQC2123-*bp2923*$P_{120}$A+$P_{121}$A-*lacZ* | Translational fusions between the first 10 codons of *bfrG and lacZ,* with the indicated mutations in *bp2923; the* EcoRI-XhoI fragments are from the corresponding pUC57-*cruR* plasmids. The fragment comprising *bp2924* and the intergenic region between it and *bp2923* drives homologous recombination in the chromosome of BP*Δ2923* |
| | pQC2123-*bp2923*$P_{141}$A+$P_{142}$A-*lacZ* | |
| | pQC2123-*bp2923*$C_{90}$S+$C_{93}$S-*lacZ* | |
| | pQC2123-*bp2923*$Y_{50}$STOP-*lacZ* | |
| | pQC2123-*bp2923*$R_{139}$A+$A_{140}$S-*lacZ* | |
| | pQC2123-*bp2923*$C_{51}$S-*lacZ* | |
| | pQC2123-*bp2923*$FS_{117-123}$-*lacZ* | |
| | pQC2123-*bp2923*$FS_{5-44}$-*lacZ* | |
| | pQC2123-*bp2923*$W_{133}$STOP-*lacZ* | |
| | pQC2123-*bp2923*-2-*lacZ* | |
| | pQC2123-*bp2923*-1-*lacZ* | |
| | pQC2123-*bp2923*+1-*lacZ* | |
| | pQC2123-*bp2923*+2-*lacZ* | |
| | pQC2123-*bp2923*+5-*lacZ* | |
| | pQC2123-*bp2923*-OCU-*lacZ* | |

### Table S3. Oligonucleotides used in this study

| | | Sequences (5'-3') |
|---|---|---|
| Deletions | 2923 EcoRI UP | ATGAATTCACGCCGTAGCATTTCTCCATGT |
| | 2923 XbaI LO | ATTCTAGAGCCGGTCTGGATCATGGT |
| | 2923 XbaI UP | TATCTAGATGAGCGCGGCAGCCTTCC |
| | 2923 HindIII LO | ATAAGCTTCTCGCGGTTGATGACGGTCAC |
| | bfrg UP | GGATCCCAGGAGCAATCGCTTCCCGT |
| | bfrG XbaI LO | TGTAGCGGTAGGTCTCGAGGT |
| | bfrG XbaI UP | ACCTCGAGACCTACCGCTACA |
| | bfrG LO | TGAAGCTTCGCCTGCTCCCTCAGAAGC |
| *bp2923* complementation | 2923comp UP | TAGGATCCCGCGTCAGTCACGGGCAAGG |
| | 2923comp LO | TATCTAGAAAGGCTGCCGCGCTCAGG |
| Gene knock out | efp UP | ATAAGCTTGACCCGCTCGTCGTCCAGA |
| | Efp LO | TAGGATCCGCCGTCGTAGAAGACCACCT |
| 5'RACE | bfrG | GCCAGCGCCAGTCCGACC |
| | 2923-OCU | CTCCGTGCAGTAGTCGACCGAT |
| | 2923-OCU nested | ATGGTCCGAGATTATAAACAGCG |
| *lacZ* translational fusions | Fus-UP | ATGAATTCTCAAGACCCGGTC |
| | 2923Fus LO | ATCTCGAGCAGGCGACGAGAGCGCGA |
| | bfrGFus LO | ATCTCGAGGGCGTAGTAAAGGGAAGTCTGT |
| | 2921Fus UP | TAGAATTCACCACCGGCACCCAGGTCGAG |
| | 2921Fus LO | TACTCGAGGGCGCGCAAGACTAGGGTCGT |
| RT-PCR | RT UP | ACCTGGAACATTGCCCGTTCT |
| | RT LO | CACGACCACCGGCGCGAGTTG |
| qRT-PCR | *bp3416* UP | GTTCCTGGAAGTGCTGCTG |
| | *bp3416* LO | GATGTCGAAGGCATTCTGG |
| | *bp2923* UP | CCGCTCGCATGGATCGCCTT |
| | *bp2923* LO | CGTCCCCTGCTCGGTGCAAT |
| | *bfrG* UP | CTCGACACGCAGGAAATCGC |
| | *bfrG* LO | CAACTGGCTGACCTGCGAAC |
| | *bp2921* UP | CAAGCGCTGGCTGTATTTG |
| | *bp2921* LO | CAGATTGGGATAGCCGACATAC |
| Mutagenesis | G5+1 UP | GGACCATGATCCAGACCAGGCTCGCGCTCTCGTCG |
| | G5+1 LO | CGACGAGAGCGCGAGCCTGGTCTGGATCATGGTCC |
| | L44-1 UP | CAACGGCATGCGCCGCTGCGCTGTCGGTCGACTA |
| | L44-1 LO | TAGTCGACCGACAGCGCAGCGGCGCATGCCGTTG |
| | C51S UP | CGCTGTCGGTCGACTATAGCACCGAGCAGGGGACG |
| | C51S LO | CGTCCCCTGCTCGGTGCTATAGTCGACCGACAGCG |
| | 116+1 UP | CTTGCCGGCGCGCGCCGATTTCCTATCC |
| | 116+1 LO | GGATAGGAAATCGGCGCGCGCCGGCAAG |
| | 124-1 UP | CGATTTCCTATCCTCCTCTCTTTACGTTGCCCG |
| | 124-1 LO | CGGGCAACGTAAAGAGAGGAGGATAGGAAATCG |
| | W133* UP | CGTCGCAGCATGCCTGAACCGGCGCGCAGCCGC |
| | W133* LO | GCGGCTGCGCGCCGGTTCAGGCATGCTGCGACG |
| In vitro transcription | FWT1b | TTATCAAAAAGAGTATTGACTTAAAGTCTAACCTATAGGATACTTACAGCCAGATTGTGCGCGAGCGTGGG |
| | FWT2b | TTATCAAAAAGAGTATTGACTTAAAGTCTAACCTATAGGATACTTACAGCCAGGGCGACCTGGCCC |
| | FWT3b | TTATCAAAAAGAGTATTGACTTAAAGTCTAACCTATAGGATACTTACAGCCAGCATGCCTGGACCGGCGCGC |
| | FWT4-6 | TTATCAAAAAGAGTATTGACTTAAAGTCTAACCTATAGGATACTTACAGCCAGGGCGACCTGGCCC |
| | REVT1-3 | CGAGGGCGTAGTAAAGGGAAGTCTG |
| | REVT4 | CGGCGGGCATGTCGATGACGGGC |

| | | |
|---|---|---|
| | REVT5 | GGCCCGGCCGCAAGGCGGACGGGC |
| | REVT6 | GCACGGCTGGCCGTGGGCGCTGG |
